# Cellular and Spatial Drivers of Unresolved Injury and Functional Decline in the Human Kidney

**DOI:** 10.1101/2025.09.26.678707

**Authors:** Blue B. Lake, Ricardo Melo Ferreira, Jens Hansen, Rajasree Menon, Jeannine Basta, Heather Thiessen Philbrook, Stephanie Reinert, Robin Fallegger, Asmita K. Lagwankar, Xi Chen, Soumya Maity, Katerina V. Djambazova, Brittney L. Gorman, Nicholas Lucarelli, Debora L. Gisch, Insa M. Schmidt, Viji Nair, Fadhl Alakwaa, Eirini Kefaloyianni, Bo Zhang, Amanda L. Knoten, Madhurima Kaushal, Edgar A. Otto, Melissa A. Farrow, Dinh Diep, Dusan Velickovic, Angela R. Sabo, Elijah Cole, Ian Tamayo, Jovan Tanevski, Kimberly Y. Conklin, Rachel S. G. Sealfon, Yongqun He, Michelle Brennan, Lynn Robbins, Ying-Hua Cheng, Markus Bitzer, Aditya Surapaneni, Steven Menez, Peter V. Kharchenko, Charles E. Alpers, Ulysses G. J. Balis, Laura Barisoni, Ian H. de Boer, Dawit Demeke, Agnes B. Fogo, Joel M. Henderson, Leal Herlitz, Gilbert W. Moeckel, Parmjeet S. Randhawa, Avi Z. Rosenberg, Jennifer A. Schaub, Suman Setty, Frank C. Brosius, Maria L. Caramori, Steven G. Coca, Robert S. Figenshau, Eric H. Kim, Krzysztof Kiryluk, James P. Lash, R. Tyler Miller, John F. O’Toole, Paul M. Palevsky, Eugene P. Rhee, Ana C. Ricardo, Sylvia E. Rosas, Prabir Roy-Chaudhury, Minnie M. Sarwal, John R. Sedor, Robert D. Toto, Aydın Türkmen, Sushrut S. Waikar, James C. Williams, F P. Wilson, E. Steve Woodle, Evan Z. Macosko, Julio Saez-Rodriguez, Pierre C Dagher, Morgan E. Grams, Petter Bjornstad, Tarek M. El-Achkar, Olga G. Troyanskaya, Nikole Bonevich, Pinaki Sarder, Sanjeev Kumar, Christopher R. Anderton, Jeffrey M. Spraggins, Kumar Sharma, Michael Rauchman, Jonathan Himmelfarb, Joseph P. Gaut, Kidney Precision Medicine Project, Kun Zhang, Ravi Iyengar, Matthias Kretzler, Jeffrey B. Hodgin, Chirag R. Parikh, Michael T. Eadon, Sanjay Jain, HuBMAP consortium and Kidney Precision Medicine Project

**Affiliations:** Altos Labs San Diego Institute of Science, San Diego, CA, USA; Department of Medicine, Indiana University School of Medicine, Indianapolis, IN 46202, USA; Department of Pharmacological Sciences, Icahn School of Medicine at Mount Sinai, New York, NY 10029, USA; Mount Sinai Institute for Systems Biomedicine, Icahn School of Medicine at Mount Sinai, New York, NY 10029, USA; Departmentof Computational Medicine and Bioinformatics, University of Michigan, Ann Arbor, MI 48109, USA; Division of Nephrology, Department of Medicine, Washington University School of Medicine, St. Louis, MO 63110, USA; Division of Nephrology, Johns Hopkins School of Medicine, Baltimore, MD 21287, USA; Institute for Computational Biomedicine, Heidelberg University and Heidelberg University Hospital, Heidelberg, Germany; Center for Computational Biology, Flatiron Institute, Simons Foundation, New York, NY, USA; Lewis-Sigler Institute of Integrative Genomics, Princeton University, Princeton, NJ, USA; Center for Precision Medicine, The University of Texas at San Antonio, San Antonio, TX, USA; Department of Cell and Developmental Biology and Mass Spectrometry Research Center, Vanderbilt University School of Medicine, Nashville, TN 37232, USA; Pacific Northwest National Laboratory, Richland, WA, USA; Department of Medicine-Nephrology & Intelligent Critical Care Center, University of Florida, Gainesville, FL, USA; Department of Medicine, Indiana University School of Medicine, Indianapolis, IN, 46202, USA; Department of Medicine, Boston University Chobanian & Avedisian School of Medicine, Boston, MA, USA; Department of Internal Medicine, Division of Nephrology, University of Michigan, Ann Arbor, MI 48109, USA; Center for Computational Medicine and Bioinformatics, University of Michigan, Ann Arbor, MI 48109, USA; Department of Medicine, Washington University School of Medicine, St. Louis, MO 63110, USA; Institute of Informatics, Department of Medicine, Washington University School of Medicine, St. Louis, MO 63110, USA; Center for Precision Medicine, The University of Texas Health San Antonio, San Antonio, TX, USA; Division of Nephrology, Department of Medicine, The University of Texas Health San Antonio, San Antonio, TX, USA; Unit for Laboratory Animal Medicine, Department of Learning Health Sciences, University of Michigan, Ann Arbor, MI 48109, USA; Department of Biochemistry and Molecular Biology, Saint Louis University School of Medicine, St. Louis, MO, 63103, USA; St. Louis Veteran Affairs Medical Center, St. Louis, MO, 63106, USA; Department of Medicine, New York University, New York, USA; Department of Laboratory Medicine and Pathology, University of Washington, Seattle, Washington, USA; Department of Pathology, University of Michigan, Ann Arbor, MI 48109, USA; Department of Pathology, Division of AI and Computational Pathology, Duke University, Durham, NC, USA; Department of Medicine, Division of Nephrology, Duke University, Durham, NC, USA; Kidney Research Institute, Division of Nephrology, 325 Ninth Avenue, Box 359606, Seattle, WA 98104, USA; Department of Pathology, Microbiology and Immunology, Vanderbilt University Medical Center, Nashville, TN 37232 USA; Department of Pathology and Laboratory Medicine, Boston University Chobanian & Avedisian School of Medicine, Boston, MA, USA; Department of Anatomic Pathology, Cleveland Clinic, Cleveland, OH, USA; Department of Pathology, Yale University, New Haven, CT, USA; Department of Pathology, Thomas E Starzl Transplant Insititute & University of Pittsburgh School of Medicine, Pittsburgh, PA 15213, USA; Department of Pathology, Johns Hopkins University, Baltimore, MD, USA; Department of Pathology, University of Illinois at Chicago, Chicago, IL, USA; Department of Medicine, University of Arizona, Tucson, AZ, USA; Department of Endocrinology, Diabetes and Metabolism, Cleveland Clinic Foundation, Cleveland, OH, USA; Division of Nephrology, Icahn School of Medicine at Mount Sinai, New York, NY, USA; Washington University in Saint Louis, St. Louis, MO, 63103, USA; Department of Surgery, University of Nevada Reno School of Medicine, Reno, NV 89502, USA; Department of Physiology and Cell Biology, University of Nevada Reno School of Medicine, Reno, NV 89502, USA; Department of Medicine, Columbia University, New York, NY 10032, USA; Department of Medicine, University of Illinois Chicago, Chicago, IL, USA; Department of Internal Medicine, University of Texas Southwestern Medical Center, Dallas, TX, USA; Medicine Service, VA North Texas Health Care System, Dallas, TX, USA; Pak Center for Mineral Metabolism and Clinical Research, University of Texas Southwestern Medical Center, Dallas, TX, USA; Lerner Research and Medical Specialties Institutes, Cleveland Clinic, Cleveland, OH 44195, USA; Department of Medicine, University of Pittsburgh School of Medicine, Pittsburgh, PA 15213, USA; Division of Nephrology, Department of Medicine, Massachusetts General Hospital, Boston, MA, USA; Kidney and Hypertension Unit, Joslin Diabetes Center and Harvard Medical School, Boston, MA 02215, USA; Department of Medicine, Division of Nephrology and Hypertension, University of North Carolina School of Medicine, Chapel Hill, NC, USA; Division of Multi-Organ Transplantation, Department of Surgery, University of California, San Francisco, San Francisco, CA, USA; Department of Internal Medicine, UT Southwestern Medical Center, Dallas, TX 75390, USA; Division of Nephrology, Istanbul School of Medicine, Istanbul, Turkey; Section of Nephrology, Boston University School of Medicine and Boston Medical Center, Boston, MA 02118, USA; Department of Anatomy, Cell Biology & Physiology, Indiana University School of Medicine, Indianapolis, IN 46202, USA; Clinical and Translational Research Accelerator, Department of Medicine, Yale University School of Medicine, New Haven, CT 06510, USA; Department of Surgery, University of Cincinnati, Cincinnati, OH, USA; Broad Institute of Harvard and MIT, Cambridge, MA 02142, USA; European Bioinformatics Institute, Wellcome Genome Campus, Hinxton, Cambridgeshire, CB10 1SD, UK; Heidelberg University, Faculty of Medicine, and Heidelberg University Hospital, Institute for Computational Biomedicine (ICB), Heidelberg; University of Washington Medicine Diabetes Institute, University of Washington, Seattle, Washington, USA; Department of Medicine, Division of Nephrology, University of Colorado School of Medicine, Aurora, CO, USA; Department of Computer Science, Princeton University, Princeton, NJ 08544, USA; Lewis-Sigler Institute for Integrative Genomics, Princeton University, Princeton, NJ 08544, USA; Princeton Precision Health, Princeton University, Princeton, NJ 08544, USA; Center for Computational Biology, Flatiron Institute, Simons Foundation, New York, NY 10010, USA; Department of Medicine – Section of Quantitative Health, University of Florida, Gainesville, FL, USA; Department of Medicine, Regenerative Medicine Institute, Cedars Sinai Medical Center, Los Angeles, CA, 90048, USA; Icahn School of Medicine at Mount Sinai, New York, NY 10029, USA; Department of Pathology and Immunology, Washington University School of Medicine, St. Louis, MO 63110, USA; Department of Pediatrics, Washington University School of Medicine, St. Louis, MO 63110, USA; Kidney Translational Research Center, Washington University School of Medicine, St. Louis, MO 63110, USA

## Abstract

Building upon a foundational Human Kidney resource, we present a comprehensive multi-modal atlas that defines spatially resolved versus unresolved repair states and mechanisms in human kidney disease. Homeostatic interactions between injured kidney epithelium and its surrounding milieu determine successful repair outcomes, while pathogenic signaling promotes unresolved inflammation and fibrosis leading to chronic disease. We integrated multiple single-cell and spatial modalities across ∼700 samples from >350 patients (∼250 research biopsies), analyzing ∼1.7 million cells alongside complementary mouse multi-omic profiles spanning acute-to-chronic injury and aging (>300,000 cells) and spatial transcriptomic analysis of >150 human biopsies. This cross-species atlas delineates functional pathways and druggable targets across the nephron and defines gene regulatory networks and chromatin landscapes governing tubular, fibroblast, and immune cell transitions from injury to either recovery or failed repair states. We identified distinct cellular states associated with specific pathological features that show dynamic distributions between acute kidney injury (AKI) and chronic kidney disease (CKD), organized within unique spatial niches that reveal progression mechanisms from early injury to unresolved disease. Gene regulatory analyses prioritized key transcription factor activities (SOX4, SOX9, NFKB1, REL, KLFs) and their target networks establishing disease states and tissue microenvironments. These regulatory programs were directly linked to clinical outcomes, identifying molecular signatures of recovery and secreted biomarkers predictive of AKI-to-CKD progression, providing a key resource for therapeutic development and precision medicine approaches in kidney disease.

## Introduction

The kidneys have critical systemic roles to filter waste products from the blood, maintain body fluid homeostasis, and regulate blood pressure. Given their design and function, they remain highly susceptible to injuries that can arise from toxins, ischemia, hemodynamic changes, or infiltration of immune cells. Following injury, kidney epithelial cells undergo complex transitions to various altered cell states depending on the extent and nature of the injury, including: degenerative or dedifferentiated (d) states associated with elevated stress and cell-death pathways; adaptive (a) states where injured or activated cells may follow a path of recovery or progress to an unresolved state of failed repair (fr); transitional states associated with trans-differentiation; or repair states that re-activate developmental and cell cycle programs to repair or replace lost cells^1^. Within the interstitial space, activation of repair states in stromal cells facilitates extracellular matrix (ECM) remodeling and tissue stabilization, while immune cell recruitment contributes to debris removal and tissue remodeling. Resolution of these states permits successful recovery of the injury niche to a functional or adaptive state. However, a failure to resolve one or more of these states may lead to persistent epithelial dysfunction associated with reduced tissue function, excessive ECM deposition, ongoing inflammation and senescence. Understanding molecular programs governing cell state transitions along resolving versus non-resolving paths of tissue repair is essential for identifying pathogenic processes that drive fibrosis following acute injury, and the cell states that promote chronic disease progression.

Prior human kidney single cell studies, including those within the Kidney Precision Medicine Project (KPMP) ^2^ and the Human BioMolecular Atlas Program (HuBMAP)^3^, have established foundational knowledge of major cell types and altered cell states in the human kidney including the original human kidney atlas or HKAv1^1,4–7^. However, several critical gaps remain: limited spatial resolution of injury niches, incomplete characterization of rare cell states, limited human kidney biopsy samples and insufficient integration across disease states and clinical outcomes. We address these limitations by defining the key regulatory programs in relation to treatment response, disease progression or resolution. To this end, we provide an expansive dataset that enabled defining a comprehensive set of common and rare cell identities, including immune, stromal and endothelial subpopulations and altered states not previously reported for the kidney. We performed extensive orthogonal validation and discovery with four spatial transcriptomic technologies and mechanistic studies. Critically, we captured the full spectrum of the global kidney disease burden, including AKI with and without progression, CKD and tissues without significant pathology (healthy reference tissue – HRT). A parallel multiomic single cell mouse atlas defining the time course of acute and chronic injury states and ageing was constructed and mapped to the human disease trajectories and regulatory networks to better define the evolution of these injury repair states and niches. We deciphered spatial niches that reflect epithelial repair, a set of soluble proteins associated with AKI and CKD progression exemplary of early-mid injury, and transcription factor networks regulating kidney function decline, including *SOX4* which modulates resolution versus unresolved failed repair states.These maps enable development of novel drug targets that may change the course of kidney disease, discovery of soluble markers that predict disease progression, capturing the efficacy and defining the mechanism of new treatments of kidney disease.

## Results

### Building a kidney cell atlas

To expand upon the existing human kidney atlas (HKAv1) ^1^, we studied kidney tissue and data from a population of 353 unique participants in the KPMP and HuBMAP consortia or local sites **(Figure 1a, Supplementary Tables 1-9).** From these, we processed and analyzed 308 single nucleus RNA-seq and multiome (RNA / ATAC, 223 patients), 127 single cell RNA-seq (112 patients), 58 Cut&Run (17 patients), 153 Visium (135 patients), 8 Slide-seq2 (6 patients), 3 CosMx (3 patients), 22 Xenium (21 patients), 3 Codex/MxIF (3 patients) and 27 imaging mass spectrometry (27 patients – spatial metabolomics (9), spatial lipidomics (5), spatial N-glycomics (13)) datasets. This provided a total coverage of 709 datasets (578 total new datasets from HKAv1) across all modalities. Patient samples were carefully assessed for clinical and pathological features in order to categorize HRT, AKI or CKD. Further characterization of disease tissues within KPMP included clinical adjudication of diagnoses and pathology descriptor scoring.

**Figure 1.**
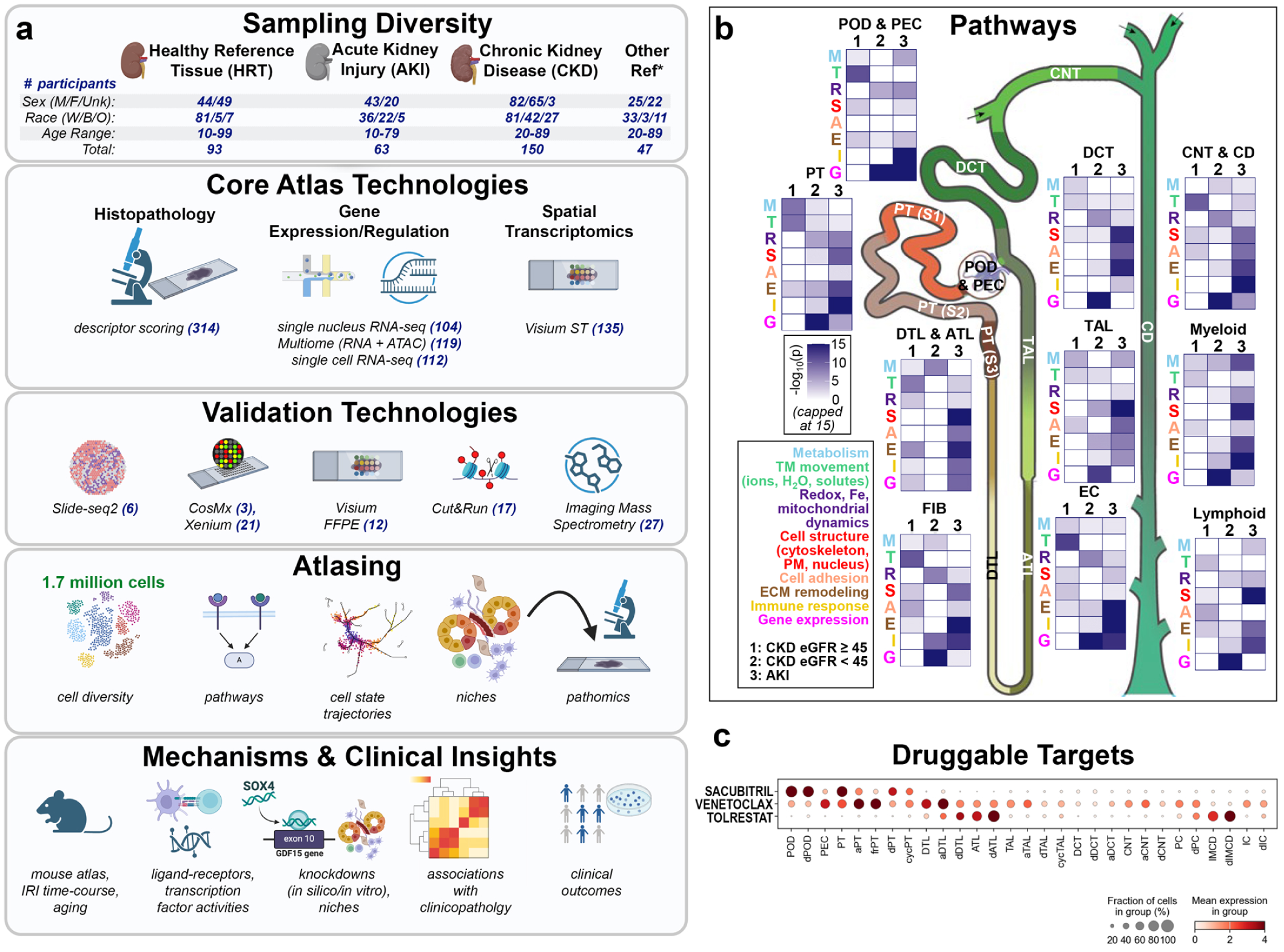
Overview of HKAv2 and impact. **a.** Human kidney samples from healthy reference tissues (HRT), as well as disease tissues associated with AKI and CKD, were processed using one or more assays. The strategy included deep single cell and spatial phenotyping, validations with orthogonal technologies and using analytical methods to delineate spatial contexts of recovery and failed repair. This included pathological staining and assessment, 10X Genomics single nucleus / cell RNA-seq and Multiome (RNA + ATAC), 10X Genomics Visium, Slide-seq2, 10X Xenium, Nanostring CosMx and spatial metabolomics. The atlas demonstrates clinical impact and utility to inform on pathogenetic mechanisms, druggable targets, non-invasive markers that predict clinical outcomes from early injury signatures and the underlying regulatory networks that contribute to AKI progression. Abbreviations: Sex (M = male, F = Female, Unk = unknown); Race (W = white, B = Black, O = other); Age range associated with decade-binned age values per participant. Asterix (other reference tissues including Diabetes Mellitus – Resilient (DM-R) biopsies and reference samples with unknown clinical status. **b.** Pathways predicted for each disease group in each FTU were grouped into 14 whole cell functions, 8 of which are shown. Heatmaps show the sum of the −log10(p-values) of all pathways annotated to the same whole cell function. ATL/DTL: ascending/descending thin limb, CD: collecting duct, CNT: connecting tubule, DCT: distal convoluted tubule, EC: endothelial cell, FIB: fibroblast, PT: proximal tubule, S: segment, TAL: thick ascending limb, POD: podocyte, PEC: parietal epithelial cell. **c.** Map of novel druggable targets specific to cell states. Created in BioRender. Jain, S. (2025) https://BioRender.com/rfy8fos. Source data provided.

To define cell types and states, single nucleus data were first aligned to the HKAv1, then broad cell type categories were independently clustered to identify fine-grained cell subpopulations (see Methods, **ED Figure 1, SD Figure 1, Supplementary Table 10**). Clusters were well integrated across patients and showed variable contributions from different patient conditions associated with their altered state status (**Supplementary Table 11**). This strategy enabled a high level of cell type resolution, identifying 128 distinct cell identities including 60 altered cell states (**ED Figure 1**) having distinct marker gene profiles (**SD Figure 2, Supplementary Table 12**). Cell type annotations incorporated HKAv1 cell type signatures, published kidney and cross-tissue studies^1,4–6,8–17^, clinicopathological associations (**ED Figure 1, SD Figure 1**) and spatial transcriptomic (ST) mapping that included Slide-seq2, CosMx, Visium and Xenium (**ED Figure 2, SD Figure 3-4, Supplementary Tables 3-6, 13, 14**). Cell states were further aligned to a mouse model of ischemia-reperfusion injury (IRI) and across the adult age-span, incorporating snRNA-seq and 10X Multiome data from 70 mouse samples (**ED Figure 3, SD Figure 5, Supplementary Table 15**)^14,18^. Integrative analyses identified 111 annotated cell types and 44 altered cell states with distinct marker gene profiles (**Supplementary Table 16, 17, SD Figure 5**). Alignment to HKAv2 revealed variable conservation of cell states across species, highlighting both similarities and differences, and the extent to which this injury model system may recapitulate the complexity of human kidney disease (**ED Figure 3)**.

**Figure 2.**
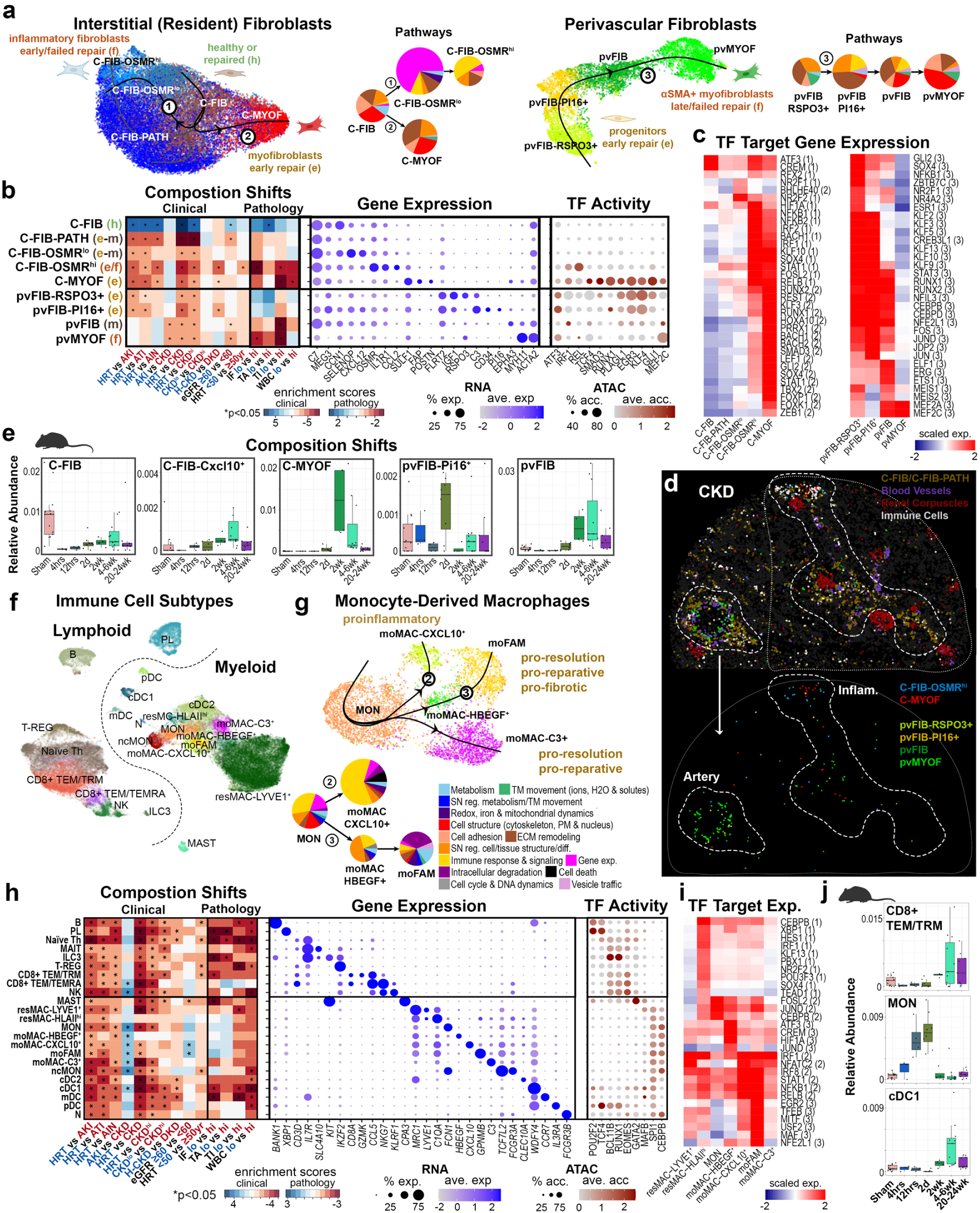
Fibroblast and immune cell types and states. **a**. UMAP embeddings showing interstitial (resident) and perivascular fibroblasts and their associated Slingshot predicted trajectories. Pie slices represent predicted fractional overall whole cell functions. Pie slice areas within the same trajectory network are proportional to the sum of the −log10 p-values of all grouped pathways for the indicated function (sum of all −log10(p) for C-FIB-OSMR^lo^ and pvMYOF: ∼124, ∼46, respectively). Arrows represent slingshot trajectories. For legend see (**g**). Numbers map to numbers in UMAP trajectories. **b**. Left, heatmap showing cell type enrichments as t statistics for the comparison of cell type abundance between two different patient groupings. Asterix indicates p values less than 0.05. Only biopsy samples (excluding nephrectomy and deceased donor) were used for all categories except age associations. Abbreviations, grouping strategies and number of patients per group are indicated in ED Figure 1b and legend. Middle, dotplot showing average expression values for selected marker genes. Right, dotplot showing average binding site accessibilities for selected TFs. **c**. Heatmap of scaled expression scores for TF target genes associated with GRNs built from trajectories shown in (**a**). Numbers indicate the specific trajectory used to derive the target gene set. **d**. Slide-seq2 processed CKD tissue area colored by fibroblasts: C-FIB/C-FIB-PATH, blood vessels, renal corpuscles and immune cells (top only), as well as C-FIB-OSMRhi, C-MYOF, pvFIB-RSPO3+, pvFIB-PI16+, pvFIB and pvMYOF (top and bottom). Tissue puck is 3mm in diameter. **e**. Bar plots showing relative abundance for different fibroblast states within mouse samples collected at different time points from IRI and compared to sham controls. **f**. UMAP embedding showing lymphoid and myeloid immune cell subtypes. **g**. UMAP embedding showing monocyte-derived macrophage subtypes and predicted Slingshot trajectories. Pie charts represent overall whole cell functions predicted for macrophage subtypes. Numbers indicate the specific trajectory used to derive the target gene set (sum of all −log10(p) for moMAC-CXCL10^+^: ∼86). **h**. Heatmap and dotplots for immune cells as in (**b**). **i**. Heatmap of scaled expression scores for TF target genes associated with GRNs built from trajectories shown in (**g**) and (**ED Figure 6d**). **j**. Bar plots showing relative abundance for different immune subtypes as shown in (**e**). Source data provided.

**Figure 3.**
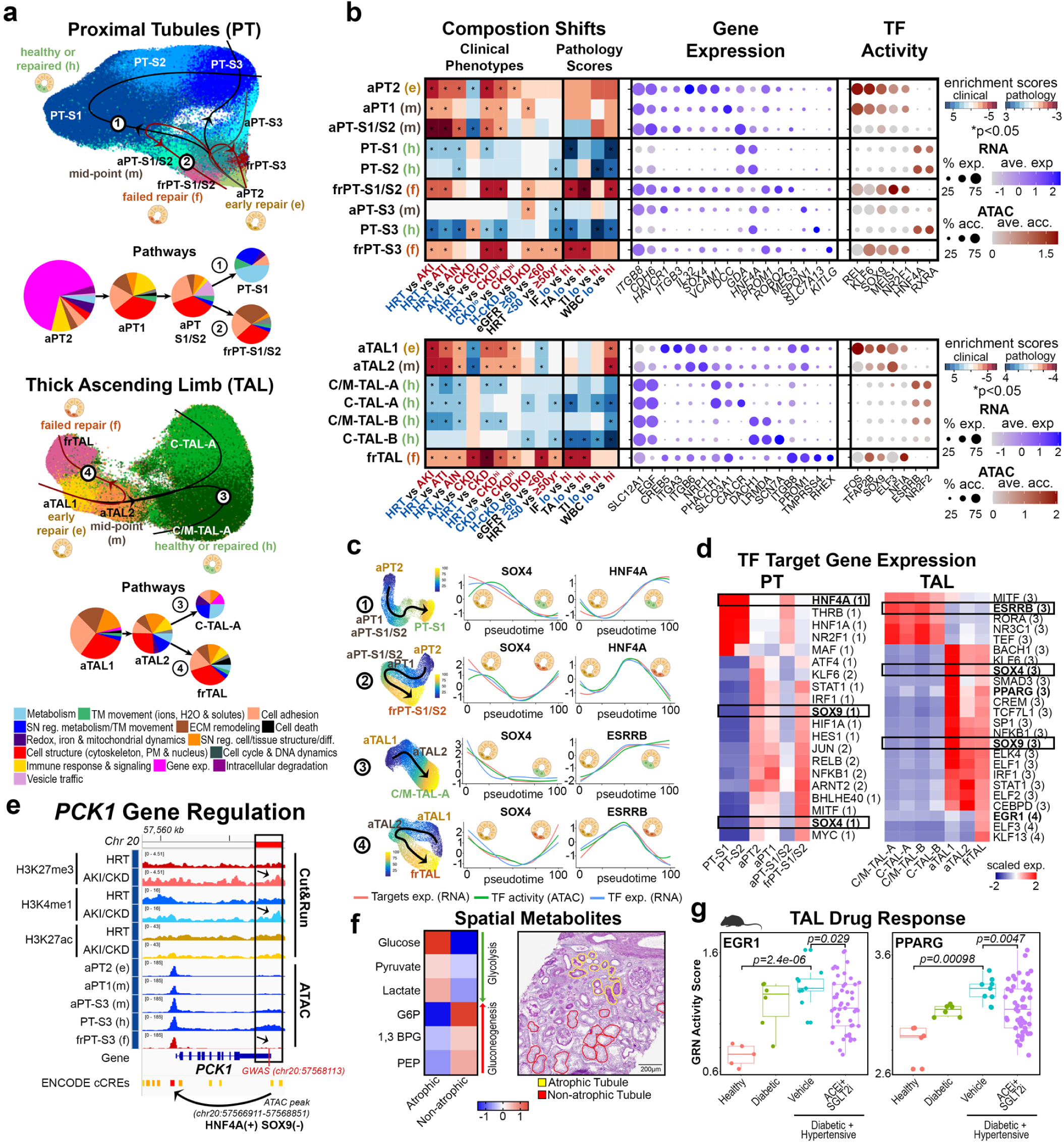
Resolving and unresolved epithelial repair states. **a**. UMAP embedding showing proximal tubule (PT) and thick ascending limb (TAL) states and associated Slingshot predicted trajectories. Pie charts represent whole cell functions predicted for proximal tubule and thick ascending limb subtypes (sum of all −log10(p) for aPT2 and aTAL1: ∼138 and ∼72, respectively). **b**. Left, heatmap showing cell type enrichments as t statistics for the comparison of cell type abundance between two different patient groupings. Asterisk indicates p values less than 0.05. Only biopsy samples (excluding nephrectomy and deceased donor) were used for all categories except age associations. Abbreviations, grouping strategies and number of patients per group are indicated in **ED Figure 1b** and legend. Middle, dotplot showing average expression values for select marker genes. Right, dotplot showing average binding site accessibilities for select TFs. **c**. Left, UMAP embeddings for individual trajectories shown in (**a**) colored by estimated pseudotime. Right, scaled target gene expression (RNA), binding site activity (ATAC) and gene expression (RNA) levels for select TFs as a function of pseudotime. **d**. Heatmaps of scaled expression scores for TF target genes associated with GRNs built from trajectories shown in (**a**). Numbers indicate the specific trajectory used to derive the target gene set. **e**. Genomic region of *PCK1* visualized with Cut&Run tracks for histone marks (H3K27ac, H3K4me1, and H3K27me3) in healthy reference and AKI/CKD kidney tissue, and pseudo bulk ATAC coverage tracks for PT trajectory states. Arrows indicate an HNF4A/SOX9 regulatory region (**Supplementary Table 24**) that overlaps with the GWAS SNP (rs2235826) and showed decreased open chromatin for adaptive PT states and increased repressive histone marks (H3K27me3 and H3K4me1) in AKI/CKD. **f**. Right: heatmap of normalized intensities of spatially localized glycolysis and gluconeogenesis metabolites in atrophic (n = 270) and non-atrophic (n = 270) tubules. Glucose, pyruvate, and lactate were used as indicators of glycolysis, while phosphoenolpyruvate (PEP), 1,3-bisphosphoglycerate (1,3-BPG), and glucose-6-phosphate (G6P) served as gluconeogenesis markers. Left: H&E image indicating atrophic or non-atrophic tubules, scale = 200µm. **g.** Activity of EGR1 and PPARG **r**egulatory networks (**d**) as ssGSEA scores (**see Methods**) in mouse models of diabetic and hypertensive kidney injury ^94^. ACEi = ACE inhibitor; SGLT2i = SGLT2 inhibitor. Source data provided.

**Figure 4.**
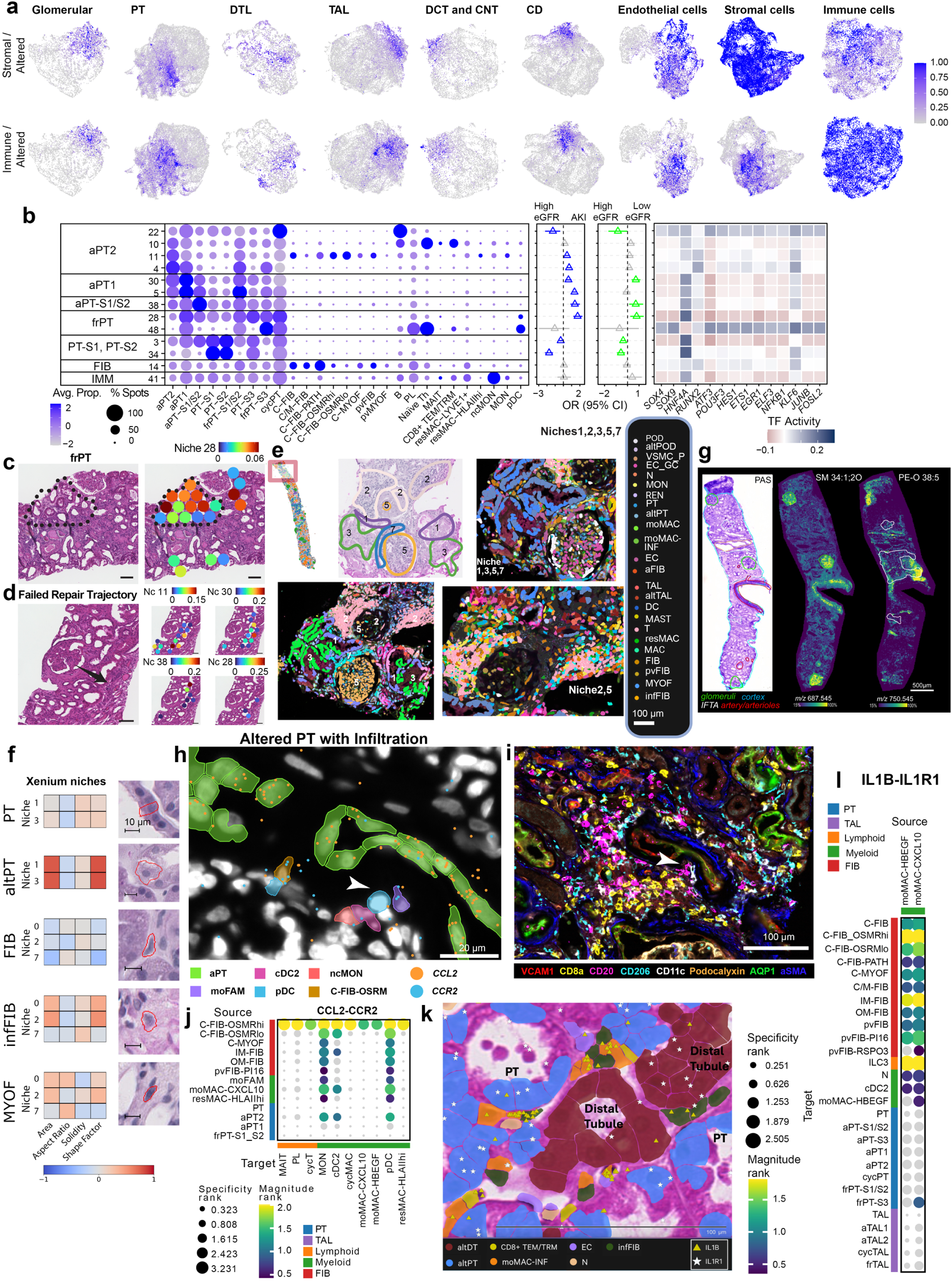
Spatial niches of Proximal Tubules. **a.** Distribution of altered states, stromal and immune cell types across the spots grouped by major nephron segments or kidney structures; the distribution is displayed as a ratio between stromal and altered or immune and altered (capped to 1). **b.** Characterization of selected PT niches with cell type distribution, log odds ratio of association with clinical status (with confidence interval), and transcription factor network activities. High eGFR samples had an eGFR above 60 ml/min/1.73 m^2^. **c.** Niches were mapped to FFPE samples. Here, the failed repair PT niche maps to an annotated region with evidence of tubular dilatation, epithelial simplification, immune cell infiltration, interstitial fibrosis and tubular atrophy, scale = 100 µm. **d.** A spatial distribution of niches demonstrates that the stages of the failed repair trajectory map over regions of tubular dilatation, epithelial simplification, intraluminal cast formation, and interstitial fibrosis. The failed repair trajectory is reflected in the juxtaposition of niches along the annotated arrow, scale = 100 µm. **e.** Neighborhood analysis on CKD samples using 10X Xenium. Results from a niche analysis using BANKSY software on four diabetic nephropathy samples (3 patients) are shown. **f.** Pathomic features of Xenium niches showing increased size of altered PT (altPT) cells, and larger, rounder inflammatory fibroblast (infFIB) subtypes across niches. **g.** Ion images of lipid biomarker candidates for IFTA (PA 32:0, *m/z* 647.466) and glomeruli (SM 34:1;2O, *m/z* 687.545. **h.** Xenium image showing a niche consisting of an injured proximal tubule expressing *CCL2* surrounded by macrophages, stromal and dendritic cells expressing *CCR2.*). **i.** A sequential CODEX image to (**h**) which visualizes the same adaptive PT (white arrow) with colocalized macrophages, T and B cells. **j.** Modelling of cell communication on snRNA-seq identifies the potential cellular communications between *CCL2* and *CCR2* expressing cells, confirming the cell types shown in (**h)** and (**i)**. Non-specific interactions are in grey. **k.** Xenium ST of the IL1B-IL1R1 niche**. l.** Liana+ analysis of IL1B-IL1R interaction between inflammatory macrophages, PT and TAL states that were colocalized in Xenium niches. Non-specific interactions are in grey. Created in BioRender. Jain, S. (2025) https://BioRender.com/h4kdr1x. Source data provided.

**Figure 5.**
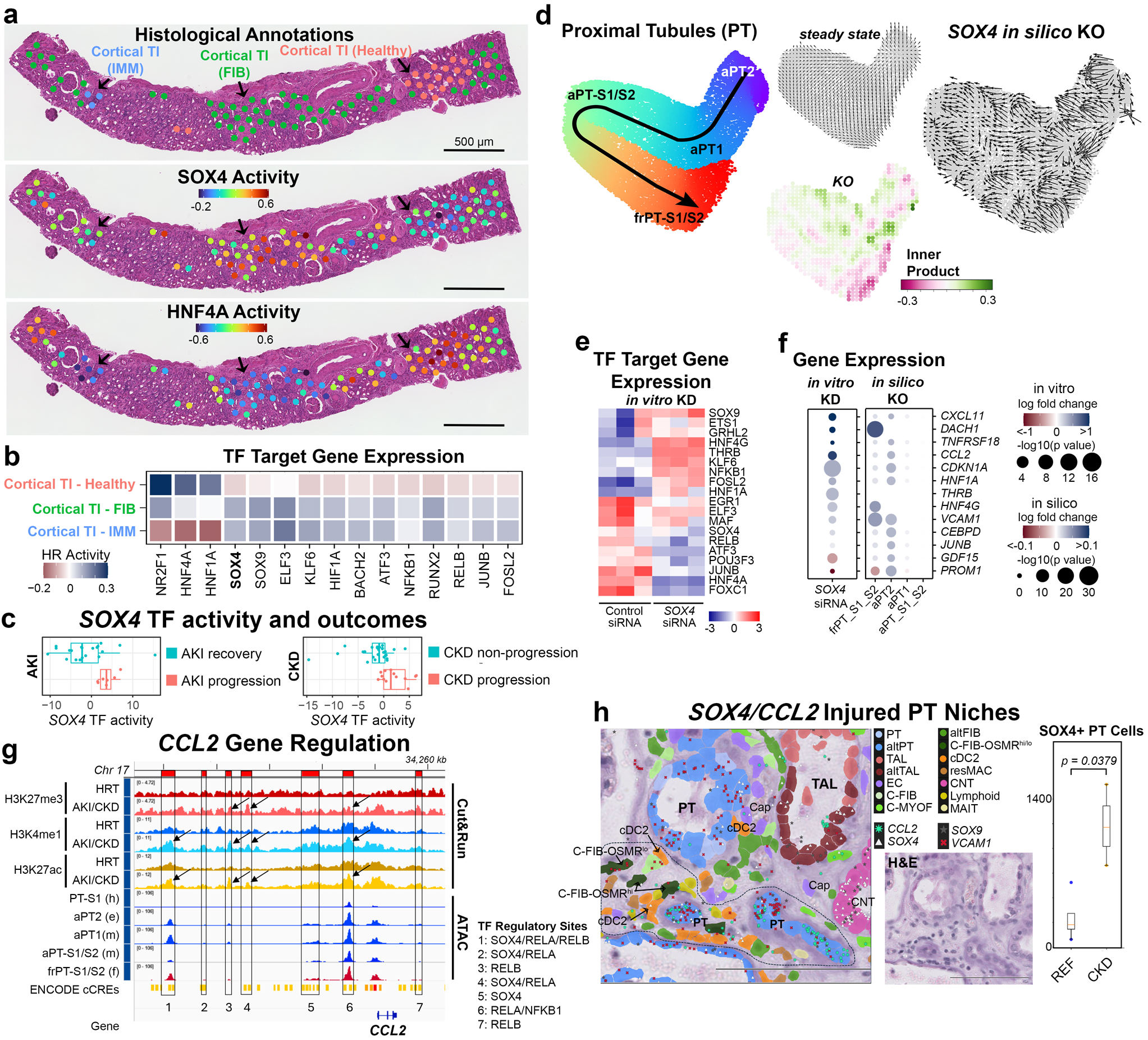
SOX4 regulation of aPT states *in vivo*. **a.** *SOX4* and *HNF4A* activities based on target gene expression calculated on FFPE samples. **b.** Heatmap of transcription factor activities predicted across histologically annotated spots (FFPE samples). **c**. SOX4 inferred transcription factor (TF) activity was mapped across all tubulointerstitial spots of frozen Visium samples. SOX4 TF activity was elevated in participants with AKI (N=27) who failed to recover (p = 0.038) and in participants with CKD and 2 or more years of follow-up (N=39) whose eGFR declined by 15% or more (p = 0.0026). **d.** *in silico* knock-out of *SOX4* within the PT trajectory **(Figure 3a)**. **e.** Heatmap of scaled expression scores for TF target genes associated with the failed repair PT-S1/S2 trajectory after *in vitro SOX4* siRNA knockdown in human proximal tubule cells (**Figure 3a). f.** Changes in gene expression for *in silico* knock out (KO) and *in vitro* knock down (KD) of *SOX4*. **g.** Genomic region of *CCL2* visualized with Cut&Run tracks for histone marks (H3K27ac, H3K4me1, and H3K27me3) in healthy reference and AKI/CKD kidney tissue, and pseudo bulk ATAC coverage tracks for PT trajectory states. Candidate regions (Supplementary Table 25) regulated by transcription factors (SOX4, NFKB1, RELA, RELB) upstream (regulatory regions 1-6) and downstream (regulatory region 7) of the *CCL2* gene. Arrows indicate regions showing histone marks that vary between healthy and disease samples. Distal regulatory region 1 showed increased chromatin activities in adaptive PT and active histone marks (H3K27ac and H3K4me1) in AKI/CKD (arrows). **h**. 10X Xenium data using the WU300KID panel showing fibro-immune environment surrounding altered tubules expressing *SOX4*. Note *SOX4* expression in altered PT, TAL and CNT cells. Dotted line indicates niche region containing C-FIB-OSMR^hi/lo^ and cDC2 cells surrounding altered PT cells (altPT) expressing *SOX4*. *CCL2* expression is seen in several of these alt PTs and in C-FIB-OSMR^hi^ fibroblasts forming a signaling axis with *CCR2-*expressing cDC2 cells to maintain the inflammatory environment, scale = 100µM. The bar graph shows a comparative analysis of *SOX4*+ cells in reference (4) and CKD (3) samples showing significantly higher number of *SOX4*+ cells in CKD (p = 0.037). Created in BioRender. Jain, S. (2025) https://BioRender.com/a50f9of. Source data provided.

To further increase molecular coverage of cell types and states, single cell RNA-seq data were independently clustered (**Supplementary Tables 18-19)** using complementary computational analyses with single nucleus RNA-seq. We further mapped functional pathways^19^ to the key functional tissue units (FTUs) of the nephron, interstitium and vasculature, stratified by disease condition or cell types (**Figure 1b**, **ED Figure 4**, **Supplemental Table 20**). Pathways predicted in the different FTUs for patients with mild CKD mostly described normal physiological functions (e.g., ion transport) while those for patients with advanced CKD or AKI suggested cell and tissue remodeling as well as immune response activities.

Utilizing the comprehensive ChemBL database comprising over 2 million compounds, we mapped putative drug targets across all the states across the nephron to find over 2,300 compounds with predicted activity in the kidney. We identified several approved drugs with known target structures in the nephron segments but now provide greater cell-type specificity (**SD Figure 6)**. We also identified three highly cell type-specific candidate drugs (Sacubitril, Venetoclax and Tolrestat) (**Figure 1c).** Our findings suggest that Venetoclax, a chemotherapy agent, may exert kidney toxicity on injured PT, possibly contributing to AKI or reduced recovery for patients exhibiting AKI or CKD ^20^.

**Figure 6.**
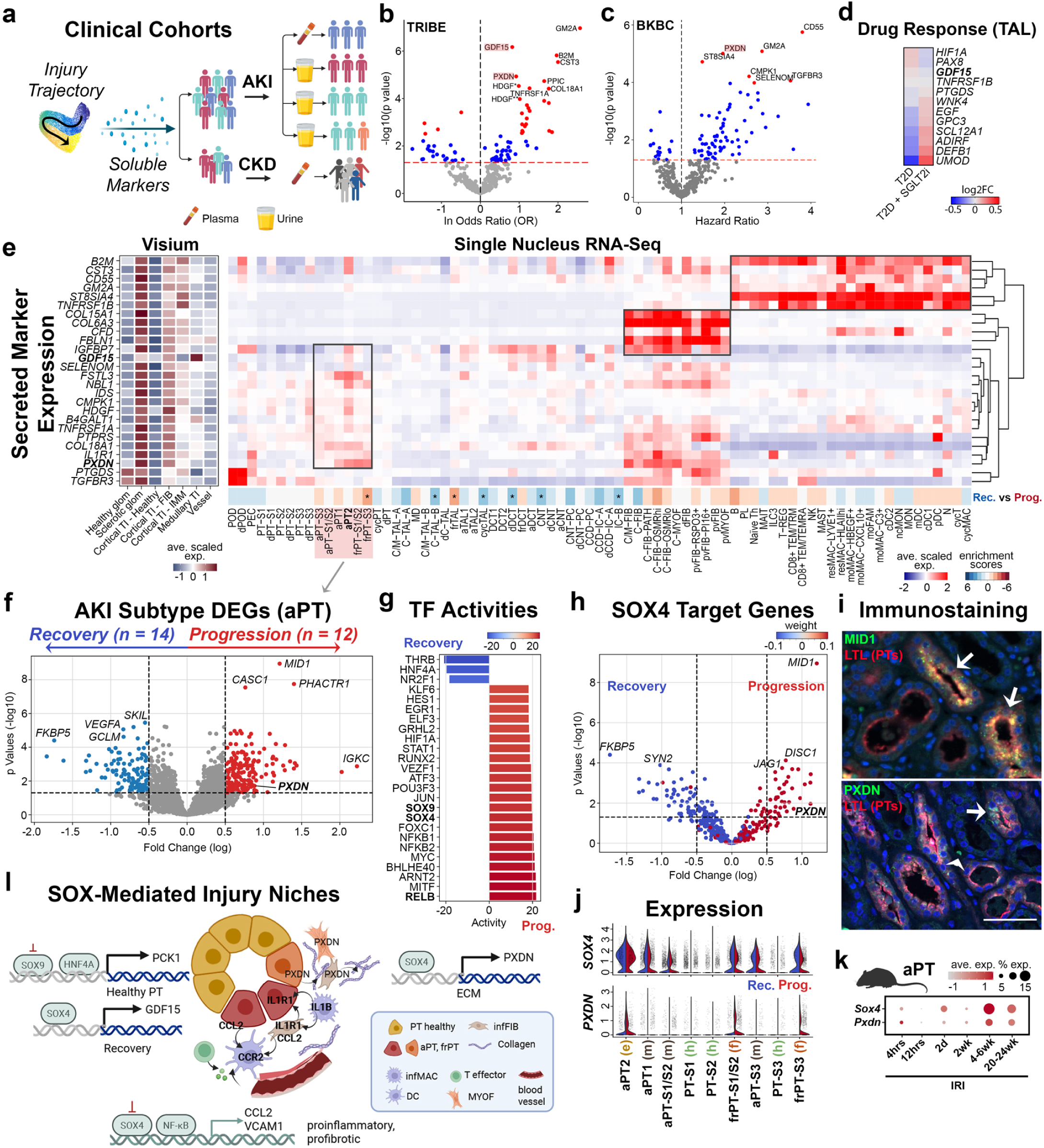
Clinical insights from HKAv2. **a.** Illustration summarizing application of atlas insights to clinical relevance in outcomes using plasma and urine proteomics from external cohorts. **b**. Circulating plasma proteins associated with AKI in TRIBE study, adjusted for age, sex, diabetes, and baseline eGFR. The red dotted line represents the naïve α = 0.05 significance level on the −log10 scale, red dots represent adjusted p values (Benjamini-Hochberg) < 0.05. **c**. Circulating plasma proteins associated with development of end-stage kidney disease in the Boston Kidney Biopsy Cohort adjusted for age, sex, race, eGFR, and proteinuria. The horizontal red dotted line shows the significance threshold at p=0.05 and markers highlighted in red show adjusted p values (Bonferroni) < 0.05. Markers highlighted in red remain significant after Bonferroni correction (Bonferroni-adjusted p value < 0.05). The vertical dotted line marks the Hazard Ratio of 1. **d.** SGLT2i treatment leads to increased expression of TAL recovery markers and reduction in injury markers. **e**. Cell type expression of secreted markers identified from clinical cohorts as associated with AKI or CKD progression. **f**. aPT state genes associated with recovering or worsening kidney function after up to 18 month follow up of KPMP AKI patients. **g**. TF activities associated with AKI progression. **h**. SOX4 target genes associated with AKI recovery or progression as in (**f**). **i**. Staining for the PT-specific Lotus tetragonolobus lectin (LTL) and either MID1 on healthy reference (top) or PXDN on AKI (bottom) tissue sections, scale = 50µm. **j**. Expression of *PXDN* and *SOX4* in human aPT states. **k.** Expression of *Pxdn* and *Sox4* during the mouse IRI time-course. **l**. Schematic showing SOX4 regulatory activities that may drive progression of unresolved repair niches. Source data provided.

### Steady-state and pathogenic fibroblast subtypes

#### Interstitial fibroblasts

A central response to tissue injury is the activation of fibroblasts for ECM deposition, scar formation and stabilization of organ integrity^21^. Any failure of resolution may lead to fibrosis and declining organ function, necessitating discovery of potential fibroblast subtypes associated with resolving and unresolving repair (**Figure 2**). Fibroblasts represent 5.9% of the total cells (N = 1.3 million) sampled by single nucleus sequencing. This deep sampling of the HKAv2 atlas has enabled sufficient fibroblast coverage to reveal rare subtypes, including the spectrum of heterogenous populations that span the cortico-medullary axis (**ED Figure 5a, Supplementary Table 10**). This includes interstitial or resident fibroblast subtypes that become expanded in injury or disease (C-FIB-PATH) or that may progress along trajectories either to a cross-tissue associated CXCL10+/CCL19+ pro-inflammatory state (C-FIB-OSMR^hi^) or a SPARC+COL3A1+ myofibroblast state (C-MYOF) (**Figure 2a**, **ED Figure 5b, Supplementary Table S21**)^8,15,22^.

Pathway enrichment analysis suggests high immune-response, in particular complement-related activities for C-FIB-OSMR^hi^ and interstitial ECM remodeling, including collagen synthesis activities for C-MYOF. C-FIB-OSMR^hi^ fibroblasts were highly associated with interstitial fibrosis and tubular injury (**Figure 2b**), localized to T-cell enriched niches^15^, and expressed genes associated with the ability to respond to and produce inflammatory cytokines, including *IL1R1*, *CCL2*, *CXCL10, CCL19* and the Oncostatin-M receptor (*OSMR*) (**Figure 2b**, **SD Figure 2, Supplementary Table 12**).

To identify robust gene regulatory networks (GRNs) underlying cell state shifts, we used two different approaches, scMega and MAGICAL, that link transcription factor binding site activities to accessible chromatin peaks in trajectory-relevant genes^23,24^ (**Supplementary Table 22**). The C-FIB-OSMR^hi^ fibroblasts, which showed activation of NF-kB, TNFα, and interferon (INF) pathways, were associated with increased TF activity of *NFKB1*, *NFKB2*, *REL* and *IRF1* (**Figure 2b-c, ED Figure 5c-d**). Alternatively, the C-MYOF subtype showed TGFβ pathway expression with a corresponding increase in SMAD3 activity (**Figure 2b-c, ED Figure 5c-d**). These cells showed expression signatures consistent with a vascular niche associated myofibroblast subtype seen in injured and inflamed tissues^15,22^, including the expression of *POSTN*, *FAP* and *INHBA* (**Figure 2b**, **SD Figure 2, Supplementary Table 12**). Both subtypes became more enriched weeks after the acute injury in the mouse model, indicating potential roles in the fibrotic phase (**Figure 2d**). Consistently, the C-MYOF subtype showed the most significant enrichment for the synthesis of collagen and other interstitial ECM components (**Figure 2a)**^25^, was enriched in patients with tubular injury and high white blood cell (WBC) scores (**Figure 2b**) and was found colocalized to areas of fibrosis and immune infiltration (**Figure 2e, ED Figure 5e**). In summary, we identified two interstitial fibroblast states, immunomodulatory and matrix producing, that were shared across tissues and may have become activated by T-cell or vascular endothelial cell (EC) signals^15^. Shifts to these interstitial fibroblast states were conserved in the mouse IRI model, with the exception of an additional early activated fibroblast state (eaC-FIB) that was enriched within hours of the acute injury and that may also have given rise to the C-MYOF subtype (**ED Figure 5f-g**).

#### Perivascular fibroblasts

We identified distinct perivascular fibroblast subtypes that correspond to those annotated in the kidney and other organs. For instance, pluripotent universal RSPO3+ and CD34+/PI16+ adventitial fibroblasts which are known to give rise to multiple fibroblast subtypes^8,15^. These were rare in healthy kidney tissues, localizing to afferent arterioles and the descending vasa recta (**ED Figure 5h**). The pvFIB-RSPO3+ progenitor population expressed epithelial development signals (*WNT5B, IGF1*) and showed GLI1 TF activity, known to regulate αSMA+ myofibroblast transition in models of heart, kidney, lung and liver fibrosis^6^.

A second progenitor subtype (pvFIB-PI16+) expressed *CD34, MFAP5* and *PI16*, a shear-stress regulated matrix metalloproteinase (MMP) inhibitor which senses fluctuations in kidney blood flow^26^ for possible regulation of vascular stress and ECM remodeling. These cells were enriched in AKI (**Figure 2b**) and early time-points following IRI in the mouse (**Figure 2d**) and may act as sentinels that sense local injury and promote either immune infiltration^27^ or tissue regeneration^28^. Slingshot trajectory analyses predicted their differentiation into derivative fibroblast (pvFIB) and αSMA+ myofibroblast (pvMYOF) states that were expanded in CKD patients with high interstitial fibrosis and tubular injury, or in mice at late time-points following IRI (**Figure 2a-b, d, ED Figure 5f, Supplementary Table 21**). This suggests potential alignment of this molecular trajectory with disease progression from AKI to CKD. Consistently, the latter perivascular fibroblast subtypes were spatially localized to small arteries and arterioles in disease, with expansion into the interstitial space, and the gain of pathways associated with actin filament dynamics and contraction (**Figure 2a-b, ED Figure 5h**)^29,30^.

The perivascular fibroblast subtypes may also give rise to interstitial fibroblast or myofibroblast subtypes, as indicated from lineage tracing in mice^8^. A recent perturbation study found that the key regulators of the CD34+PI16+ state included KLF4 and KLF5, also seen in our human kidney data (**Figure 2c, ED Figure 5d**), activation of which could shift fibroblasts towards this progenitor state *in vitro* while suppressing the SPARC+COL3A1+ (C-MYOF) state^31^. These cell state shifts indicate a plasticity in the interstitial and perivascular subtypes that provide insights into key regulatory processes. One identified TF was SOX4, which demonstrated regulatory activity both during shifts to C-FIB-OSMR^hi^ and C-MYOF states, but also within the perivascular fibroblast progenitors (**Figure 2c**), positioning this TF as a key regulator of the repair niche. These transitions were well conserved in the mouse IRI model which showed conserved expression profiles for human TF regulatory networks (**ED Figure 6a**). Therefore, the HKAv2 has enabled the discovery of distinct fibroblast subtypes associated with fibrosis, contractility and inflammation that may colocalize to niches of resolving and non-resolving epithelial repair in AKI and CKD.

### Clinicopathologically-linked immune subtypes

The immune cell subtypes that colocalize in the pro-fibrotic and pro-inflammatory fibroblast niches may ultimately influence repair outcome. To identify states associated with resolving and non-resolving repair, we first clustered snRNA-seq data, then performed integration with scRNA-seq data to better classify these subpopulations (**SD Figure 7, Methods**). An improved sampling depth within the HKAv2 enabled greater resolution of immune subtypes, including extensive heterogeneity in lymphoid and myeloid cells with enrichment in AKI or CKD patients (**Figure 2f-h**). While all lymphoid cells were enriched for both acute and chronic injury, terminally differentiated cytotoxic T cells (CD8+ TEM/TEMRA) and natural killer cells (NK) were more enriched in AKI patients (**Figure 2h**). Alternatively, plasma cells (PL), naïve T cells, innate lymphoid cell subpopulation III cells (ILC3) and cytotoxic T cells (CD8+ TEM/TRM) were all found to be enriched with high interstitial fibrosis, with several of these subtypes also enriched in patients having high risk for end-stage kidney disease (CKD^hi^, **Figure 2h**). This is consistent with lymphoid cells showing late enrichment within mouse IRI tissues corresponding to the fibrotic phase (**SD Figure 5c, ED Figure 6b**).

The HKAv2 also resolved heterogeneous populations of myeloid cells, including cross-tissue and evolutionarily conserved subtypes (**ED Figure 6c**) ^9,11^, that showed early enrichment following injury (**Figure 2h, SD Figure 5c**). This heterogeneity suggests activation of both regenerative and remodeling roles to phagocytose injured tissue, promote immune cell recruitment and remodel the ECM, in part through polarization of fibroblasts to myofibroblasts ^17^. Tissue resident macrophages expressing LYVE1 (resMAC-LYVE1+), potentially originating near blood vessels ^32^, were expanded in AKI (**Figure 2h**), with a corresponding subtype in mice (resMAC) that became expanded within two days after IRI (**SD Figure 5c, ED Figure 6b**). resMAC-LYVE1+ cells expressed genes associated with efferocytosis (*MERTK*), endocytic internalization pathways and growth factors associated with tissue repair (*IGF1*, *PDGFB* and *PDGFC*), and this state remained expanded in CKD patients and late fibrotic states of the mouse injury model. Trajectory analyses predicted a shift of these cells into a M2-like pro-resolving resMAC-HLAII^hi^ state that was more associated with acute tubular injury (ATI) (**ED Figure 6d; Supplementary Table 21**). This state displayed pathways involved in lysosomal antigen degradation ^33^ and exogenous and endogenous antigen presentation. This is consistent with resMAC-LYVE1+ and resMAC-HLAII^hi^ representing previously characterized polarization states that may ultimately originate from CD14+ monocytes ^11^. In mice, the resMAC-HLAII^hi^-associated state (resMAC-H2^hi^) arose after the onset of resMAC expansion (**SD Figure 5c, ED Figure 6b**) and persisted into late phases following acute tubular injury (ATI), consistent with a potential role in the resolution of fibrosis that was observed in this mouse model. In support of this, we find low enrichment for resMAC-HLAII^hi^ in human CKD patients (**Figure 2h**). Polarization to this reparative M2-like state was associated with the Hippo signaling pathway mediator TEAD1 ^34^ as well as the activities of several transcription factors that may play a role in suppressing inflammation, including CEBPB, PBX1, and Notch pathway mediator HES1 ^35–37^ (**Figure 2i, ED Figure 6e**). These results are consistent with pro-reparative and pro-resolution roles of the tissue-resident macrophages following acute injury.

A return to tissue homeostasis may ultimately rely on the balance of pro-resolution and pro-inflammatory myeloid states ^17^. To identify candidates that may promote inflammation and possible disease progression, we looked for possible polarization states of infiltrating monocytes, which were enriched in pathways of immune response (**Figure 2g**). *FCN1*+ classical monocytes (MON) were more enriched in AKI over CKD and showed an early infiltration into the mouse kidney within 12 hours of ATI (**Figure 2h-i**, **SD Figure 4c**). Their potential derivatives showed persistent enrichment in CKD patients and late fibrotic-stage mouse kidneys (**Figure 2h, ED Figure 6b**, **SD Figure 5c**). These MON cells polarized towards an M1-like pro-inflammatory *CXCL10*+ *CXCL9*+ *CCL2*+ state (moMAC-CXCL10+, **Figure 2g, ED Figure 6e-f**), that was associated with inflamed tissues and induction by IFN-gamma and TNFα signaling^38,39^. Consistently we found the polarization to this state was regulated by IFN-associated TFs IRF1 and IRF8, as well as NF-kB pathway-associated NFKB1 and RELB, potential mediators of TNFα signaling ^38^ (**Figure 2g, ED Figure 6e**). While myeloid subtypes in general were less well conserved between humans and mice (**ED Figure 3b**), this inflammatory state was observed after mouse ATI where it was enriched at late timepoints (>20 weeks post IRI) concomitant with the enrichment for plasma (PL) and T cells (**SD Figure 5c, ED Figure 6b**). Therefore, this state may represent a late pro-inflammatory macrophage that promotes lymphoid cell recruitment and potential disease progression.

We also found another inflammatory population marked by expression of growth factors *HBEGF* and *AREG*, as well as proinflammatory genes *PLAUR*, *IL1B, OSM* and *CXCL8*, and that is consistent with a subtype found in rheumatoid arthritis ^40^ (**Figure 2g-h, ED Figure 6f**). This subtype (moMAC-HBEGF+) expressed genes that have been shown to promote parenchymal cell proliferation or regeneration (*AREG*, *IL1B* and *OSM*), which chronic exposure would impair (*IL1B*), supporting its role in an early transitory state (**ED Figure 6f**) ^17^. Shift to this state was regulated in part by HIF1A (**Figure 2i**), consistent with Wnt-mediated HIF1A promoting a pro-inflammatory state in the lung ^41^. These cells may also represent an intermediate population moving towards a *GPNMB*+ *SPP1*+ polarized subtype previously described as deriving from FCN1+ monocytes and resembling lipid-associated macrophages (LAM), scar-associated macrophages (SAM), tumor-associated macrophages (TAM) and disease-associated microglia (DAM) ^11,42^. Given the link to fibrosis across tissues ^42^, we dubbed these monocyte-derived fibrosis-associated macrophages (moFAM). However, this subtype also expressed anti-inflammatory *TREM2,* and showed high lysosomal degradation activities that may support clearance of debris and dead cells as well as genes consistent with pro-resolution and pro-repair (**ED Figure 6f**) ^17^, indicating diverse roles in injured tissue. moFAM cells were enriched in AKI more so than CKD, but not within tissues showing high interstitial fibrosis, indicating early repair activities (**Figure 2h**). Unlike moMAC-HBEGF+, the moFAM state was conserved in the mouse IRI model, arising early at around two days after ATI (**SD Figure 5c**). This supports a potential role for moFAM in early repair, and a continued presence at late fibrotic states that may be consistent with both pro-resolution and potentially pro-fibrotic roles. As found previously, we identify TFEB as a key transcription factor mediating the polarization to this moFAM state ^43^ (**Figure 2i, ED Figure 6e**). Therefore, we find that infiltrating monocytes show a strong enrichment after acute injury and may contribute to pro-inflammatory and M1-like macrophages, as well as potential pro-repair and pro-resolution subtypes. Their continued enrichment in CKD patients indicates potential contributions to ongoing inflammation and a likely basis for AKI-CKD disease progression. We also find that MAST and dendritic cells (cDC1 and mDC) show significant enrichment in patients showing high interstitial fibrosis (**Figure 2h**), with dendritic cell subtypes also arising late following mouse IRI (**Figure 2j**, **SD Figure 5c**). This potentially suggests additional roles for these cells in fibrosis.

### Resolved versus unresolved epithelial repair

Injured epithelia can secrete cytokines, chemokines and growth factors for activation of local macrophages and fibroblasts, which through reciprocal interactions support transitional epithelial repair states within the niche^21^. Dysregulated communication within these niches can lead to a failure to resolve epithelial repair, which has been linked to the progression to end stage kidney disease ^44^ and idiopathic pulmonary fibrosis ^45,46^. The injured tubular epithelial cell states are associated with a mesenchymal-like signature, downregulated tight-junctions, partial loss of apical-basal polarity and the upregulation of developmentally relevant genes. Degenerative states were frequently enriched for pathways involved in translation, mitochondrial homeostasis, oxidative phosphorylation and iron storage, pathways often associated with oxidative stress and ferroptosis in kidney injury (**ED Figure 4, Supplementary Table 20**) ^47,48 49^. In adaptive and failed repair states, structural pathways like cell adhesion and ECM dynamics were enriched. The failed repair proximal tubule (PT) and thick ascending limb (TAL) subtypes also showed expression of cell cycle checkpoint genes, consistent with cell cycle arrest during progressive tubular injury^44,50^.

The larger disease sample size in HKAv2 allows distinction of resolving and failed or unresolved repair states within the PT, TAL and distal convoluted tubule (DCT) (**Figure 3, Supplementary Table 10)**. To orient cell populations along paths for successful or failed repair, we inferred slingshot lineages for both the PT and TAL (**Figure 3a, SD Figure 8, Supplementary Table 21)**. This enabled the alignment of early through mid-repair substates that then progressed either to healthy / recovered states or to unresolved / failed repair states. Consistently, these states showed signatures associated with epithelial cell repair, integrity and differentiation, and unresolved states were enriched in patients with CKD, low eGFR, and high interstitial fibrosis (**Figure 3b**). Early injury states (aPT2 and aTAL1) showed expected expression of gene signatures associated with cell adhesion and cytoskeletal dynamics, inflammation, TGFβ and / or NF-kB signaling, as well as genes encoding plasma proteins enriched in ATI (**Figure 3c, ED Figure 7a**) ^51–53^. aPT2 expressed the injury markers *CDH6, VCAM1 and HAVCR1* with REL (NF-kB) and KLF6 binding site activities (**Figure 3b**) ^1,13,18,54,55^. *EGF* was downregulated in aTAL1, while *CREB5*, *ITGA3, and ITGB6* were upregulated with increased activity for activator protein AP-1 and AP-2 signaling pathways (FOS and TFAP2B, respectively), both associated with stress response and distal epithelial repair (**Figure 3b**) ^1^. Successful repair accompanied the restoration of canonical functions (metabolism and reabsorption), while failed repair was associated with persistence of ECM remodeling, cytoskeleton and cell adhesion dynamics as well as cell cycle checkpoint regulation ^44^. Failed repair (fr) states were further marked by the expression of *PROM1* (frPT and frTAL), *ROBO2* and *MEG3* (frPT), or *ITGB8* and *TMPRSS4* (frTAL) ^1,56,57^.

Previously, SOX9, an injury-induced pioneer TF, was found to regulate the transition between early kidney injury to fibrosis ^58^, which is supported by the HKAv2 (**Figure 3b, ED Figure 7a, Supplementary Table 23**). We additionally find a key role for another pioneer TF, SOX4, which shows similar regulatory activities across these epithelial trajectories (**Figure 3c,d**, **ED Figure 7b-g**). Early PT and TAL repair states both increased activity of SOX4/SOX9, which, as potential regulators of developmentally relevant pathways, became inactive upon injury resolution (SOXon/off) concomitant with restored activity of HNF4A or ESRRB (**Figure 3b-d**). However, in unresolved trajectories, a failure to inactivate (SOX9 on/on) was associated with low HNF4A/ESRRB activities, and an expected failure to re-establish full apical-basal polarity ^58^. This suggests important and potentially cooperative roles for these SOX TFs in regulating the balance between successful and failed epithelial repair. Consistent with this, GWAS analysis of SNPs associated with estimated glomerular filtration rate (eGFR) ^59^ (**Supplementary Table 24**) identified a variant within a SOX9 (negative) and HNF4A (positive) regulatory site of the Phosphoenolpyruvate carboxykinase 1 (*PCK1*) gene (**Figure 3e, Supplementary Table 25**). Critical for normal tubular physiology and gluconeogenesis, loss of this metabolic factor has been associated with tubular injury and compromised kidney function ^60^. Consistently, *PCK1* expression was reduced in altered PTs progressing to failed repair states (**ED Figure 7h-i**). Spatial metabolomics analysis revealed suppression of gluconeogenesis and increased glycolysis in atrophic compared to non-atrophic tubule regions, confirming a functional role of PCK1 in healthy tubules in CKD (**Figure 3f**). In addition, the citric acid cycle, purine metabolism, glycerolipid metabolism, arginine biosynthesis, biosynthesis of unsaturated fatty acids, and glutathione metabolism were dysregulated in atrophic tubules (**ED Figure 7j)**. As several of these pathways have been linked to progression of CKD ^61–63^, these functional readouts highlight a potential regulatory mechanism by which unresolved SOX activity may further promote failed repair through the establishment of an altered metabolic state within these epithelial cells.

In addition to SOX9 and SOX4, we identified several TFs and their associated GRNs across both the PT and TAL trajectories, including key regulatory roles for EGR1 and the previously characterized ELF3 in unresolved repair (**Figure 3c-d**, **ED Figure 7e-f**, **Supplementary Table 21**) ^54^. These regulatory processes were further conserved in the mouse model system (**ED Fig 7g**), with EGR1 and PPARG regulatory networks showing reversal by ACE and sodium-glucose cotransporter-2 (SGLT2) inhibitors in diabetic and hypertensive TAL (**Figure 3g**). This demonstrates the clinical relevance of these factors in driving unresolved cell states.

### Niches of resolved and unresolved repair

To decipher the spatial organization and cellular communications between altered cell states and their cellular environment, we used 10X Visium ST deep sampling (135 participants) to generate niche maps of major nephron segments (**SD Figure 9**). Our analysis revealed immune and stromal niches that colocalized with altered cell states in all major nephron segments and functional tissue units (FTUs) (**Figure 4a**). We next explored tubule-dominant niches (PT, TAL) that aligned with either healthy states or epithelial repair trajectories that were enriched in AKI/CKD or that colocalized with immune or stromal cells **(Figure 4b, ED Figure 8a-b)**. Niche compositions for PTs revealed a progression from early injury to failed repair (**Figure 4b**). Early niches co-localized with signatures for healthy PT (niches 3, 34), cycPT with B cells (niche 22), lymphocytes (niche 10) or fibro-immune cell types (niche 11). Alternatively, late niches associated with frPT (28 and 48) enriched for plasma, T and dendritic cells. These trends were similarly observed for aTAL niches (**ED Figure 8a-b**) and aligned with the human and murine snRNA-seq abundance trends (**Figure 2**). Transcription factors driving epithelial repair states (**Figure 3d**) inferred in these niches showed consistent trends, including early KLF6/HES1 activities and oscillatory SOX4/SOX9 (**Figure 4b, ED Fig 8a**). Finally, we identify a fibrosis-associated niche (14) enriched for aPT1 and frPT-S1/S2s, altered fibroblasts, myofibroblasts, and immune cells. In addition, we find several temporal stromal niches that were enriched in AKI/CKD that colocalized C-FIB-OSMR^hi^, plasmacytoid (p) DC, and B lymphocytes (niche 7) with high inferred PRRX1, IRF2, and IRF1 TF activity (**ED Figure 8c-k**). In summary, we defined the niche trajectories with their respective tubular and interstitial temporal repair states, transcription factor activities and clinical associations.

To improve histologic resolution, Visium niches were projected to formalin-fixed-paraffin-embedded (FFPE) samples (N=12) ^64^ with annotated regions of interstitial fibrosis and tubular atrophy or immune infiltration (**SD Figure 10**, **Figure 4c**). frPT niches more frequently mapped to regions of the cortex having chronic tubular injury, fibrosis and immune infiltration. We identified a transition of niches from regions of acute injury (11, 30, & 38) to chronic injury (28), providing spatial context to the temporal frPT trajectory (**Figure 4d**). Since frPT states were enriched for pathways of ECM synthesis and enzymatic processing **(Supplementary Table 20),** this suggests that tubule-interstitial crosstalk during injury ^65,66^ may have an effector role in scar formation and matrix deposition.

To further understand how repair states organize and communicate within niches defined at the cellular level, we incorporated additional ST technologies (Slide-seq2, CosMx and Xenium ST) that had single-cell resolution yet lower sampling coverage (**Supplementary Tables 4-6**). This enabled placement of more rare perivascular fibroblast progenitor subtypes in close proximity to lymphoid cells (**ED Figure 9a-e**), consistent with their T-cell mediated expansion ^12^. In support of this, neighborhood analysis found that these progenitor states became more associated with lymphoid cells in diseased tissues (**ED Figure 9f**). Similarly, the polarized macrophage state, moMAC-CXCL10+, and associated IRF TF-regulated network, also localized to this area of fibrosis and immune infiltration in CKD tissue (**ED Figure 9g**). Specific cytokine signals have been shown to promote these fibroblast progenitor phenotypes within tumor-adjacent regions, including CXCL12, IL6 and LIF ^12^. Interestingly, the IL6 family members IL11 and LIF were both expressed by early repair TAL states (aTAL1), and pvFIB-PI16+ cells were enriched adjacent to frTAL in disease (**ED Figure 9h**) indicating a potential association between the repairing TAL and these progenitor states located on the nearby arteries and arterioles. This is supported by the spatial co-localization of pvFIBs with vascular smooth muscle in Visium ST niches (**Extended Data Figure 9i-j)**. Spatial mapping of GRN-associated TF activities confirmed proximity of aPT/frPT and frTAL (SOX9 and ELF3) with C-FIB-OSMR^hi^ (IRF2) and C-MYOF (RUNX2) within CKD tissues, providing further evidence for the association of altered epithelial states with fibrosis and inflammation (**ED Figure 9f-g**).

Neighborhood analyses of Xenium ST data with associated histopathology confirmed the juxtaposition of PT injury with niches consisting of varying immune and fibroblast cell states (**Figure 4e, ED Figure 10a).** These niches, similarly identified in a second cohort of diabetic nephrectomies (10 samples, **ED Figure 10a**), showed co-localizations suggestive of niche communities that drive failed repair (e.g. Xenium altered PT niche #1 and #3 next to fibro-immune, lymphoid, or myeloid niches 2 and 0). Interestingly, we identify distinct fibrotic niches, enriched with ECM-producing fibroblasts (Xenium niche 7) or fibro-immune cell states (Xenium niche 2) (**ED Figure 10a)**. These latter niches were enriched for inflammatory fibroblasts (C-FIB-OSMR^hi^), macrophages (moMAC-HBEGF/moMac-Cxcl10^+^) and lymphoid cells, consistent with that found for Visium ST (**Figure 4b**, **ED Figure 8c**). To histologically ground these niches, we integrated the ST data with histology by leveraging our recently developed histo-omic mapping tool FUSION ^67^. FUSION facilitated the extraction of pathomic features associated with morphological cell state transitions within these niches, including the loss of a cuboidal shape for altered epithelial (PT) cells (**Figure 4f).** This also revealed distinct morphological changes in fibroblast states, with inflammatory fibroblasts in fibro-immune niches showing a shift from spindle-like to a more rounded shape consistent with activation by inflammatory cytokines like IL-1B ^68,69^. This is in contrast to myofibroblasts within fibrosis niches (7) showing more elongated morphologies consistent with αSMA deposition and contraction ^68^.

Using imaging mass spectrometry (5 biopsy specimens), we confirmed the enrichment of pro-inflammatory biomarkers in pathologically assigned areas of interstitial fibrosis and tubular atrophy (IFTA, **Figure 4g, ED Figure 10**). This identified PA 32:0 (m/z 647.466), PA 36:1 (*m/z* 701.524) and PE O-38:5 (m/z 750.545) as robust IFTA biomarkers associated with promoting inflammation. PE O-38:5, an ether linked lipid may regulate lipid raft signaling domains in innate immune cells as well as T and B cell receptors ^70^, and PA 32:0 and PA 36:1 may regulate immune responses by inducing TNF alpha, IL-1β and IL-6 ^71^. Given the extensive fibro-immune co-localizations and inflammatory biomarkers associated with IFTA, we examined putative ligand-receptor (L-R) cell-cell communications to better understand how PT and TAL states might communicate within their niches to drive injury progression (**ED Figure 10, 11, SD Figure 11**). Early aPT (aPT2) cells showed elevated immune L-R signaling, including CD59-CD2 (lymphoid), CALM1-KCNQ5 (TEM, PL, B), RPS19-C5AR1 (myeloid), HLAA-APLP2 (myeloid), and VIM-CD44 (pan-immune). Within the TAL, all altered states (aTAL1, aTAL2, and frTAL) showed elevated immune signaling through ADAM10 with distinct immune receptor specificity: CADM1 (cDC1), IL6R (mDC), TSPAN5 (MAIT), TSPAN14 (ncMON), EPHA3 (MYOF). We also find that failed repair states (frPT and frTAL) showed consistent signaling to fibroblasts states through NRG3-EGFR.

In addition to these, our spatial analyses have identified multiple niches co-localizing cells that were potential sources of CCL2 and IL1B (**Figure 4h-l**), both associated with CKD progression ^5,18,72^. Fibro-immune niches consisted of inflammatory macrophages (moMAC-HBEGF^+^ and moMAC-CXCL10^+^ both expressing *IL1B*) adjacent to CD8^+^ TEM/TRM, altered PT cells (expressing *IL1R1*), and C-FIB-OSMR^hi^ (expressing both *IL1B* and *IL1R1*). T cells can recruit inflammatory macrophages which may signal to injured PTs and C-FIB-OSMR^hi^ cells to sustain a fibro-immune environment poised for failed repair^73,74^. Further perpetuating this recruitment, both altered PTs and inflammatory fibroblasts (aPT2, C-FIB-OSMR^hi^) themselves expressed *CCL2* and co-localized with macrophages and dendritic cells that expressed *CCR2*, consistent with Visium stromal (6, 7) and PT (10, 11, 28) niches **(Figure 4h-j).** While an acute response by early inflammatory macrophage states (**Figure 2**), may establish an initial IL1B-IL1R1 mediated fibro-immune environment (**Figure 4k,l)** suppressing TGFβ-mediated myofibroblast differentiation ^75^, a subsequent transition to a fibro-cellular environment would be needed for tissue repair and resolution ^17^. The perpetuation of these signals and resultant states within the niche may lead to continued fibro-immune recruitment and a failure to resolve. Therefore, this work has identified the spatially organized cellular identities of these clinically important molecules, and their inter-cellular communications that may drive fibrotic progression.

### Regulation of recovery and failed repair by SOX4

Gene regulatory analyses and spatial cell state mapping identified SOX4 as a potential key player associated with injury states and niches. Compared to HNF4A in healthy tubules, inferred SOX4 activity localized to histologically-annotated tubulointerstitial injury in FFPE samples **(Figure 5a, b)**. Within the KPMP frozen Visium dataset, increased SOX4 activity was associated with non-recovery of AKI and progression of CKD **(Figure 5c)**. Velocity analysis of the SOX4 frPT and frTAL multiome trajectories indicated a key regulatory point in the mid-to-failed repair transition, with *in silico* knockout leading to trajectory disruption and acceleration to failed repair **(Figure 5d, ED Figure 11c)**. A subset of mid-repair cells also projected to the healthy trajectory, suggesting that SOX4 may hold a dimorphic regulatory role at a critical juncture of transitioning to failed repair or recovery.

To further support this, we performed siRNA knockdown of *SOX4* in human hTERT-converted PT cells (**Figure 5e-f**). This led to a partial recovery, with increased inferred activity of some healthy or early repair state TFs (THRB, HNF4G, KLF6) and the reduction of certain failed state TFs (RELB, EGR1, ELF3) (**Figure 5e**). However, we did find increased activity of NFKB1, including target genes *CCL2* and *VCAM1*, that have been associated with CKD progression ^5,18,76^. Consistently, there was an observed increase in the senescence / cell cycle arrest genes *CDKN1A* and *KHDRBS1* and decrease in pro-reparatory *GDF15* gene ^77^ after *SOX4* perturbation (**Figure 5f)**. These observations suggest that SOX4 may promote an early PT repair state following injury, while expression reduction in mid-repair states may help shift the balance towards a secretory senescence phenotype under conditions favoring NFKB1 and REL-mediated CCL2 expression. Examination of *CDKN1A* and *CCL2* promoter regions showed overlapping *SOX4*, *NFKB1* and *RELA/B* upstream binding sites, indicating potential cooperativity between these TFs to modulate *CCL2* (**Figure 5g, ED Figure 11d**)^78,79^. A SOX4/RELA binding site upstream of the promoter of *CCL2* (potential regulatory region 1) was well supported as a potential active enhancer given increased occupancy of H3K27ac and H3K4me1 histone marks in AKI and CKD diseased tissue. A *SOX4* binding site was also identified in the *GDF15* promoter near a pathogenic SNP associated with a decline in kidney function, and translational ribosomal profiling analysis of injured PT support that both Sox4 and Gdf15 proteins were translated **(ED Figure 11e-f)**. These results suggest coordinated activities of SOX4/NFKB1/REL or GDF15 may contribute to the divergence of epithelial repair trajectories towards a pro-senescent failed or a resolved state.

SOX4 activity subsequently re-emerges in failed repair (**Figure 3c**), potentially due to sustained TGFβ-SMAD3 signaling in kidney injury and fibrosis ^80^. This may generate an “on/on” type phenotype, as described for SOX9, to maintain both a progenitor-like and pro-fibrotic secretory cell state ^58^. Consistent with this, *SOX4-*positive cells showed a marked increase in abundance in CKD and were found co-localized with the senescent marker *CCL2* in injured tubules **(Figure 5h**). These co-positive cells neighbored dendritic cells (cDC2, target of *CCL2*^+^ cells), inflammatory fibroblasts (C-FIB-OSMR^hi^) and macrophages, consistent with niches described in **Figure 4**. By identifying histologically-defined disease-associated niches, our analyses provide insight into the TF activities and intercellular communications that may ultimately drive unresolved repair.

### Translation of molecular signatures to clinical biomarkers and therapeutic insights

To translate our atlas discoveries into clinically actionable insights, we identified secreted proteins from repair state signatures that could serve as early biomarkers of disease progression and therapeutic response. Our approach leveraged cell-type-specific expression patterns from altered epithelial, fibroblast, and immune states to prioritize candidates with the highest translational potential (**Figure 6a**). We systematically analyzed secreted repair state markers from proximal tubule, thick ascending limb, fibroblast, and macrophage lineages (**Supplementary Table 26**) for their association with clinical outcomes of AKI or CKD in independent patient cohorts (**Figure 6b-c**).

### Biomarkers of acute kidney injury and early progression

Analysis of the TRIBE-AKI cohort (784 cardiac surgery patients) revealed 27 plasma proteins that conferred greater odds of AKI (**see Methods**) in plasma samples (**Figure 6b**)^81^. These biomarkers fell into three distinct categories:

*Established kidney injury markers:* First, we identified proteins with previously established associations with kidney injury (B2M, CST3, IGFBP7), which served as positive controls and validated our approach, and markers confirmed in two additional AKI cohorts (ASSESS and NAIKID, **Supplemental Table 27**).

*Novel fibrosis-associated markers:* Second, we discovered several proteins linked to fibrotic processes: COL6A3 (may promote a profibrotic environment ^82–84^; NBL1 which has been implicated in glomerular pathology and ESKD; and FSTL1 and FSTL3 which are associated with glomerular diseases and eGFR decline ^85,86^.

*Fibro-Immune-epithelial interaction markers:* Third, we identified IL1R1, which our atlas mapped to aPT-fibro-immune niches (**Figure 4**) and was implicated in AKI severity ^72^. Notably, several biomarkers (COL15A1, COL18A1, PTPRS, PXDN, IDS and B4GALT1) represent novel candidates with limited prior associations with kidney disease yet potentially regulate ECM activities in AKI and its remodeling effector functions **(Figure 1b),** highlighting the discovery potential of our atlas-guided approach. B4GALT1 is a key enzyme that modifies N-glycan structures on glycoproteins by adding a stereospecific galactose to a N-acetylglucosamine residue. N-glycans are ubiquitous protein post-translational modifications, which are essential for the proper membrane localization of several essential component proteins (e.g., nephrin, podocin, and Crumbs2). Changes in their chemical structure have recently been associated with DKD. Using spatial N-glycomics analyses, we discovered increased abundance of galactose containing N-glycan containing glycoproteins in injured tubulo-interstitium supporting the transcriptomics findings of increased *B4GALT1* (**ED Figure 11g**).

### Biomarkers associated with chronic kidney disease progression

To identify biomarkers of disease progression, we analyzed plasma proteomics data from 418 participants in the BKBC cohort of which 115 progressed to ESKD during a median follow-up of 3.1 years (**Figure 6c, Supplementary Table 28**). This analysis revealed 82 unique differentially expressed proteins between progressors and non-progressors after multivariable adjustment at a significance level of p < 0.05. Pathway analysis revealed enrichment in ECM remodeling, neutrophil degranulation, metabolic dysfunction, and immune system activation. Importantly, 7 proteins were significantly associated with increased risk of ESKD after Bonferroni correction (**Supplementary Table 28**). Key discoveries included: PXDN (peroxidasin homolog), which was also associated with greater odds of AKI in the TRIBE-AKI cohort and found upregulated in multiple fibrosis models ^87^, suggesting a conserved role in progressive fibrosis; GM2A, which has been associated with AKI severity and failed recovery ^88^; SELENOM, that has been associated with progressive kidney function decline ^89,90^ (**Supplementary Table 27**).

### Therapeutic response monitoring and mechanistic insights

Demonstrating the clinical utility of our atlas for monitoring therapeutic response, *GDF15*, associated with AKI severity in both the TRIBE-AKI and NAIKID cohorts ^53^, showed expression in the TAL that was responsive to sodium-glucose cotransporter-2 inhibitor (SGLT2i) therapy in a cohort of individuals with youth-onset diabetes mellitus (**Figure 6d**) ^91^. This molecular response correlated with a transcriptomic signature consistent with tissue recovery, characterized by increased expression of canonical healthy TAL genes (*SLC12A1, WNK4, EGF, UMOD, and DEFB1*), cellular survival genes (*ADIRF and GPC3*), and reduction of the injury marker *HIF1A*. Our results demonstrate how our atlas enables real-time translation of tissue-level discoveries into early blood and urine markers that can monitor both disease progression and treatment response in clinical practice.

### Cellular sources and mechanistic validation

To definitively establish the cellular origins of these biomarkers and validate their disease associations, we integrated their expression patterns with histopathological features and atlas cell types and states (**Figure 6e**). This analysis confirmed their preferential association with fibrotic and inflammatory regions and demonstrated their origination from altered PTs, fibroblasts and immune cells, providing direct mechanistic links between tissue pathology and circulating biomarkers.

To elucidate the molecular mechanisms underlying disease progression, we performed differential gene expression analysis of KPMP participants with AKI who exhibited decline in eGFR at 18-month follow-up (progression) compared to participants who recovered (**see Methods**, **Figure 6f, Supplementary Table 29, 30).** Genes associated with AKI-to-CKD progression showed enrichment for regulatory targets of key transcription factors identified in our failed repair trajectories, including REL, NFKB1, and SOX4 (**Figure 6g**). Several genes associated with progressive decline in kidney function were identified as putative SOX4 targets, including *MID1* and *PXDN,* the latter of which was identified as a secreted marker in both the TRIBE and BKBC cohorts (**Figure 6b-c, h**). MID1 represents an E3 ubiquitin ligase upregulated in human DKD and murine fibrosis models ^92^ that may regulate STAT3 to modulate EMT and inflammation. Both MID1 and PXDN proteins were immunohistochemically confirmed to localize to PTs, with PXDN also localizing to interstitial fibroblasts in AKI (**Figure 6i).**

The expression patterns of *PXDN*, which encodes a collagen IV cross-linking enzyme that increases ECM stiffness ^87,93^ and *SOX4* were synchronous across transitions from early injury to failed repair states in both human and mouse kidney atlases (**Figure 6j-k**). Consistent with a direct regulatory connection, the *PXDN* gene showed four accessible SOX4 binding sites with active enhancer (H3K27ac/H3K4me1) histone marks through CUT&RUN analysis in participants with AKI or CKD (**ED Figure 11h**). This provides a direct molecular connection between SOX4 activity and the ECM-remodeling and scar-promoting activities mediated by PXDN. These findings demonstrate that our atlas not only identifies clinically relevant biomarkers but also reveals the underlying regulatory mechanisms driving disease progression. This integrated approach supports the concept that select soluble factors secreted during the early-to-mid injury states are predictive of kidney function decline through specific transcriptional programs. The HKAv2 atlas thus provides a comprehensive framework that identified the regulatory networks governing the underlying biomarkers but also the mechanisms of injury progression and the cellular sources, enabling precision medicine approaches for kidney disease.

## Discussion

HKAv2 leverages an immense dataset of over 1.7 million nuclei and cells, with rigorous spatial validation from five ST datasets, multiplexed immunofluorescence, and spatial metabolomics to define temporally regulated cell states and their relationship to injury recovery or disease progression. A key aspect of HKAv2 is the multiomic data culled from a prospective cohort of altruistic kidney tissue donors, including those with common presentations of CKD and AKI that may not be biopsied. This enabled delineation of the time course of acute and chronic injury, defining mechanisms that are most important for kidney diseases with the largest public health burden, including early stages that are amenable to primary and secondary prevention. This was further strengthened by expert adjudication of clinical diagnoses and detailed pathology descriptor scoring, enabling the association of 159 cell states with clinical and pathological variables. Integration of the HKAv2 with a simultaneously constructed multiome mouse atlas clearly identified the order of appearance of specific injured tubular and interstitial cell states from early injury to failed repair. Spatial technologies mapped these states into discreet AKI/CKD associated niches that identify cell-cell interactions in various stages of injury. HKAv2 helped identify specific inflammatory macrophages, inflammatory fibroblasts and dendritic cells in areas of injured tubules defining key signaling mechanisms contributing to tubulointerstitial injury. This has highlighted a role for SOX4 in early injury recovery by inhibiting senescence or in failed repair progression by promoting a profibrotic and pro-inflammatory niche through cooperation with REL/NFKB (**Figure 6l**). Importantly, HKAv2 identifies links between the mechanistic, clinical and histopathological domains, demonstrating interdisciplinary synergy and clinical utility of adding spatiotemporally defined molecular data. Using secreted markers grounded to HKAv2 spatial and molecular insights, we infer the dominant cellular states underlying progression (from early to mid to failed repair) and identify markers that may indicate prognosis for AKI recovery, CKD progression, or that may improve therapeutic selection, as demonstrated by reversion to recovery states with SGLT2i therapy. From this we identify a mechanism of SOX4-regulated secretion of PXDN from injured proximal tubular cells that may predict long-term AKI progression to CKD. The detailed pathways mapped across cell states, clinical conditions and druggable targets further provide novelty in designing therapeutic strategies or repurposing existing drugs to alter the course of kidney disease progression.

## Extended Data Figure Legends

**ED Figure 1.**
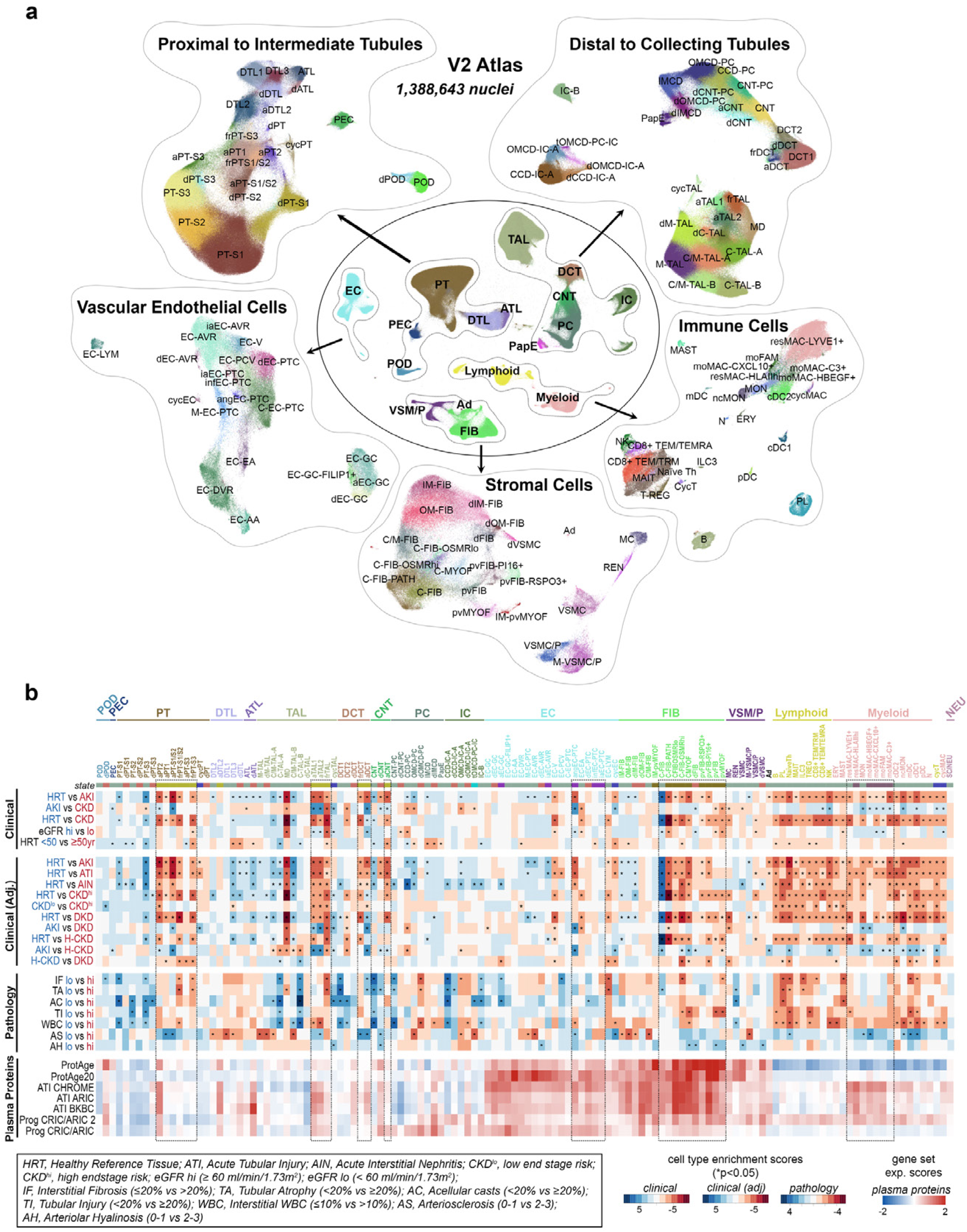
Human Kidney Atlas (HKA)v2. **a**. UMAP embeddings showing broad subclasses (center plot) and the associated groupings that were sub-clustered independently for a high-level cell type resolution (outer plots). **b**. Heatmaps showing cell type enrichments as t statistics for the comparison of cell type abundance between two different patient groupings. Asterix indicates p values less than 0.05. Comparisons were done for broad and adjudicated (KPMP) clinical categories as well as for groupings based on pathology descriptor scoring. Only biopsy samples (excluding nephrectomy and deceased donor) were used for all categories except age associations. Number of patients for clinical: HRT (41), AKI (38), CKD (73), eGFRhi (69), eGFRlo (56), HRT <50 (34), HRT ≥50 (38). Number of patients for adjudicated clinical: HRT (41), AKI (28), ATI (20), AIN (11), CKDhi (26), CKDlo (9), DKD (31), H-CKD (24). Number of patients for pathology: IF lo (48), IF hi (33), TA lo (45), TA hi (36), AC lo (72), AC hi (8), TI lo (38), TI hi (43), WBC lo (45), WBC hi (36), AS lo (45), AS hi (34), AH lo (49), AH hi (32). Bottom panel represents a heatmap of averaged scaled expression scores for select gene sets associated plasma proteins that were linked to aging ^95^, acute tubular injury (ATI) ^53^ or CKD progression ^96^. Source data provided.

**ED Figure 2.**
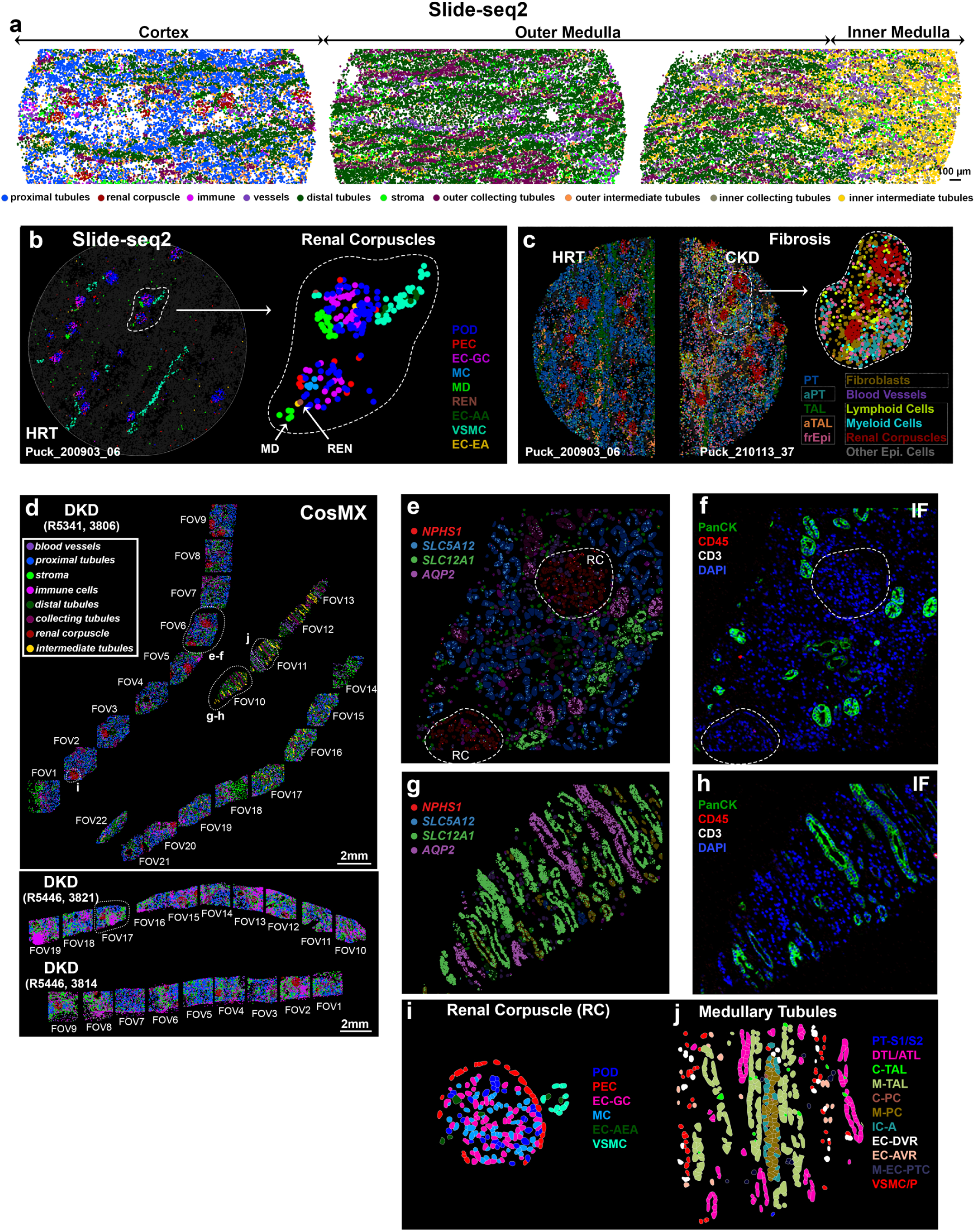
Spatially mapped HKAv2 cell types. **a**. Slide-seq2 cell types grouped by structure and spatially localized along the cortico-medullary axis. **b**. Left, a single Slide-seq HRT puck showing renal corpuscle associated cell types. Right, enlarged area of that shown in the left panel for the same cell types and highlighting correct spatial localization of rare MD and REN cells. Tissue pucks are 3mm in diameter. **c**. Slide-seq2 tissue pucks (halved) associated with HRT (left) and CKD (right), showing enriched mapping for adaptive state epithelial cells (aPT, aTAL, failed repair PT and TAL epithelial cells or frEpi) as well as general fibroblast, lymphoid and myeloid cells in the diseased tissue. **d**. CosMx cell types grouped by structure and spatially localized to different cortico-medullary fields of view (FOV) for three different diabetic kidney disease (DKD) biopsies. **e**. Cortical FOV indicated in (d) showing spatial mapping of cell type specific transcripts. Renal corpuscles (RC) are indicated. **f**. Same FOV as in (**e**) showing protein immunofluorescent staining for epithelial pan cytokeratin (panCK) that highlighted collecting ducts, lymphoid CD45 and T-cell CD3 antibodies. Minimal staining of the latter two is consistent with the low immune cell infiltration found within this FOV. **g**. Medullary FOV indicated in (**d**) showing spatial mapping of cell type specific transcripts. **h**. Immunofluorescence staining as in (**f**) and highlighting medullary collecting ducts. **i**. Enlarged area shown in (**d**) highlighting RC cell types. **j**. Enlarged area shown in (**d**) highlighting medullary cell types.

**ED Figure 3.**
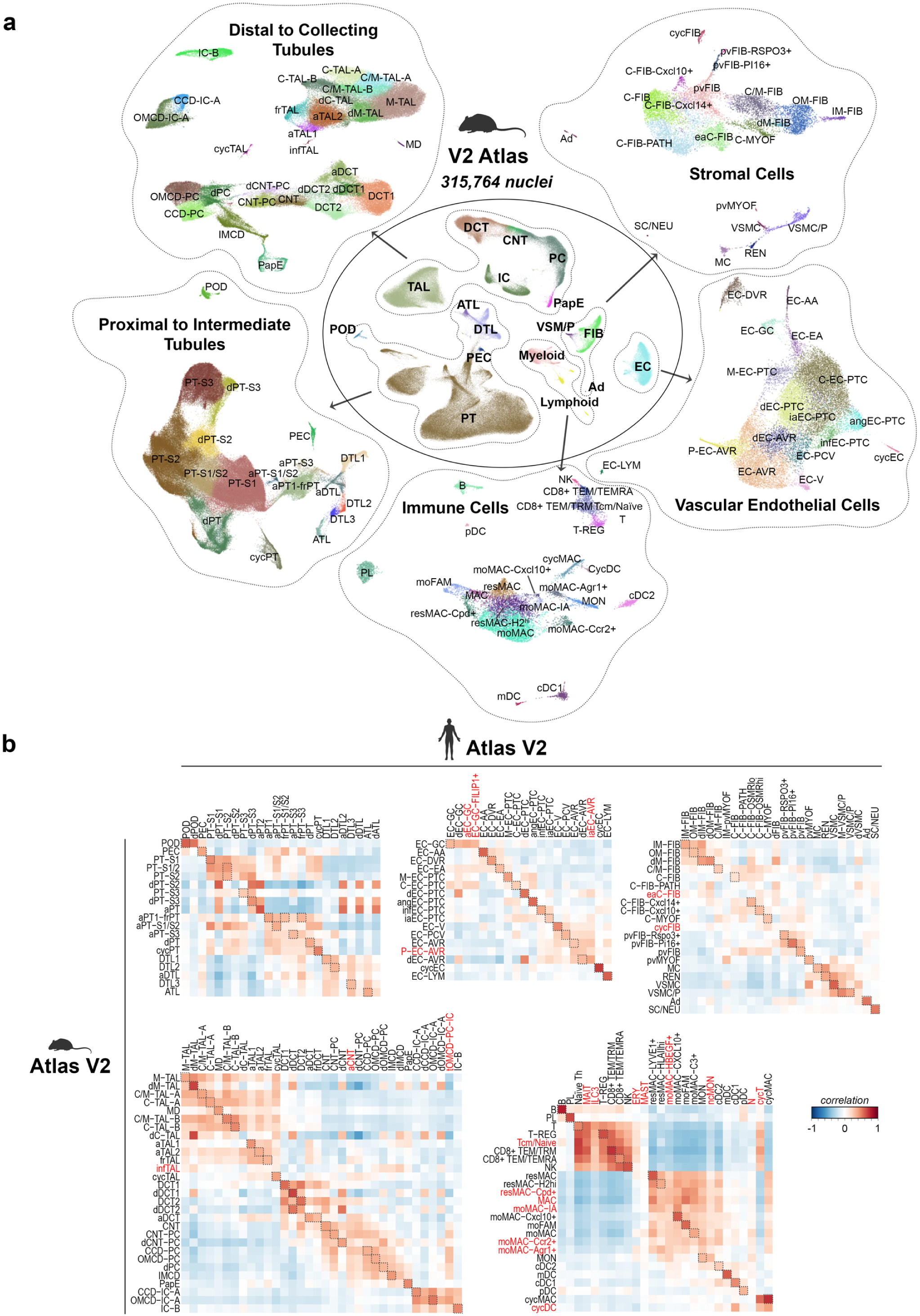
Mouse Kidney Atlas. **a**. UMAP embeddings showing broad subclasses (center plot) and the associated groupings that were sub-clustered independently for a high-level cell type resolution (outer plots). **b**. Heatmaps showing correlation values for human and mouse cell types. Boxes indicate potential conserved cell types or states, red text indicates subtypes found in only one species. Source data provided.

**ED Figure 4.**
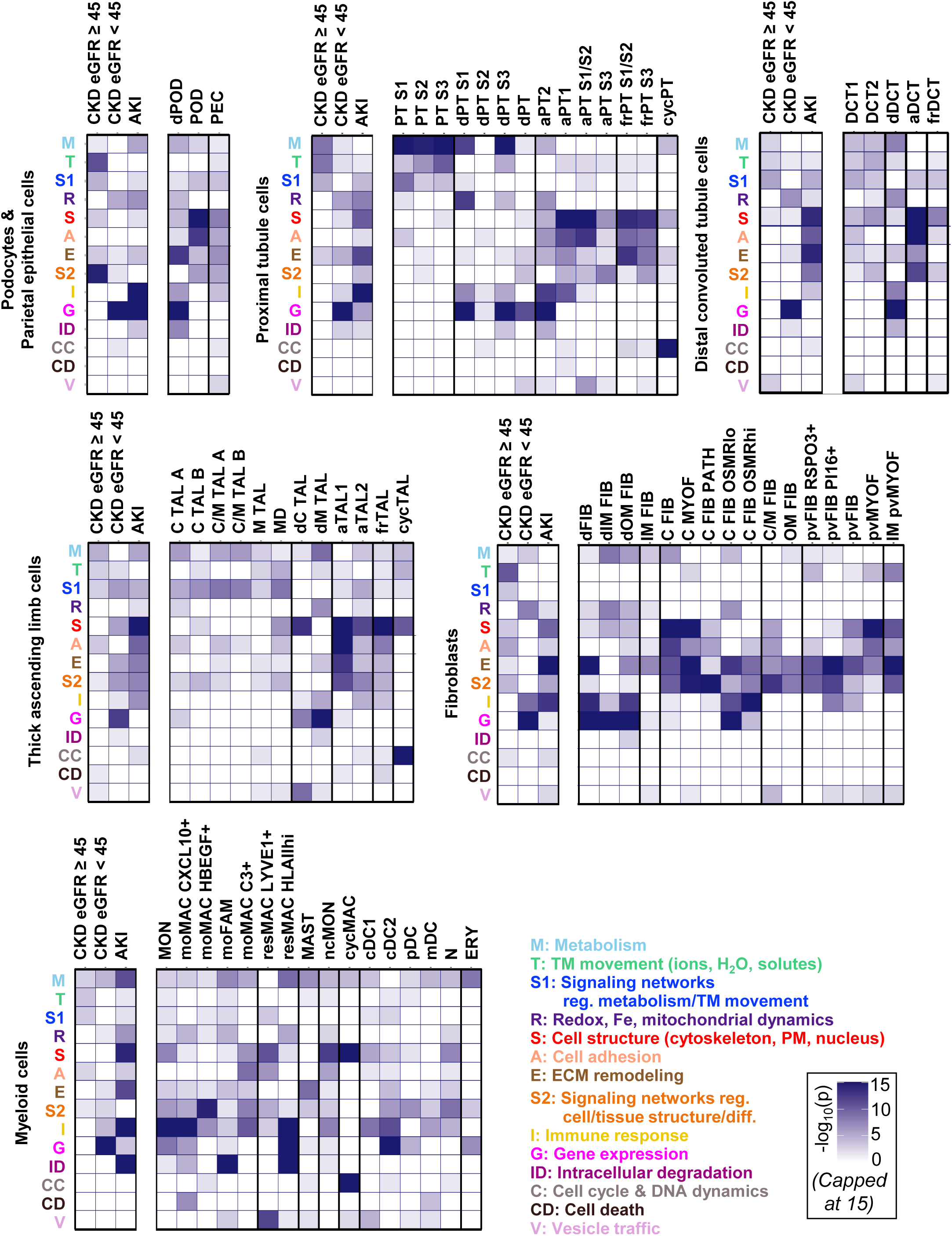
Cell state and condition-selective whole cell functions. For each cell type individually, we calculated condition- or cell subtype/state-selective marker genes by comparing gene expression profiles in all cells of a given cell type that were annotated to the same clinical condition or cell subtype versus those cells of the same cell type that were annotated to the other clinical conditions or subtypes. Note that in the first case we only considered cells annotated to the three clinical conditions, while ignoring cells annotated to any other conditions. Condition- or subtype-selective marker genes were then submitted to pathway enrichment analysis using MBCO level-3 subcellular processes, followed by expert-curation and annotation of predicted pathways (p-value ≤ 0.05) to shown 14 whole cell functions. −log10(p-values) of pathways annotated to the same whole cell function were summed up, color-coded and visualized in heatmaps. For subtype counts in the different conditions, see **ED Figure 1B**.

**ED Figure 5.**
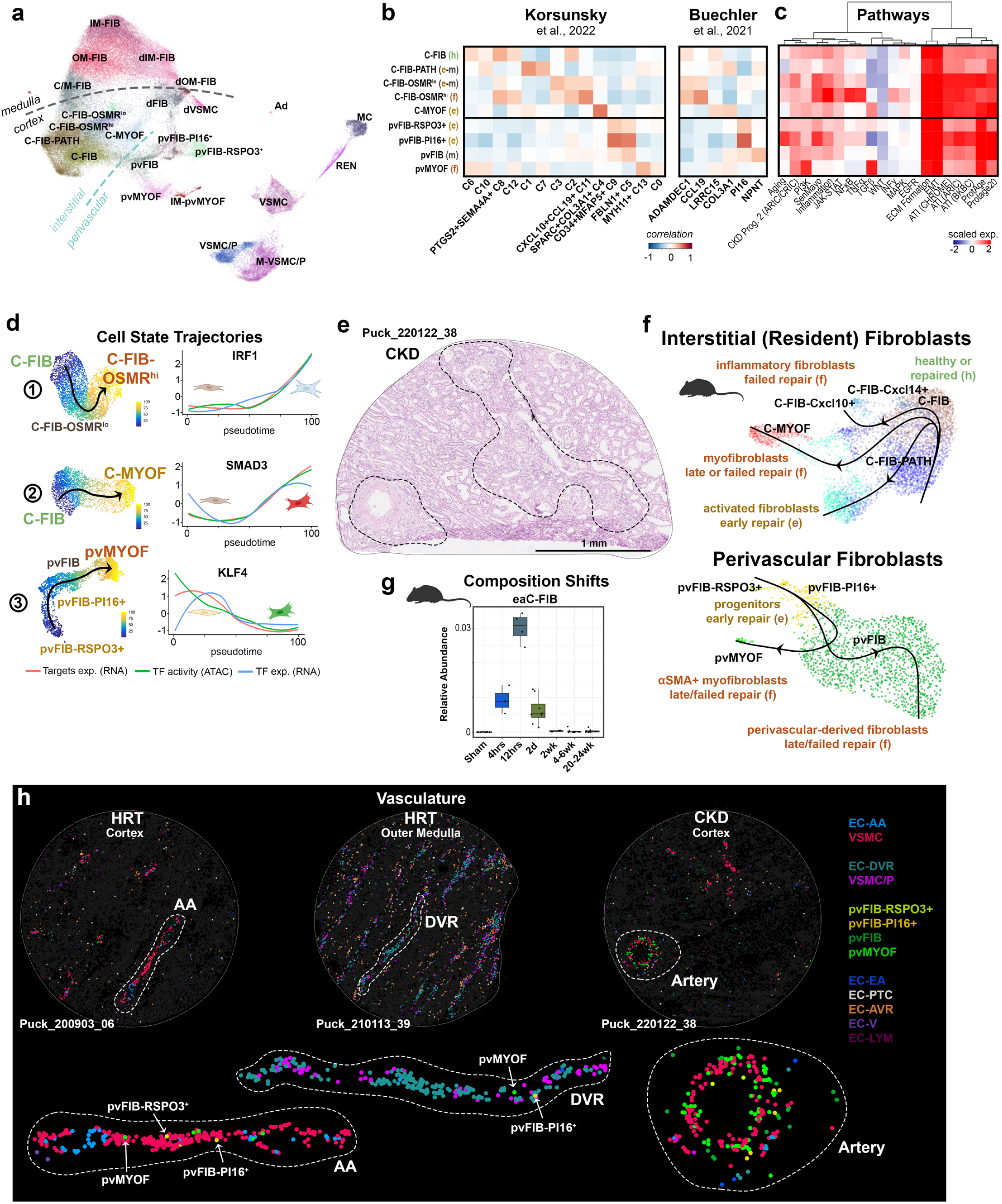
Distinct and conserved fibroblast subtypes. **a**. UMAP embeddings showing fibroblasts subtypes spanning the corticomedullary axis and with resolution of cortical interstitial and perivascular subtypes. **b**. Heatmaps showing correlation of cortical fibroblast subtypes with subtypes found in two separate cross-tissue studies ^8,15^. **c**. Heatmap of averaged scaled expression scores for signaling pathways or functional gene sets. **d**. Left, UMAP embeddings for individual trajectories shown in **Figure 2a** colored by estimated pseudotime. Right, scaled target gene expression (RNA), binding site activity (ATAC) and gene expression (RNA) levels for select TFs as a function of pseudotime. **e**. Histology of tissue region shown in **Figure 2d**. **f**. UMAP embedding of mouse data showing interstitial (resident) and perivascular fibroblasts and their associated Slingshot predicted trajectories. **g**. Bar plot showing relative abundance for early activated fibroblasts within mouse samples collected at different time points from IRI and compared to sham controls. **h**. Slide-seq2 tissue pucks showing localization of vascular-associated cell types. Lower panels represent enlarged regions that are indicated in the upper panels, highlighting perivascular fibroblast subtype localizations within the afferent arterioles (AA) and descending vasa recta (DVR) of HRTs and expansion around small arteries (and in interstitial spaces) in CKD tissue. Tissue pucks are 3mm in diameter. Source data provided.

**ED Figure 6.**
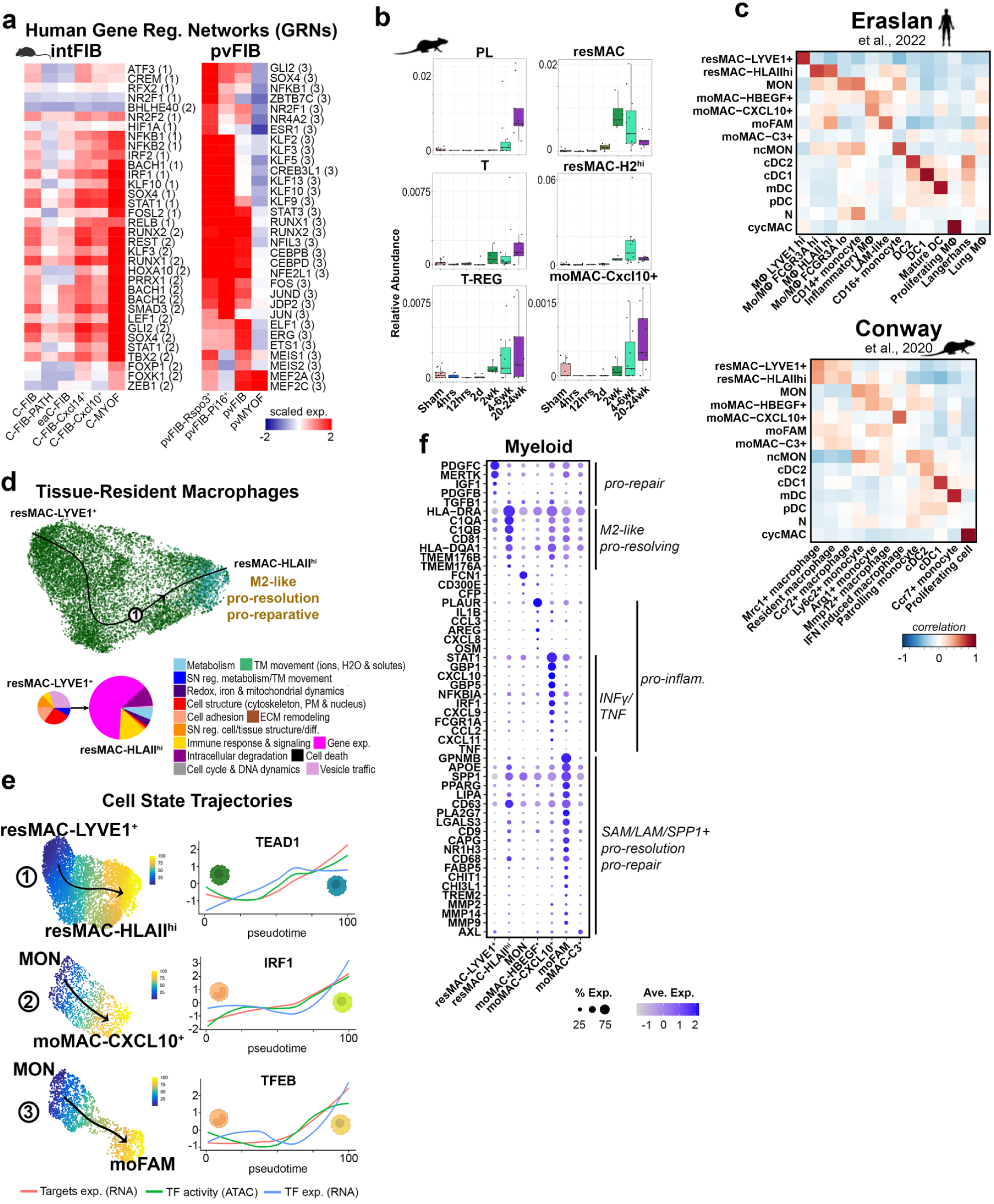
Fibroblast and myeloid signatures associated with injury and disease. **a**. Heatmap of scaled expression scores in mouse fibroblast subtypes for human TF target genes associated with GRNs built from trajectories shown in **Figure 2a**. Numbers indicate the specific trajectory used to derive the target gene set. **b.** Bar plots showing relative abundance for different immune subtypes within each mouse sample collected at different time points from IRI and compared to sham controls. **c**. Heatmaps showing correlation of human myeloid cell subtypes with subtypes found in a cross-tissue profiling study ^11^ and a murine reversible unilateral ureteric obstruction model^9^. **d**. UMAP embedding showing tissue-resident macrophage subtypes and their associated Slingshot predicted trajectories. Pie charts represent overall whole cell functions predicted for resident macrophages (sum of all −log10(p) for resMAC HLAIIhi: ∼149). **e**. Left, UMAP embeddings for individual trajectories shown in **Figure 2g** and (**d**) colored by estimated pseudotime. Right, scaled target gene expression (RNA), binding site activity (ATAC) and gene expression (RNA) levels for select TFs as a function of pseudotime. **f**. Dotplot for human myeloid cell subtypes showing average expression values for select marker genes. Source data provided.

**ED Figure 7.**
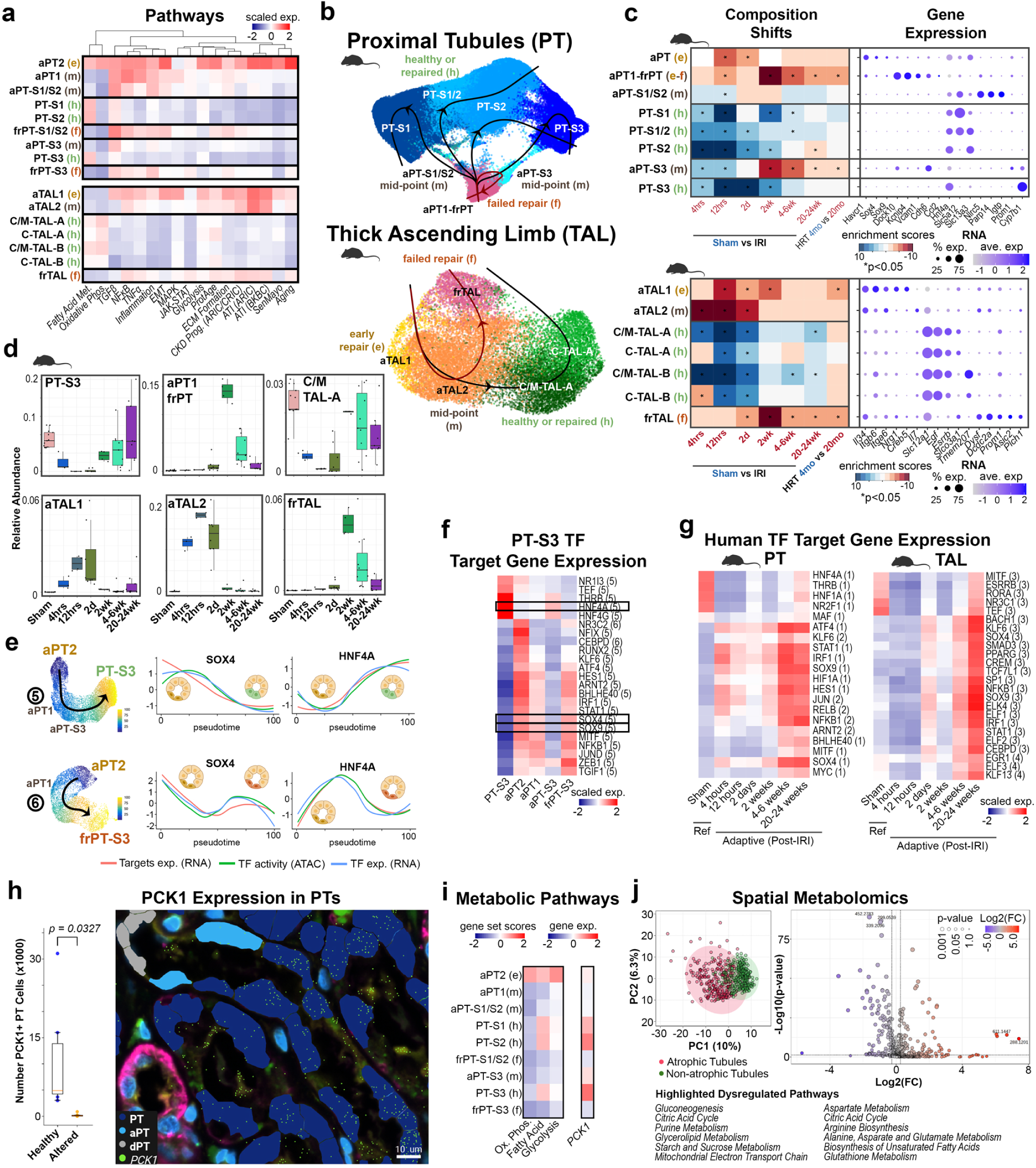
Resolving and unresolved epithelial repair states. **a**. Heatmap of averaged scaled expression scores for signaling pathways or functional gene sets. **b**. UMAP embedding of mouse data showing proximal tubule (PT) and thick ascending limb (TAL) states and associated Slingshot predicted trajectories. **c**. Left, heatmaps for mouse data showing cell type enrichments as t statistics for the comparison of cell type abundance between two different conditions (Sham versus different time points from IRI, or 4 month- versus 20 month-old healthy reference tissues (HRT). Asterisk indicates p values less than 0.05. Middle, dotplot showing average expression values for select marker genes. Right, dotplot showing average binding site accessibilities for select TFs. **d**. Bar plots showing relative abundance for different epithelial repair states within each mouse sample collected at different time points from IRI and compared to sham controls. **e**. Left, UMAP embeddings for individual PT-S3-associated trajectories (human) shown in **Figure 3a** and colored by estimated pseudotime. Right, scaled target gene expression (RNA), binding site activity (ATAC) and gene expression (RNA) levels for select TFs as a function of pseudotime. **f**. Heatmap of scaled expression scores for TF target genes associated with GRNs built from trajectories shown in **Figure 3a** and (**e**). Numbers indicate the specific trajectory used to derive the target gene set. **g**. Heatmap of scaled expression scores in mouse PT and TAL subtypes for human TF target genes associated with GRNs built from trajectories shown in **Figure 3a**. Numbers indicate the specific trajectory used to derive the target gene set. **h.** 10X xenium data showing enrichment of *PCK1* in PT compared to aPT and dPT in the same tubule, scale = 10µm. Bar plots show quantification. i. Heatmap of scaled expression scores for metabolic pathway gene sets or *PCK1*. **k.** Untargeted analysis of spatial metabolomics in atrophic and non-atrophic tubules (9 CKD biopsies). From each sample, 30 atrophic and 30 non-atrophic tubules were selected, for a total of 270 atrophic and 270 non-atrophic tubules. Right: PCA plot. Left: Volcano plots using a 1.2-fold change cutoff and FDR-adjusted P value < 0.05. Highlighted dysregulated pathways summarized from enrichment analysis with KEGG and pathway analysis with the SMPDB database using MetaboAnalyst 6.0. Created in BioRender. Jain, S. (2025) https://BioRender.com/dgtgsay. Source data provided.

**ED Figure 8.**
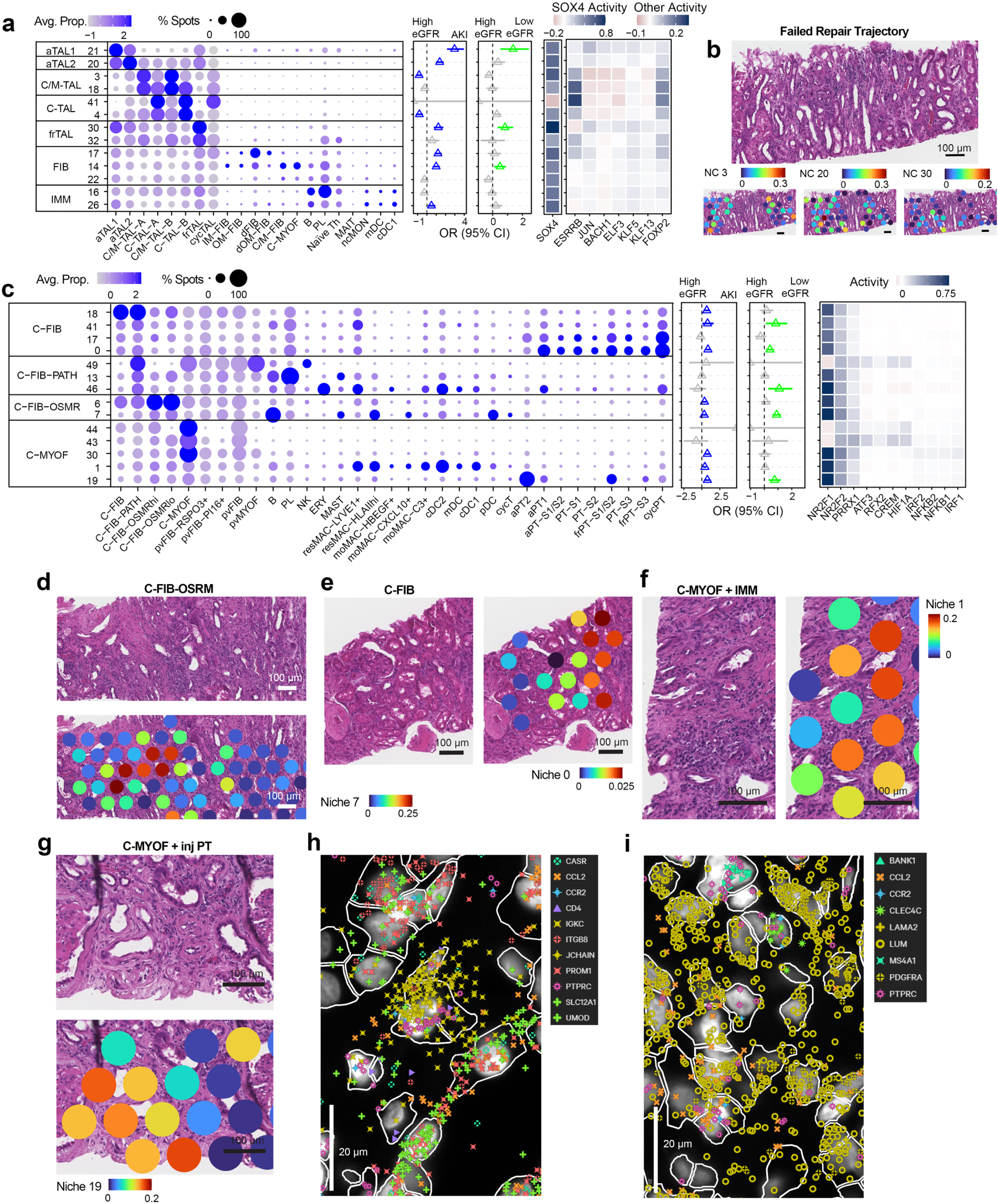
Visium niches. **a.** Characterization of selected TAL niches with cell type distribution, transcription factor network activities, and association with clinical status, with log odds ratio and confidence interval. **b.** Examples of mapping of reference, adaptive, and failed repair TAL niches on FFPE histology. **c.** Characterization of Fib niches along the trajectory from C-FIB to C-FIB-ORSMR niches with cell type distribution, transcription factor network activities, and association with clinical status, with log odds ratio and confidence interval. **d.** A region with fibrosis and heavy immune infiltration with high mapping of a C-FIB-OSRM niche 7. **e.** A region of cortical tubulointerstitium with high mapping of a C-FIB niche 0 over a region fibrosis and more modest chronic tubulointerstitial nephritis compared to **d**. **f.** C-MYOF niche 1 localizes to a region of severe intertsitial fibrosis, tubular atrophy, and chronic tubulointerstitial nephritis. **g.** C-MYOF niche 19 localizes to a fibro-immune or fibrotic areas on FFPE histology. **h**. Xenium example of a niche with injured TAL expressing *CCL2* with Stromal and Immune cells expressing *CCR2* (consistent with Visium TAL niche 16). **i.** Xenium example of a niche with C-FIB-OSMR surrounded by B and pDC cells (consistent with Visium FIB niche 7). Source data provided.

**ED Figure 9.**
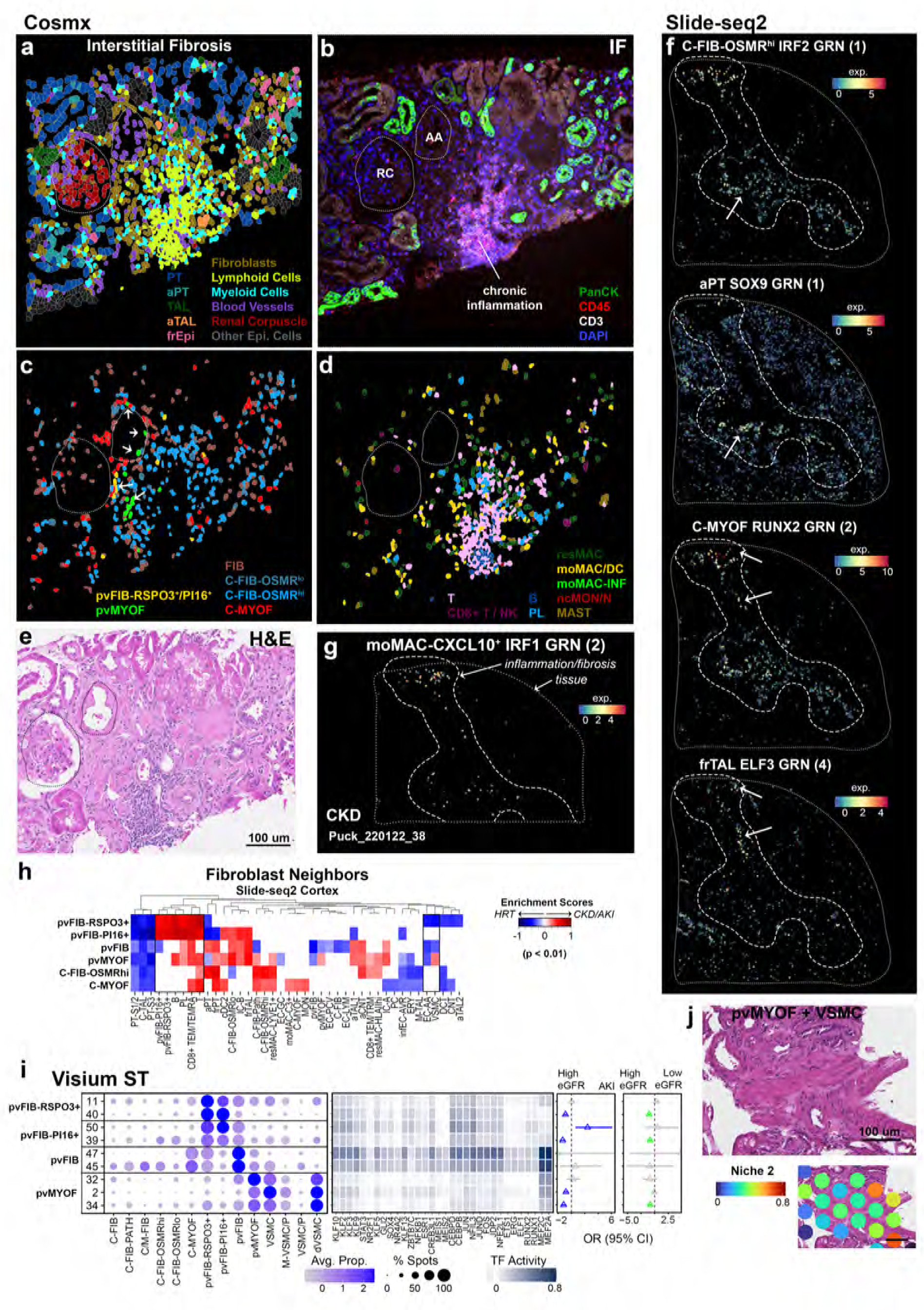
Spatial niches of epithelial, fibroblast and immune cell states. **a**-**e**. CosMx cortical FOV indicated in **ED Figure 2d. a**. Spatial mapping of specific cell types and structures. Renal corpuscles (RC) are indicated. **b**. Protein immunofluorescent staining for epithelial pan cytokeratin (panCK) highlighted collecting ducts and distal tubules, lymphoid CD45 and T-cell CD3. **c**. Spatial mapping of fibroblast subtypes. **d**. Spatial mapping of immune subtypes. **e**. Corresponding histology image. **f**. Slide-seq2 subset region of tissue in **Figure 2e** showing TF GRN target gene set expression scores within the associated broad cell types. **g**. Slide-seq2 tissue sub-region of that found in **Figure 2ed** showing TF GRN target gene set expression scores within macrophage cell types. **h.** Heatmap of enrichment scores for fibroblast subtype neighbors (Slide-seq2 cortical or cortico-medullary tissues) that show significant (p < 0.01) difference in enrichment scores between HRT and diseased (CKD/AKI) tissues. **i.** Visium ST characterization of pvFIB niches with cell type distribution, transcription factor network activities, and association with clinical status, with log odds ratio and confidence interval. j. Example of pvMYOF niche 2 mapped to FFPE histology. Source data provided.

**ED Figure 10.**
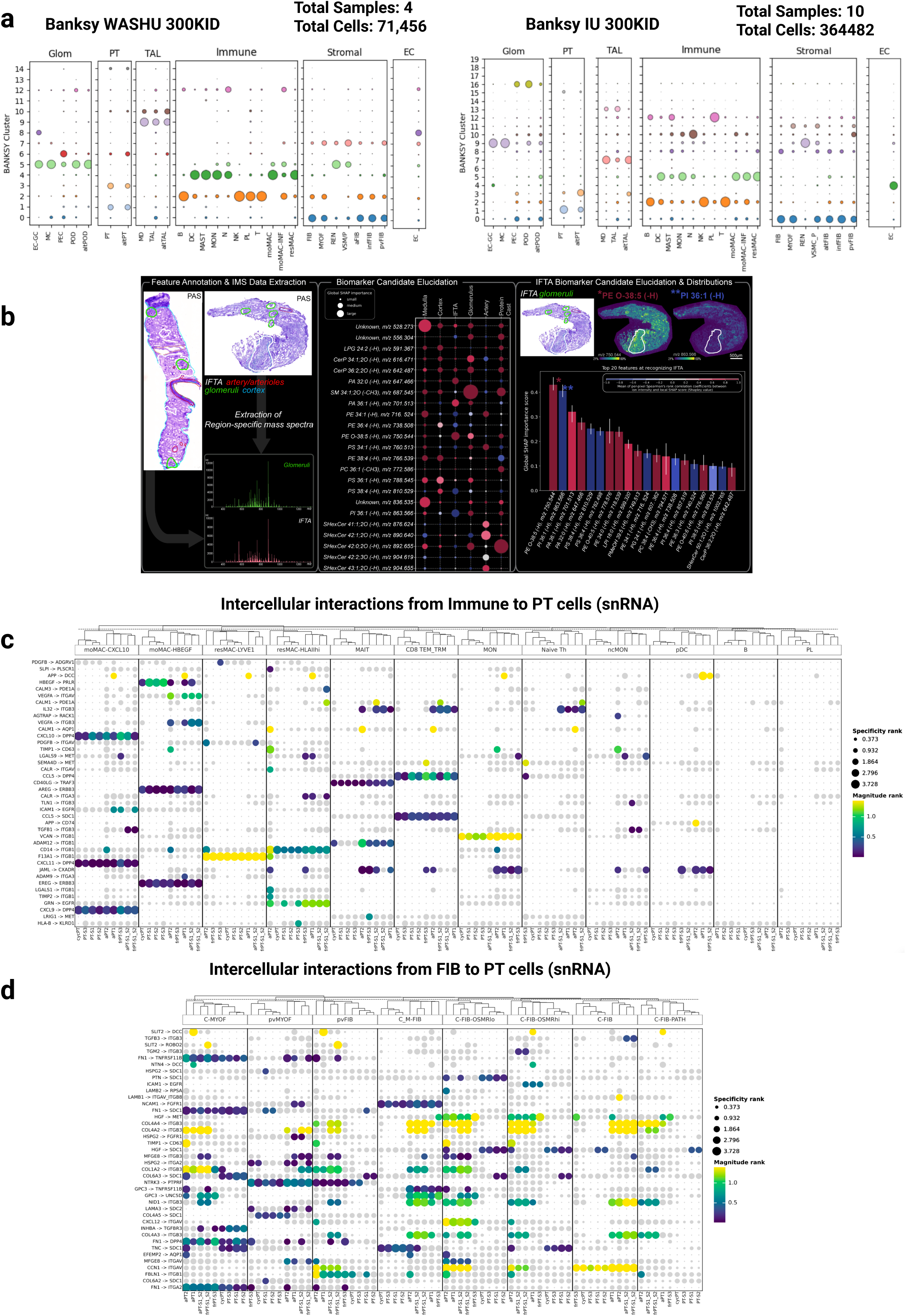
Spatial niches and L-R analyses. **a**. Xenium niche analysis on WU 4 diabetic nephropathy kidney biopsy. **b**. Xenium niche analysis on IU 13 diabetic nephropathy kidney biopsies. **c**. Lipids Panel shows PAS stains of 2 biopsies, with manual annotations of features – glomeruli (green), IFTA (white), artery/arteriole (red), and cortex (cyan). The annotations were used to extract IMS pixels and generate average mass spectra of regions of interest (Glomeruli - green; IFTA – pink). The SHAP bubble plot highlights the top 20 biomarker candidates (rows) for each histological feature (column). The marker’s size corresponds to the global SHAP importance score; the marker color corresponds to the Spearman rank-order correlation: positive (red) and negative (blue). The bar graph highlights the top 20 biomarker candidates for IFTA, with ion images of the top positively (red) and negatively (blue) correlated ions seen above the bar graph. **d, e**. putative ligand receptor (L-R) interactions between immune or fibroblast with PT cells ranked by how variable they are across PT states. Non-specific interactions are in grey. Color bars are –log10-transformed. Created in BioRender. Jain, S. (2025) https://BioRender.com/qu690b3.

**ED Figure 11.**
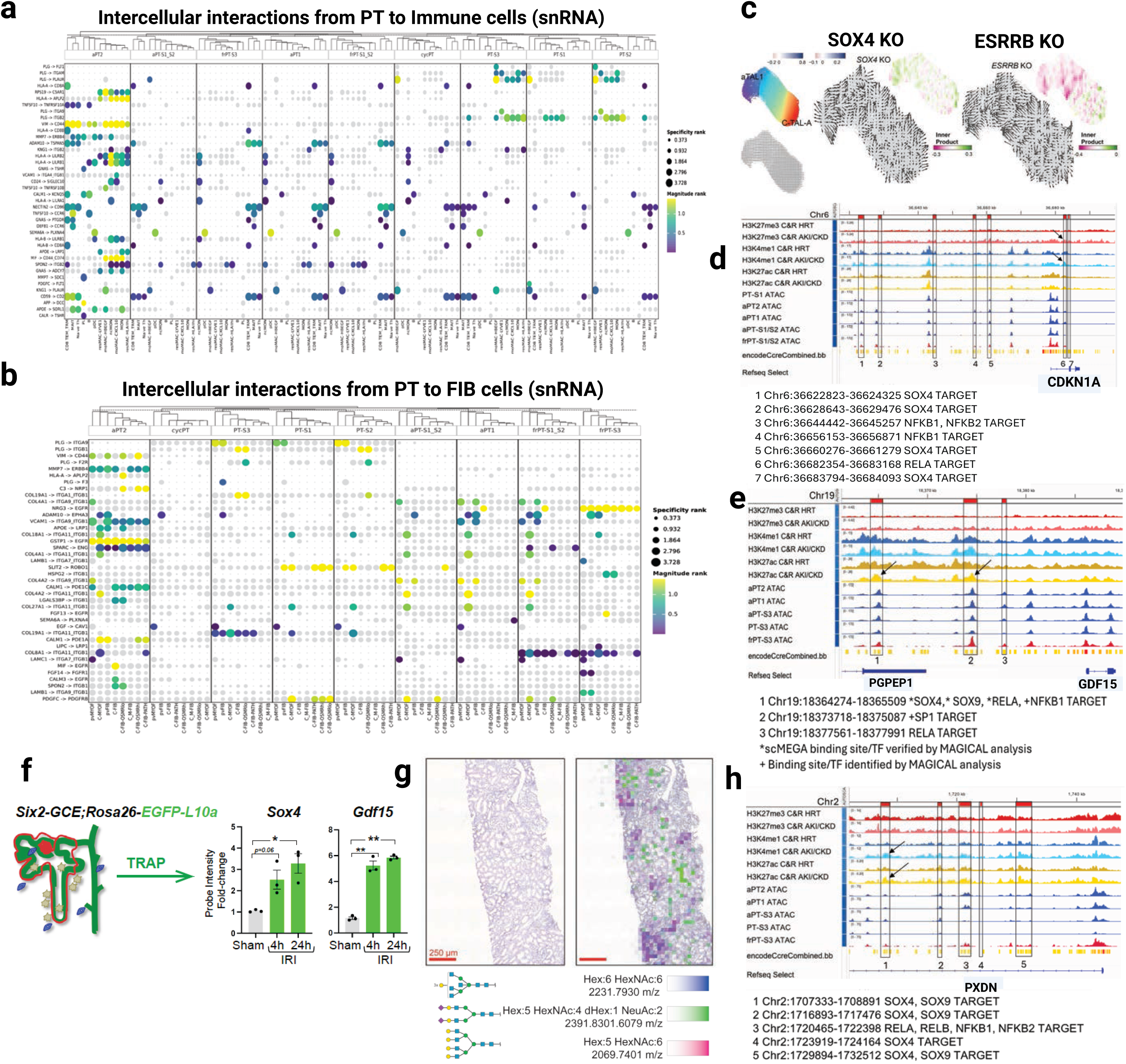
Intracellular signaling and gene regulatory activities. **a, b**. L-R analysis using Liana+ between PT-immune and PT-fibroblast. Non-specific interactions are in grey. Color bars are –log10-transformed. **c.** Flow in the successful TAL repair trajectory is disrupted after *in silico* knock-out of *SOX4* and *ESRRB.* **d**. Genomic region of *CDKN1A* visualized with Cut&Run tracks for histone marks (H3K27ac, H3K4me1, and H3K27me3) in healthy reference and AKI/CKD kidney tissue, and pseudo bulk ATAC coverage tracks for PT trajectories. scMEGA analysis identified candidate regions regulated by potential transcription factors (SOX4, RELA, NFKB1) upstream (regulatory regions 1-5) and downstream (regions 6, 7) of the *CDKN1A* gene. Intronic regulatory region 1 has an increased chromatin activity in the aPT1, aPT2, and frPT-S1/S2 cell populations of the trajectory compared to healthy PT-S1 cells and it correlates with the increased occupancy of H3K4me1 and H3K27me3, as well as binding of H3K27ac, indicating an active regulatory region. **e.** Genomic region of *GDF15* visualized with Cut&Run tracks for histone marks (H3K27ac, H3K4me1, and H3K27me3) in healthy reference and AKI/CKD kidney tissue, and pseudo bulk ATAC coverage tracks for PT trajectories. scMEGA analysis identified candidate regulatory regions regulated by transcription factors (SOX4, SOX9, RELA, NFKB1, SP1) upstream of the *GDF15* promoter. Upstream regulatory region 1 has increased open chromatin in the aPT cell populations of the trajectory compared to healthy PT-S3 cells that correlates with increased occupancy of active histone mark H3K27ac. This regulatory region was identified by both scMEGA and MAGICAL analysis and has binding sites for SOX4, SOX9, RELA, and NFKB1. **f.** Translating Ribosome Affinity Purification (TRAP) microarray analysis shows *Sox4* was upregulated in the nephron epithelia early after IRI-induced AKI. *Sox4* and *Gdf15* expression in *Six2-Tet-GFP-Cre;Rosa26-EGFP-L10a–*labeled nephrons at 4 h and 24 h post-IRI vs. sham. Mean ± SEM, n = 3/group. **g.** N-glycation analysis related to *B4GALT1* expression. Example Hematoxylin&Eosin-stained biopsy tissue section from a patient with AKI (left) with overlayed ion images of galactose containing N-glycans (right), created by expression of B4GALT1. **h**. Genomic region of *PXDN* visualized with Cut&Run tracks for histone marks (H3K27ac, H3K4me1, and H3K27me3) in healthy reference and AKI/CKD kidney tissue, and pseudo bulk ATAC coverage tracks for PT trajectories. Candidate regulatory regions (**Supplementary Table 25**) regulated by transcription factors (SOX4, SOX9, NFKB1, NFKB2, RELA, RELB) in the intronic region of *PXDN*. Increased open chromatin was observed in adaptive PT compared to healthy PT-S3 cells corresponding with active histone marks (H3K27ac and H3K4me1) (arrows). Created in BioRender. Jain, S. (2025) https://BioRender.com/m5rjwdo. Source data provided.

## Source Data

**Source data Supplementary DataTables:**

Supplementary Table 1. Summary of single-nucleus omic experiments

Supplementary Table 2. Summary of single-cell omic experiments

Supplementary Table 3. Summary of Visium spatial transcriptomic experiments

Supplementary Table 4. Summary of Slide-seq2 spatial transcriptomic experiments

Supplementary Table 5. Summary of CosMx spatial transcriptomic experiments

Supplementary Table 6. Summary of Xenium spatial transcriptomic experiments

Supplementary Table 7. Summary of MxIF/CODEX spatial imaging experiments

Supplementary Table 8. Summary of spatial metabolomic experiments

Supplementary Table 9. Summary of Cut&RUN experiments

Supplementary Table 10. Single nucleus cluster annotations

Supplementary Table 11. Single nucleus cluster metrics

Supplementary Table 12. Single nucleus cell type marker genes

Supplementary Table 13. Slide-seq2 cell type annotations.

Supplementary Table 14. CosMx cell type annotations.

Supplementary Table 15. Mouse omic experiments.

Supplementary Table 16. Mouse Single nucleus cluster annotations.

Supplementary Table 17. Mouse single nucleus cluster metrics

Supplementary Table 18. Single cell cluster marker genes

Supplementary Table 19. Single cell cluster metrics

Supplementary Table 20. Pathway analysis on AKI, CKD and cell states

Supplementary Table 21. Transcription factor networks for cell state trajectories

Supplementary Table 22. scMega defined transcription factor target genes

Supplementary Table 23. Gene sets associated with pathways

Supplementary Table 24. Disease-associated GWAS variants using MAGICAL

Supplementary Table 24. Summary of Cut&Run Experiments

Supplementary Table 25. PT transcription factors from scMEGA_MAGICAL analysis

Supplementary Table 26. Cell state marker genes associated with secreted proteins

Supplementary Table 27. Soluble protein markers significant in AKI severity and progression

Supplementary Table 28. Soluble protein markers significant in progression to ESKD in BKBC

Supplementary Table 29. KPMP AKI participants that recovered or progressed to CKD

Supplementary Table 30. Genes differentially expressed in AKI recovery or progression

**Source data Figure panels**

## Methods

### Statistics and Reproducibility

For immunofluorescence validation studies, commercially available antibodies were used; The manuscript leverages heavily of multimodal orthogonal validations with many particpant tissue interrogated by more than one technology. In some cases the same tissue block was used to generate multimodal data (**Figure 1b and source data)**. There is extensive validation where spatial transcriptomic annotations revealed similar marker gene expressions in snCv3/scCv3, multiome, as well as spatial localization which corresponded with histologically validated Visium spot mapping, SlideSeq2, Xenium, CosMx, MxIF and spatial metabolomics, and also where available supported by published literature. This heterogeneous sampling approach, the broad spectrum of patient samples representing tissue from various sources ensured cell type discovery while minimizing assay dependent biases or artifacts encountered when using different sources of kidney tissue. We recognize that the heterogeneity of sample sources for several technologies is a potential limitation due logistics and limited patient biopsy material.

Statistical approaches are built in the high throughput omics analysis and additionally listed in methods section and figure legends where graphical data are presented.

### Ethical Compliance

*Human studies.* We have complied with all ethical regulations related to this study. Experiments on human samples followed all relevant guidelines and regulations. Human samples (**Supplementary Tables 1-9**) collected as part of the Kidney Precision Medicine Project (KPMP) consortium (KPMP.org) were obtained with informed consent and approved under a protocol by the KPMP single IRB of the University of Washington Institutional Review Board (IRB#20190213). Samples as part of the Human Biomolecular Atlas Program (HuBMAP) consortium were collected by the Kidney Translational Research Center (KTRC) under a protocol approved by the Washington University Institutional Review Board (IRB #201102312). Informed consent was obtained for the use of data and samples for all participants at Washington University, including living patients undergoing partial or total nephrectomy or from discarded deceased kidney donors. Samples from the Biopsy Biobank Cohort of Indiana (BBCI)^97^ were acquired under waiver of consent as approved by the Indiana University Institutional Review Board (IRB # 1906572234). The TRIBE-AKI and ASSESS-AKI Studies (IRB00169832), the Hopkins Healthy Reference Study (IRB00199993) and the NAIKID study (IRB00221958) were approved by the Johns Hopkins University institutional review board. For the Boston Kidney Biopsy Cohort (Study, the Mass General Brigham institutional review board approved the study protocol (IRB #2012P000992). Data from Renal-HEIR (ClinicalTrials.govIdentifier: NCT03584217) and the IMPROVE-T2D study (ClinicalTrials.gov Identifier: NCT03620773) were included in this analysis. The Renal-HEIR and IMPROVE-T2D cohorts have intentionally harmonized study protocols and both were approved by the Colorado Multiple Institutional Review Board. Participants and/or parents provided written informed assent and/or consent, as appropriate for age. Participants who opted to undergo the optional kidney biopsy specifically and additionally provided consent to the research and biopsy teams.

Medication use was recorded for all participants, and T2D treatment was prescribed at the discretion of their medical provider. Normative reference tissue was provided by 6 healthy adult participants in the Control of Renal Oxygen Consumption, Mitochondrial Dysfunction, and Insulin Resistance (CROCODILE) study (ClinicalTrials.gov Identifier: NCT04074668) ^91^.

*Animal studies.* The animal study protocol was approved by the Institutional Animal Care and Use Committee of the Washington University School of Medicine, in adherence to standards set in the Guide for the Care and Use of Laboratory Animals (#22-0105, approval 15 July 2022).

### Clinicopathological Assessment of Human Tissues

<u>Pathology Descriptor Scoring.</u> The Kidney Precision Medicine Project (KPMP) tubulointerstitial and vascular (TIV) descriptor scoring parameters were developed based on the NEPTUNE Digital Pathology Scoring System ^98^. This form includes 64 unique TIV descriptors, of which 10 are scored as percentages observed across the cortex and medulla, while the remaining descriptors are assessed categorically within the cortex or medulla. Two KPMP pathologists—a primary scorer and a quality control (QC) scorer—review and score all stained sections (H&E, PAS, trichrome, and Jones silver histochemical stains, two of each) scanned into whole-slide images and stored in the KPMP Digital Pathology Repository, following the KPMP Manual of Operating Procedures (https://drive.google.com/file/d/1bcM_Z0GDLRTLZKXQmDYvAyCm-1tu9zBG/view). Scoring discrepancies between the primary and QC pathologists are discussed and resolved through adjudication, with a third pathologist consulted if necessary. These descriptors provide a comprehensive list of non-glomerular structural and cellular abnormalities observed in these regions.

<u>Clinicopathologic adjudication.</u> The KPMP Biopsy Adjudication Committee is comprised of nephrologists and nephropathologists from across the consortium. The adjudication process consists of a structured clinical case presentation followed by histopathology review of light microscopy, immunofluorescence, and electron micrographs. Nephropathologists systematically grade glomerular, tubulointerstitial, and vascular compartment features. Following clinical and pathology data review and discussion, committee members select a consensus primary diagnosis from the following categories: “diabetic nephropathy”, “vascular nephrosclerosis”, “cannot determine,” or “other”. Diabetic nephropathy is defined using Renal Pathology Society criteria ^99^. “Cannot determine” is used for primary diagnosis in cases without predominant classical features of diabetic nephropathy or vascular nephrosclerosis, or clear evidence of alternative kidney pathology.

### Single cell Processing of Tissue Specimens

<u>Tissue processing and dissolution.</u> Single cells (sc) and single nuclei (sn) were isolated from human specimens using protocols outlined previously^1^. For 10X snRNA-seq and Multiome assays, tissues were processed according to the following protocol: dx.doi.org/10.17504/protocols.io.568g9hw and nuclei were isolated from cryosectioned tissues according to the following protocol: dx.doi.org/10.17504/protocols.io.ufketkw. For 10X multiome assays, tissue sections were cut then stored on dry ice until nuclei isolation and Protector RNase Inhibitor (Sigma-Aldrich, Catalog #3335402001) was used at a concentration of 1.0 U/μl. To confirm tissue composition for nuclei preparations, 5 µm sections (flanking the thick sections used for isolations) were obtained for histology and the relative amount of cortex or medulla composition. For 10X scRNA-seq assays, tissues were preserved using CryoStor® (Stemcell Technologies). Single cells were isolated from frozen tissues according to the following protocol: dx.doi.org/10.17504/protocols.io.7dthi6n. The single cell suspension was immediately transferred to the University of Michigan Advanced Genomics Core facility for further processing.

<u>10X Chromium Assays and Sequencing.</u> 10X snRNA-seq and 10X Multiome was performed according to dx.doi.org/10.17504/protocols.io.86khzcw and dx.doi.org/10.17504/protocols.io.5qpvoby69l4o/v2, respectively. 10X scRNA-seq was performed according to dx.doi.org/10.17504/protocols.io.7dthi6n. These assays included both the 10X Chromium Single-Cell 3’ Reagent Kit v3 and the Chromium Next GEM Single Cell (ATAC + Gene Expression) Kit v1 or v2. RNA and ATAC libraries were sequenced separately on the NextSeq 2000 or NovaSeq 6000 (Illumina) systems (NextSeq 2000 Control Software v1.41.39716, NovaSeq Control Software v.1.6.0 and greater). Sample demultiplexing, barcode processing, and gene and ATAC peak count quantifications were performed with the 10X Cell Ranger v7.0.0 (RNA-seq) or the 10X Cell Ranger Arc v2.0.2 (Multiome) pipelines using the GRCh38 (hg38, GRCh38-2020-A-2.0.0) or a a pre-built GRCm39 (mouse) reference genome. For single nucleus data, introns were included in the expression estimates.

<u>Single cell Processing of Mouse Tissue Samples.</u> Ischemia perfusion injury (IRI) was performed on 8-10 week old male mice (18 min of renal pedicle clamp followed by perfusion) and kidneys were harvested at 2d, 28d and 5 months after the procedure, bivalved and preserved either in O.C.T. in cryocassettes at –80 °C or fixed in 4% PFA and paraffin embedded ^100^. Healthy kidney tissues from young to old mice were obtained from The Jackson Laboratory preserved in O.C.T. Nuclei were isolated from cryosections and subjected to 10X multiome pipeline using same procedures as for human samples. Control mice underwent sham surgery with no IRI at each of the time points. Serum BUN values were measured at each time point and for each mouse to monitor acute injury and recovery and adjacent slides of the sections from which nuclei were isolated were used for pathological assessment. The details are summarized in **Supplementary Table 15.** Nuclei were isolated and processed for 10X multiome as described above.

### Single Cell / Nucleus Data Processing and Quality Control

<u>10X snRNA-seq.</u> Cell barcodes passing 10X Cell Ranger filters were used for downstream analyses. Mitochondrial transcripts (MT-* for human or mt-* for mouse) were removed, doublets were identified using the DoubletDetection software (v4.2) ^101^ and removed. All samples were combined across experiments and processed using Seurat (v5.1.0) ^102^ to keep cell barcodes having greater than 400 and less than 7500 (5000 for mouse) genes detected. To further remove low quality datasets, a gene UMI ratio filter (gene.vs.molecule.cell.filter) was applied using Pagoda2 (v1.0.12, github.com/hms-dbmi/pagoda2).

<u>10X scRNA-seq.</u> The read count matrices were initially processed using SoupX (v1.5.0) to correct for ambient mRNA contamination. Single-cell analysis of kidney tissue typically results in high mitochondrial read content. As a quality control step, a cutoff of < 50% mitochondrial reads per cell was applied. In order to reduce doublets or multiplets from the analysis, we used a cutoff of > 500 and < 5000 genes per cell.

<u>10X Multiome.</u> RNA data was processed the same as 10X snRNA-seq. ATAC data was processed using Signac (v1.14.0) ^103^. Peaks called using Cell Ranger Arc were combined across experiments using the reduce function. Fragment objects for each experiment were prepared from Cell Ranger Arc fragment files using the CreateFragmentObject function. The combined set of peak regions was used to generate peak-by-cell matrices for each experiment using the FeatureMatrix function. Only cell barcodes that were annotated from RNA analyses were used for further analyses. Peak matrices were used to create individual Seurat objects that were merged to form four sets of combined objects based on the assigned (RNA) subclass level 1 annotations (set1: POD, PEC, PT, DTL, ATL; set 2: TAL, DCT; set 3: CNT, PC, IC, PapE; set 4: EC, VSM/P, FIB, Ad, Lymphoid, Myeloid, NEU). Accessible peaks were then called separately for multiple levels of cell type annotations within each set (clusters, subclass level 3 and subclass level 1) using the CallPeaks function and MACS (3.0.0b1; https://github.com/macs3-project/MACS). Called peaks across each set were combined using the reduce function and filtered to remove: (1) regions >10000 and <20 base pairs; (2) regions falling within nonstandard chromosomes; (3) regions occurring in blacklist regions using the blacklist_hg38_unified object from Signac. For mouse blacklist regions we instead used mm39 exclusion regions (AH107321) using the excluderanges R package (v0.99.8) ^104^. The final peak sets were used to create new peak-by-cell count matrices and Seurat objects for each subclass level 1 grouping as detailed above. Gene annotation of the peaks was performed using GetGRangesFromEnsDb(ensdb = EnsDb.Hsapiens.v86) for human and rtracklayer (v1.64.0) import of the cellranger GRCm39 reference genes.gtf for mouse. Nucleosome signal (NS) scores, transcription start site (TSS) enrichment scores and fraction of reads in peaks (FRiP) were calculated for each cell using the Nucleosome Signal, TSS Enrichment and FRiP functions. Cell barcodes further passing the following ATAC filters were kept for downstream analyses: (1) >1000 and < 70,000 peak counts per cell; (2) a FRiP greater than 0.25; (3) a NS score less than 4; (4) a TSS enrichment score >2. Combined Seurat objects were created with separate RNA and ATAC assays for the same cell barcodes.

### Single Cell / Nucleus Data Clustering, Cell Type Annotation and Analyses

<u>Clustering single nucleus RNA data.</u> Clustering and cell type annotations were performed using a previously established pipeline^1^. Given the ongoing sample collection and data production associated with the KPMP and HuBMAP projects, the clustering analysis was first performed on the first 244 samples (**Supplementary Table 1**), then the remaining 64 samples were integrated and aligned to these clusters. This provided a mechanism for new data incorporation while also assessing cluster stability across aggregated data set batches.

For the first 244 samples, cells were first annotated to the Human Kidney Atlas V1^1^ to ensure consistency of cell types between atlas versions. This involved a strategy previously used to merge data across technologies. For this, cell types from Atlas V1 were predicted in Atlas V2 data using the Seurat MapQuery function, then integrated clustering was performed on the imputed principal components using Pagoda2. Individual clusters were annotated based on V1 atlas predicted cell type labels, cluster identities, subclass correlation values, and marker gene expression profiles. Hard to resolve subpopulations were manually separated using the same process as in the V1 atlas^1^ either using gene expression (e.g. REN for REN+ cells) or through sub-clustering (EC-AEA, EC-DVR). Clusters showing overlapping or indiscriminate cell type markers were tagged as ambiguous low-quality clusters and excluded.

V2 clustering was then performed on both V1 and V2 data (first 244 samples, see Supplementary Table 1). First, for additional data clean up, a sketch object (Seurat V5) was generated consisting of 200,000 cells which were clustered using Pagoda2. These clusters were then projected to the full data set (Seurat V5) using the ProjectData function and V1 annotations assessed. Clusters showing overlapping or mixed V1 cell type labels were assessed for marker gene profiles and differentially expressed genes (DEGs). Ambiguous clusters (very low DEGs and indistinct cell type identities) or multiplet clusters (mixed cell type marker profiles) were tagged as ambiguous low quality and excluded. Clustering was then performed separately for proximal epithelial cells (POD, PEC, PT, DTL, ATL), distal epithelial cells (TAL, DCT, CNT, PC, PapE, IC), stromal cells (FIB, VSM/P, Ad), endothelial cells (EC), immune cells (Lymphoid, Myeloid) and Schwann/neural cells (NEU) in order to achieve a higher resolution of cell type discovery. For each cell subset, clustering was performed using pagoda2, where counts were normalized to the total number per nucleus, batch variations were corrected by scaling expression of each gene to the dataset-wide average. After variance normalization, all significantly variant genes were used for principal component analysis. An initial round of clustering was performed at high resolution (e.g. k = 100) in order to identify any multiplet clusters showing mixed cell type marker profiles, which were subsequently excluded. Then a second round of clustering was performed at different k values (e.g. 100, 200, 500) based on the top 50 principal components, with cluster identities determined by the infomap community detection algorithm. The primary cluster resolution (e.g. k = 200) was chosen based on the extent of clustering observed. Principal components and cluster annotations were then imported into Seurat and uniform manifold approximation and projection (UMAP) dimensional reduction was performed using the top 50 principal components identified using pagoda2. Subsequent analyses were then performed in Seurat. A cluster decision tree was implemented to determine whether a cluster should be merged, split further or labeled as an altered state. For this, differentially expressed genes between clusters were identified for each resolution using the FindAllMarkers function in Seurat (only.pos = TRUE, max.cells.per.ident = 2000, logfc.threshold = 0.25, min.pct = 0.25). Possible altered states were initially defined for clusters having one or more of the following features: low genes detected, high number of mitochondrial transcripts, high number of ER associated transcripts, upregulation of altered state markers (from Atlas V1) or enrichment in AKI or CKD samples. Clusters (k = 200) with fewer than two distinct markers were assessed for altered state features, if present then these clusters were tagged as possible altered states, if absent then these clusters were merged if possible to the largest cluster of that subclass as guided by their cluster resolution at k = 500. Any clusters that were annotated as ambiguous low quality (mixed cell type marker profiles, originating from only one or two individuals or showing indistinguishable marker gene expression) were tagged and removed. This gave a final set of 159 high quality V2 clusters showing either distinct marker gene expression or altered state features.

Data for the next 64 samples was then incorporated following a similar strategy as outlined above, where the V2 clusters were initially predicted using the Seurat MapQuery method, then co-clustering was performed using Pagoda2. Clusters (k = 200 or if necessary k = 100) were then annotated for the query nuclei based on the majority V2 cluster identities of co-occurring reference nuclei. This strategy accurately recapitulated the majority of the 159 clusters, which were confirmed by assessing cell type marker gene expression profiles. Clusters that did not resolve with the new data even at lower k values were found to occur only for degenerative states and likely represent unstable altered states not found more broadly across patient samples (see **Supplementary Table 10)**.

For mouse single nucleus data, a similar clustering strategy was applied. First, human Atlas V2 subclass labels were predicted in 10x multiome RNA data (**Supplementary Table 12**) using only mouse genes having one-to-one human orthologues (obtained from Ensembl) and the FindTransferAnchors and MapQuery functions in Seurat. Then the full mouse 10x multiome RNA count matrix was clustered using Pagoda2 and clusters annotated to a broad subclass level based on predicted annotations and marker gene expression. Ambiguous mixed identity clusters (multiplets) were removed and the data was re-clustered and re-annotated to subclass level 1. The 10X Multiome RNA data was then combined with 10X snRNA-seq data (assay version 2) ^14^ and integrated using a sketch subset (50,000 nuclei) using the reciprocal PCA (rPCA) integration strategy (Seurat). Clustering on these integrated rPCA embeddings was performed using Pagoda2 and ambiguous mixed identity clusters (multiplets) were identified based on marker gene expression. Clusters were projected to the full data set using the ProjectIntegration and ProjectData functions in Seurat and multiplets were removed before repeating the sketch, rPCA and Pagoda2 clustering process. An initial cluster annotation was established using human predicted identities, published annotations, and marker gene expression profiles derived from the human Atlas V2 (**Supplementary Table 12**) and published mouse studies ^14,16,18^. Subgroups of broad cell types, similar to human atlas subgroups, were then clustered independently using the rPCA and Pagoda2 clustering strategy. For each subset, an initial round of clustering was performed to identify and exclude ambiguous low quality clusters (multiplets), allowing for multiple rounds of data cleanup.

<u>Annotating single nucleus clusters.</u> Cell type and state annotations were assigned based on evidence from multiple strategies, leveraging a cross-consortium knowledge base, including domain expertise from clinicians, pathologists, biologists, and ontologists. As such, the assignment of identities for each cluster took in account: gene expression profiles of known cell type markers ^1,10,105,106^ (also see **Supplementary Table 10**); enrichment with clinical categories or pathology descriptor scorings; conservation with cell states found in mouse injury models, including relative abundance along a known timeline from acute injury ^14,18^; correlation of gene expression profiles with published cross-organ disease-associated cell states ^8,11,15^; expression of known signaling pathways or altered state (e.g. cell cycle) signatures; regional distribution across the corticomedullary axis or their spatial localization to anatomical structures or niches. To further align immune clusters with known immune cell types, we took in account assignments by the CellTypist python package (version 1.4.0) and associated v2 immune cell type labels^10^. Accumulated evidence from these strategies enabled the adoption of a standardized anatomical and cell type nomenclature for major and minor cell types and their subclasses (**Supplementary Table 10**), also see the HuBMAP ASCT+B Reporter: hubmapconsortium.github.io/ccf-asct-reporter. This led to a higher level of cell type resolution with more detailed altered state annotations associated with this Atlas Version (V2) release.

<u>Clustering single cell RNA data.</u> Using Seurat 5.0 ^102^ functionalities, each sample group was independently normalized, had variable genes identified, and underwent principal component analysis (PCA). Subsequently, all samples were integrated using reciprocal PCA (RPCA). The integrated dataset then underwent dimensionality reduction via Uniform Manifold Approximation and Projection (UMAP) and unsupervised clustering at a resolution of 1.5. Cell clusters were annotated based on markers of major cell types, including major renal epithelial, endothelial, interstitial, and immune cells. To identify stable cell states, cells from each major cell type were sub-clustered following the previously mentioned steps. Unsupervised clustering of each cell type was performed using Leiden algorithm at multiple resolutions (0.2-1.2). Cluster stability was calculated using the ‘calc_sc3_stability_cluster’ function in the Clustree R package ^107^, and for each cell type, we chose the resolution with the highest median stability index. The sub-clusters at the chosen resolution were then checked to ensure they contained cells from multiple sample categories. The sub-clusters are then annotated based on markers from KPMP version 1.0 and literature ^10,11,106,108^.

<u>Integrating single cell and single nucleus data sets.</u> The integration of immune cells from single nucleus and single cell data sets was performed using the ‘CCAIntegration’ functionality in Seurat (v5). Lymphoid and myeloid cell integrations were conducted separately. Cell types were annotated based on integrated cluster markers and relevant literature ^10,11^. Two immune experts, Drs. Nir Hacohen and Deepak Rao, reviewed and approved these annotations. The transcript profile of renal epithelial cell states differed between single-cell and single-nucleus analyses due to factors such as post-translational modifications (PTMs), RNA stability, and localization within the cell. Therefore, we integrated snCv3 and scCv3 renal epithelial cell types based on enriched pathway mechanisms identified from their specific markers.

<u>Cell type or cluster enrichment scoring (single nucleus).</u> To determine cluster or subclass enrichment between different conditions or pathology descriptor scoring categories (see above), the proportion of each cluster or subclass was calculated for each patient/sample. Two-sided t-tests were then used to assess significance for differences in the relative abundance of each cluster or subclass between two different patient/sample groupings. The t statistics were plotted using the pheatmap R package (version 1.0.12) and differences having a p value less than 0.05 were indicated using an Asterix. Clinical patient groupings included: healthy reference tissues (HRT), acute kidney injury (AKI), chronic kidney disease (CKD), estimated glomerular filtration rate or eGFR either high (≥ 60 ml/min/1.73m^2^) or low (< 60 ml/min/1.73m^2^), and age groupings for HRT < 50 years old or ≥ 50 years old. Clinical categories from KPMP adjudications included: AKI patient subset showing ATI; AKI patient subset showing acute interstitial nephritis (AIN), CKD with low end stage risk (CKD^lo^), CKD with high end stage risk (CKD^hi^); Diabetic Kidney Disease (DKD), Hypertensive Chronic Kidney Disease (H-CKD). Pathology descriptor scoring categories included: Interstitial Fibrosis or IF (≤20% vs >20%); Tubular Atrophy (common type) or TA (<20% vs ≥20%); Acellular Casts or AC (<20% vs ≥20%); Tubular Injury (other than atrophy) or TI (<20% vs ≥20%); Interstitial Mononuclear White Blood Cells or WBC (≤10% vs >10%); Arteriosclerosis or AS (0-1 vs 2-3); Arteriolar Hyalinosis or AH (0-1 vs 2-3). For pathology descriptor scores, only patients having >25 % cortex for the pathology scoring were included, and single nucleus patient samples that were predominantly medulla were excluded. Actual p value, t statistics and associated patient numbers per patient grouping are included in the Supplementary Data File.

<u>Correlations with published cell types or between species (single nucleus).</u> For comparison of kidney fibroblast states with those found in disease tissues, we obtained processed data from ^8^ (https://fibroXplorer.com) and ^15^. For comparison of myeloid states with those found across human tissues or found in a mouse model of reversible unilateral ureteral obstruction (UUO), we obtained processed data from Eraslan et al ^11^ (http://www.gtexportal.org/) and directly from the authors of Conway et al ^9^. For correlation of mouse to human data, only genes having one-to-one human orthologues (obtained from Ensembl) were used. In each case, average scaled expression values for a set of variable genes were calculated using Seurat (FindVariableFeatures and AverageExpression functions) and used for correlation of cell types between data sets. Correlation values were then visualized using the corrplot R package (version 0.95).

<u>Trajectory analyses (single nucleus).</u> To identify potential paths for cell state shifts, cell type subsets of healthy and adaptive state clusters were used to re-calculate PCA and UMAP embeddings using Pagoda2 and Seurat, respectively. Potential lineages from a specified starting cluster were then estimated using the R package Slingshot (version 2.12.0). To generate embeddings for individual lineages, relevant clusters from each estimated slingshot lineage were subset and PCA and UMAP embeddings were re-calculated as above.

<u>Gene set expression scores (single nucleus).</u> Per cell gene set expression scores were calculated for single nucleus RNA data using the UCell R package (version 2.8.0). Gene sets, associated genes and their sources are provided in **Supplementary Table 7**. Gene sets for signaling pathways (Androgen, EGFR, Estrogen, Hypoxia, JAK-STAT, MAPK, NFkB, p53, PI3K, TGFb, TNFa, WNT, Trail, VEGF) were obtained from the PROGENy R package (version 1.26.0) after taking the top 100 genes per pathway ranked by p value^109^. IFNB and INFG pathways were obtained from Perturb-Seq derived gene sets ^110^. MSigDB collections (gsea-msigdb.org) were obtained using the msigdbr package (version 7.5.1). CKD progression gene sets were identified from plasma protein associations in patients from the Chronic Renal Insufficiency Cohort Study (CRIC) and in the Atherosclerosis Risk in the Communities study (ARIC) ^96^. Acute tubular injury (ATI) gene sets were obtained from plasma protein associations in patients from the Boston Kidney Biopsy Cohort (BKBC), ARIC and the COVID-19 Host Response and Clinical Outcomes (CHROME) study ^53^. Proteomic age associated gene sets (ProtAge) were identified as plasma proteins predictive of chronological age ^95^. Gene sets scores were visualized by first being added to a separate assay slot in Seurat, scaled across all cell types using the ScaleData function, averaged by cell types, and then plotted using the pheatmap package (v1.0.12).

<u>Cell state marker genes for clinical cohort associations (single nucleus).</u> To identify genes that may be associated with disease progression, cell states associated with PT, TAL, C-FIB, pvFIB and myeloid cells (**Figures 2-3**) were used. Within each level 1 subclass, differentially expressed genes were identified between level 3 subclasses using the FindAllMarkers function in Seurat (only.pos = TRUE, logfc.threshold = 0.25, min.pct = 0.25). These were then intersected with a list of genes associated with secreted proteins obtained from the Human Protein Atlas (proteinatlas.org). Genes associated with proteins identified as significantly associated with AKI and AKI progression were visualized across cell types within the single nucleus RNA-seq by averaging scaled expression values for each level 3 subclass and plotting using the pheatmap package.

<u>Cell type pathway analyses.</u> Upregulated genes within each cell subtype, if compared to all other subtypes of the same cell type or group of cell types were calculated using the Seurat functionality “FindAllMarkers” ^102^. We standardized the maximum number of cells considered by “FindAllMarkers” for each subtype (i.e., Seurat ‘identity’) to 5000 to reduce potential differences in cell abundancies between the different subtypes. To ensure reproducibility and avoid selection bias, we averaged the results of 25 different sets of upregulated genes obtained by making the algorithm sample 25 different sets of 5000 cells for each subtype (i.e., by specifying 25 different random seed numbers). We assessed this procedure by averaging between 1 and 100 distinct sets of 5,000 cells (randomly drawn from a pool of 10000 different sets). Each case was repeated 100 times to evaluate consistency. To identify significant genes, we first removed all genes with an (averaged) adjusted p-value of more than 0.05 and then kept only the top 500 most significant genes, if this number was exceeded. Subtype-selective significant genes were subjected to pathway enrichment analysis using Fisher’s exact test and level-3 pathways of the Molecular Biology of the Cell Ontology (MBCO) (github.com/SBCNY/Molecular-Biology-of-the-Cell, mbc-ontology.org) ^19^. Predicted significant subtype-selective pathways (nominal p-value ≤ 0.05%) and expressed pathway genes were expert-curated to exclude pathways that were predicted based only on genes that are not selective for the function of the pathway. Pathways that were predicted based on selective genes were grouped into 14 different categories of overall whole cell functions. Minus log10 p-values of all pathways mapping to the same overall function were summed up to calculate an enrichment score for the overall whole cell function of interest. These enrichment scores were either visualized as pie slices in pie charts or color coded within heatmaps.

Disease-selective pathways and overall cellular functions were calculated similarly, except that we only considered samples obtained by KPMP biopsies. For each cell type or cell type group, we calculated marker genes for the recruitment diagnoses AKI, CKD with an eGFR below 45 and CKD with an eGFR above or equal 46. Each time cells annotated to the same cell type/cell type group and the other two recruitment diagnoses were used as the denominator.

<u>Druggable</u> <u>Targets.</u> We used Drug2cell, a computational framework developed by the Wellcome Sanger Institute [https://www.sanger.ac.uk/technology/drug2cell/] to predict drug targets from snRNA-seq data by mapping drug activity to cell types based on gene expression profiles using the ChEMBL database [https://www.ebi.ac.uk/chembl/]. Our application of Drug2cell to screen 2.5 million compounds against 89 cell clusters in the HKAv2 kidney atlas identified several cell-type-specific compounds with potential therapeutic relevance with several drugs targeting known kidney structures confirming the approach. Of these few were highlighted using manual examination of dot plots based on their selective activity in distinct kidney cell populations.

<u>Cross mapping with diabetic and hypertension mouse models.</u> The mice dataset consists of snRNA-seq data from a total of 69 mice, divided into 14 groups^94^. These groups include controls (db/m), diabetic mice (db/db), diabetic hypertensive mice (db/db + AAV), and treated diabetic hypertensive mice (db/db + AAV + treatment). Treatments administered include ACE inhibitors, SGLT2 inhibitors, and Rosiglitazone, with treatment durations of 2 days or 2 weeks. The data captures a wide array of kidney cell types, including proximal tubule cells (PT), thick ascending limb cells (TAL), and interstitial fibroblasts, among others. The normalized data were obtained from the publicly available GEO database under accession number GSE184652. To integrate the snRNA-seq data from human and mice kidneys, we employed Seurat v5.0.3, leveraging its reciprocal principal component analysis (rPCA) algorithm for batch correction. The integration process involved over 2.3 million nuclei from both species, necessitating computational strategies such as sketching to reduce data size while retaining essential biological variability. This allowed for the alignment of conserved cell types, such as proximal tubule cells (PT), thick ascending limb cells (TAL), and interstitial fibroblasts, across species. Orthologous gene mapping was conducted using the NCBI Datasets v2 toolkit. All analyses were performed using R version 4.1.0. To evaluate the activity of transcription factors (TFs) across different conditions, enrichment scores for their downstream target genes (see GRN analyses section below) were calculated using the single-sample Gene Set Enrichment Analysis (ssGSEA) method. The downstream gene sets for each TF (**Supplementary Table 21**) were aggregated at the sample level for each cell type to generate pseudo-bulk RNA-seq data, ensuring that the analysis accounted for variability inherent in single-cell data while maintaining robust biological signals. The ssGSEA method was implemented using the GSVA R package (version 1.42.0) to compute enrichment scores, which represent the relative activation or repression of each TF’s downstream gene set for individual samples. The TF activity scores were computed for all mice and compared between healthy and diabetic hypertensive mice, as well as between treated and untreated diabetic hypertensive mice. Statistical significance was assessed using the Wilcoxon rank-sum test, with a threshold of p < 0.05. Positive ssGSEA scores indicate increased activity of the TF, while negative scores suggest decreased activity.

### Gene Regulatory Network (GRN) Analyses

<u>Transcription factor activities</u>. Transcription factor binding site (TFBS) activities were generated and visualized using Signac. Jaspar motifs (all vertebrate) from either JASPAR2020 (scMEGA analyses) or JASPAR2022 were used to generate motif matrices and motif objects using the AddMotifs function. Motif activity scores were then calculated using ChromVAR (v1.26.0; https://greenleaflab.github.io/chromVAR) ^111^ using the RunChromVAR function.

<u>Single-cell Multiomic Enhancer-based Gene Regulatory Network Inference (scMEGA)</u>. To link accessible chromatin peaks to transcription factor binding site (TFBS) activities and build trajectory-associated gene regulatory networks, we used scMEGA^23^. For each cell state lineage identified from the trajectory analyses described above, we first built a pseudotime trajectory using the R package ArchR (v1.0.2) ^112^. Then scMEGA calculated the correlation of TF binding activity estimated using chromVar and the corresponding TF’s expression level. The top 10% of genes showing variable expression across the trajectory were then correlated by expression level with their peak accessibility using ArchR, with enhancers selected as peaks that were within 2k base pairs from their linked gene. scMEGA then built a quantitative GRN through correlation of TF binding activities to expression of selected genes, then subset this to an enhancer-based GRN through consideration of only chromVAR TFBS activities found within gene-associated enhancers. TF-gene interactions within these networks were weighted by their correlations. GRNs were modeled as a graph and assessed for directed (TF-to-gene) connectivity using the R package igraph (v2.1.1). To select TF-driven target gene sets from these networks (**Supplementary Table 21**), only TFs having betweenness centrality in the networks and showing correlation of their binding site activities and their associated gene expression levels that was greater than 0.6 (or 0.55 for macrophage trajectories) (**Supplementary Table 22**). Target genes for these TFs where then selected based on being in the top 200 correlated genes as ranked by p values and having a correlation value greater than 0.6. Averaged GRN scores by cell type were visualized as detailed for gene set expression scores above using the pheatmap package.

<u>MAGICAL</u>. To identify regulatory circuits involving transcription factors, chromatin sites, and genes with coordinated activity variation across cell clusters in each cell lineage, we applied MAGICAL ^113^ to single-cell multiome data. For each cluster, candidate peaks and genes were identified as significantly upregulated in the selected cells in this cluster relative to other lineage cells using Seurat V5. Transcription factor (TF) binding sites were mapped to candidate peaks using TF motifs from the chromVARmotifs library using Signac. Candidate regulatory circuits were constructed by linking TF binding sites to genes with transcription start sites (TSS) within 500 kb. MAGICAL (v1.0) was then applied to identify regulatory circuits in the selected cluster, using all lineage cells as a reference. From these, the top 10% high-confidence circuits were selected, and results across clusters were pooled to obtain the final set of circuits for the lineage. Circuit peaks were further validated by overlapping with CUT&RUN-seq peaks, retaining those with center-to-center distances under 1 kb for enhanced confidence.

<u>GWAS analyses</u>. To identify regulatory circuits potentially associated with disease, MAGICAL circuits overlapping with GWAS signals^59^ were selected and annotated at three levels. First, circuits were flagged if their peaks overlapped genome-wide significant SNPs. Second, circuits were linked if their genes corresponded to GWAS-annotated genes. Third, circuits were included if either their peaks or genes fell within a GWAS loci containing at least one genome-wide significant SNP. All GWAS SNPs and loci were converted from hg19 to hg38 coordinates using CrossMap (v0.7.3) for compatibility with the analysis.

<u>Cleavage under targets & release using nuclease (Cut&Run)</u>. Reference kidney sample processing, library preparation, sequencing and analysis has been published previously^54^. Kidney biopsy samples were processed according to a modified protocol adapted from Epicypher and available on protocols.io (dx.doi.org/10.17504/protocols.io.bp2l615o1vqe/v1). Antibodies used for CUT&RUN reactions: H3K27ac (Cell Signaling, 8173), H3K27me3 (Cell Signaling, 9733), H3K4me1 (Cell Signaling, 5326), and IgG (Cell Signaling, 2729) at a 1:50 dilution. Up to 8 ng of DNA from biopsy samples was used to prepare sequencing libraries using the NEBNext Ultra II Library Kit (NEB, E7645) following manufacturer’s instructions, using 17-18 cycles to amplify the library. For sequencing, 0.8 pm of the library was sequenced on the NovaSeq 6000 (Illumina) targeting 100 million paired-end reads. Detailed bioinformatic analysis with command line examples can be found on protocols.io ^54^. Briefly, fastq files were trimmed using Cutadapt (v3.7) and aligned to the hg38 reference genome using Bowtie2 (v2.4.5). Aligned reads were extracted and converted to a sorted bam file using SAMtools (v1.19.2). RPKM normalized BigWig files were generated using Deeptools (v3.5.1), and peaks were called using Macs2 (v2.2.6). The number of reference sample libraries processed for each antibody are: H3K27ac N=10 replicates from 5 distinct biologic, H3K27me3 N=14 replicates from 6 biologic, H3K4me1 N = 7 replicates from 2 biologic, H3K4me3 N=7 replicates from 2 biologic. The number of diseased kidney biopsy sample libraries processed for each antibody are: H3K27ac N=12 replicates from 10 distinct biologic, H3K27me3 N=12 replicates from 10 biologic, H3K4me1 N = 12 replicates from 10 biologic, H3K4me3 N=12 replicates from 10 biologic. To visualize Cut&Run coverage tracks, reference or diseased biopsy tissue bam files were merged using SAMtools (v1.9) and a merged RPKM normalized bigwig file was generated using Deeptools (v3.5.2) and visualized with the Integrative Genomics Viewer (v2.19.4) along with pseudobulk ATAC for proximal tubule cell cluster trajectories. To visualize proximal tubule ATAC coverage, single cell ATAC fragment files were split by cell cluster using the SplitFragments function, and bed files were converted to RPKM normalized bigwig files using Deeptools (v3.5.2). For validation of MAGICAL circuits, merged Cut&Run bam files for reference or diseased biopsy samples were used to call peaks using Macs2 (v2.2.6). The resulting peak set was overlapped with the high-confidence MAGICAL circuit peaks.

*<u>In silico</u>* <u>Transcription Factor Disruptions</u>. Following modeling of cell trajectories and the identification of regulatory interactions with scMEGA (v.1.0.2) ^23^, an *in silico* knockout was performed with CellOracle (v.0.12.0) using the GRN generated by scMEGA. CellOracle ^114^ simulation tools predicted the downstream effects on target genes and cell states, allowing us to infer the regulatory impact of TF loss. Individual and combined knockouts of *SOX9* and *SOX4* were simulated *in silico*, and changes in gene expression and cell velocity were calculated for the PT and TAL failed repair trajectories.

<u>DecoupleR differential expression analysis and TF enrichments</u>. To identify differentially expressed genes (DEGs) between AKI patient sub-groups (recovering versus progressing) within early to mid-repairing PT states, we used the decoupleR python package (version 1.9.1) ^115^. First, pseudo-bulk profiles were generated using raw counts for PT subclasses (level 3) for each patient. Genes were filtered using the filterByExpr function (min_count=10, min_total_count=15) and DeSeq2 was then performed for aPT cells (subclass level 2) grouped by the AKI conditions using the PyDESeq2 package (version 0.5.0). DEGs were visualized using the volcano plot (plot_volcano_df). P-values were calculated using the Wald test and adjusted for multiple testing. Transcription factor inference on these DEGs was performed using TF-gene network connections identified using scMEGA analysis of the aPT2 to PT-S1 repair trajectory. TF enrichment scores were generated using the Univariate Linear Model (ulm) method. The top 25 inactive or active TFs between AKI groups was visualized using the plot_barplot function. SOX4 target genes were visualized using the plot_volcano function.

<u>Ligand-Receptor (LR) interaction analyses</u>. To identify which cell states in the spatial niches containing tubular epithelial cells interact through CCL2-CCR2 or IL1B-IL1R1, we subset the single-nucleus atlas to the relevant epithelial cells, fibroblasts and immune cells. Ligand-receptor inference was performed on normalized log1p-transformed gene expression data using the rank_aggregate method from the liana-py package (version 1.4.0) ^116^, which provides the consensus output from 5 individual ligand-receptor inference methods, and setting the expr_prop setting to 0.05. Figure 4 shows interactions between cell states where at least one interaction has a specificity rank < 0.1. For clarity, we also add the non-significant interactions involving epithelial cells (i.e. PT states).

To find ligand-receptor interactions between epithelial cells and fibroblasts or immune cells that are differential across healthy and adaptive states, we again infer interactions with liana-py rank_aggrgate function using normalized log1p-transformed gene expression and setting the expr_prop setting to 0.1. This time we use the consensus ligand-receptor database and filter for ligands and/or receptors that are a marker gene of at least one epithelial state (i.e. either PT or TAL). Furthermore, we are narrowing down interactions to only those involving epithelial cells as source or target of the interactions. LR-interactions are narrowed down to those that have at least one significant interaction with specificity rank < 0.01, and are then ranked by how variable they are across epithelial states.

### SLIDE-Seq2 Spatial Transcriptomics

<u>Puck preparation and sequencing</u>. Tissue pucks were prepared from fresh frozen kidney tissue either embedded in OCT or frozen in liquid nitrogen and sequenced according to the step-by-step protocol: dx.doi.org/10.17504/protocols.io.bvv6n69e. Libraries were sequenced using NovaSeq 6000 with a standard loading concentration of 2nM (read structure: Read 1 - 42 bp, Index 1 - 8 bp, Read 2 - 60 bp, Index 2 - 0 bp). Demultiplexing, genome alignment and spatial matching was performed using Slide-seq tools github.com/MacoskoLab/slideseq-tools/releases/tag/0.1.

<u>Cell type deconvolutions</u>. Slide-seq2 data analysis was performed using Seurat v5.0 and deconvolution of cell types was performed using RCTD from the spacexr package (version 2.2.1, github.com/dmcable/spacexr). Broad cell types (subclass level 1) were first predicted from a downsampled 10X single nucleus RNA reference atlas (10,000 nuclei per subclass with ATL and DTL merged, excluding subclasses that tended to mis-predict: PapE, Ad, Neu) using RCTD on each tissue puck separately. The prediction weights were normalized to sum to 100 per bead and only spots showing a max weight of > 30 were used. Count matrices were then combined across all pucks, a sketched data set of 100,000 spots was generated using Seurat and then clustered using Pagoda2. The resultant clusters and embeddings were transferred to the Seurat object and projected to the full data set using the ProjectData function. Clusters were then annotated to broad subclasses (level 1) based on RCTD predictions, correlation with the reference atlas subclusters and expression of marker genes. Clusters that were poorly predicted or showed too low marker gene expression were labeled as ambiguous and removed. This enabled more reliable level 1 subclass annotations which were then used as the basis for subsequent more resolved cell type predictions (v2.subclass.sp, **Supplementary Table 13**).

For this, broad groupings of proximal to intermediate tubules, distal tubules to collecting ducts and non-epithelial cells were independently used for the corresponding level 2 RCTD predictions. Reference objects were subset to remove degenerative and cycling cell states, then downsampled to 5000 nuclei per subclass (v2.subclass.sp). Prediction weights were again scaled to 100 and stored as a separate assay within the Seurat object. Spots falling outside the tissue and having level 2 prediction weights less than 25 were removed. Subclasses were assessed for marker gene expression and spatial localization and those showing less accurate predictions were subset based on non-zero expression of distinguishing marker genes (PECs – *CFH*; MD – *BBOX1*; resMAC-HLAII^hi^ – CD163, STAB1, C1QA). C-FIB-OSMR^lo^ were over-represented and likely represented a mixed population with C-FIB-PATH. aEC-GC were also over-represented and re-labeled as EC-GC. Accuracy of final predictions was assessed through: correlation of averaged scaled expression values with reference single nucleus RNA subclasses visualized using the corrplot r package; marker gene expression visualized using the DotPlot function in Seurat; correct spatial enrichment (normalized cell type fraction per puck) in cortex, outer medulla or inner medulla regions (assigned to tissue pucks based on adjacent histological sections), visualized using the ggplot2 r package (v3.5.1). Spatial mapping of cell types was performed in Seurat using the SpatialDimPlot function. To visualize TF-associated GRNs, gene set expression scores were calculated using the UCell R package (version 2.8.0) as described above for single nucleus data. Gene set scores were then mapped only for the associated broad cell type (PT, TAL, FIB or MAC) using the SpatialFeaturePlot function in Seurat.

<u>Neighborhood enrichment analysis.</u> For each cell in each puck (cortex only), we identified the 100 nearest neighbors and computed the number of cells belonging to each annotated subclass. This neighborhood count matrix was then normalized so that each row summed to 1, giving a neighborhood composition matrix (NCM). The NCM was filtered to only contain cell types associated with the sampled region (cortex). To assess the neighborhood of each cell type independently, the NCM was filtered to only contain that cell type. Next, the NCM was split into two arrays: one containing all rows corresponding to cells from healthy controls (healthy) and one containing all rows corresponding to cells from individuals with CKD/AKI (disease). To analyze whether cells of type B were enriched in the neighborhood of cells of type A, where the k-th column of the NCM corresponds to cells of type B, then we performed a two-sample Kolmogorov-Smirnov test using the k-th column of NCM-healthy and the k-th column of NCM-disease. Positive test statistics indicated that cells of type B were depleted (in the neighborhood of cells of type A) in healthy controls or alternatively were enriched in disease. The results were visualized using seaborn clustermap, where the clustering was performed only on columns. For visualization, test statistics were only shown for p-values less than 1e-2 and colors were clipped to the [-1, 1] range.

### Visium (10X Genomics) Spatial Transcriptomics

<u>Frozen Visium preparation, imaging, and sequencing</u>: We prepared and imaged human kidney tissue according to the Visium Spatial Gene Expression (10x Genomics) manufacturer protocol (CG000240 protocol) and as previously described ^117^. We sectioned frozen samples at a thickness of 10 µm from Optimal Cutting Temperature (OCT) compound embedded blocks.

Tissue was stained with hematoxylin and eosin (H&E). We acquired histology images using a Keyence BZ-X810 microscope equipped with a Nikon 10X or 20X CFI Plan Fluor objective. We isolated mRNA from tissue sections after 12 minutes of permeabilization. mRNA bound to oligonucleotides in the fiducial capture areas was reverse transcribed and underwent second strand synthesis, denaturation, cDNA amplification, and SPRIselect cDNA cleanup (Visium CG000239 protocol) for library preparation. We then sequenced the cDNA on an Illumina NovaSeq 6000.

<u>Formalin-fixed paraffin-embedded Visium preparation, imaging, and sequencing</u>: We prepared and imaged formalin-fixed and paraffin-embedded kidney tissue in an analogous manner to the frozen tissue, except with protocol CG000408. Samples were sectioned at 7 µm thickness from the paraffin blocks and placed on standard glass slides for use with the 10x CytAssist instrument. All downstream processing steps aligned with those of the frozen protocol.

<u>Gene expression analysis</u>: We performed expression analysis, mapping, counting, and clustering with Space Ranger (v2.0 or higher) in reference genome GRCh38-2020-A. Datasets were normalized with SCTransform ^117^ and processed in Seurat (v5.1) ^102,118^. In each Visium sample, the outermost layer of spots was eliminated to reduce edge artifact. Transcription factor activity for each spot was estimated based on the expression of genes in the predicted gene regulatory network with the Seurat module score function. TF activity was compared between AKI recovery with non-recovery and CKD progression with non-progression by Student’s t-test. AKI recovery was defined by a return to an eGFR above 30 ml/min/1.73m^2^ and improvement by 10 ml/min/1.73m^2^ or more. CKD progression was defined as a decline in eGFR of 15% or more between 2 and 3 years from time of biopsy.

Cell type localization: Using Seurat (v5.1) anchors method, we transferred labels from the multiome / snRNA-seq object clusters, using transfer scores as an estimate of the proportion of signature from each cell type that contributes to each 55 µm spot.

<u>Niche analysis (Visium)</u>: To define cellular niches we first attributed all ST spots (N = 126701) from 153 samples into 10 major cellular groups. Spots with the majority of cell type proportion from epithelial, endothelial, immune, or stromal cells were assigned to each of those groups. The epithelial group was further divided based on higher level 1 cell type proportion, if more than 20%. Spots classified as PT, DTL, TAL, DCT / CNT were assigned to respectively named groups. Those classified as POD or PEC were assigned to the Glomerular group, while PC or IC classified spots were assigned to CD group. Spots that do not classify as any of those groups were labelled as “other”. Within each group, niches were created by clustering spots according to their cell type proportions with traditional Louvain algorithm as implemented by Seurat v5. Niches were associated with clinical categories through a Fisher’s exact test and displayed as a forest plot.

<u>Histological validation</u>: Visium FFPE samples were used due to better histology preservations. To localize the niches defined in the larger OCT preserved dataset, we mapped the niche annotation using the anchor’s method from Seurat. To determine histologic structures in Visium FFPE samples, we annotated every non-edge spot according to the following functional tissue unit categories: 1) non-sclerotic glomerulus, 2) globally sclerotic glomerulus, 3) large artery vessel, 4) cortical tubulointerstitium without fibrosis or atrophy, 5) cortical tubulointerstitium with fibrosis or atrophy, 6) cortical tubulointerstitium with inflammation, 7) medullary ray or medulla. These categories were used to evaluate niche group associations, transcription factor activity, and gene expression related to soluble biomarkers.

**FUSION interactive tool for histology-Visium data navigation.** The digital histopathology slide stained with hematoxylin and eosin, and associated cell type and state abundance data for each spot obtained using cell deconvolution method^1^ applied on the raw 10X Visium data is available via our recently published FUSION tool https://fusion.hubmapconsortium.org/. Data can be visualized via the *Visualization* tab for various imaging pathomics and cell type and state comparison analysis for various pathology within the digital pathology images, and selected and populated in batches via the *Dataset-Builder* tab, and for detailed method on data navigation for pathomics and omics fusion, see our FUSION work^67^. Total *n* = 150 cases from CKD, AKI, and reference are made available via FUSION.

### CosMx (Nanostring) Spatial Transcriptomics

<u>Tissue processing and imaging</u>. CosMx spatial transcriptomics assay was performed on FFPE biopsy tissue sections (5 um) from patients with diabetic nephropathy (n = 3) on a CosMx SMI system under the technology access program by Nanostring technologies. The sample processing, staining, imaging, and cell segmentation were performed as previously described ^119^. Briefly, 5 µm human FFPE kidney tissues were sectioned to VWR Superfrost Plus Micro Slide (Cat# 48311-703) for optimal adherence. Slides were then dried at 37°C overnight, followed by deparaffinization, antigen retrieval and proteinase-K mediated permeabilization dedicated to CosMx sample preparation (https://university.nanostring.com/cosmx-smi-manual-slide-prep-for-rna-assays). 1 nM RNA-ISH probes (human 1k-plex RNA panel plus 29 customized spike-ins) (**Supplementary Table S5)** were applied for hybridization at 37°C overnight. After stringent wash, a flow cell was assembled on top of the slide and cyclic RNA readout on CosMx was performed (a development-stage Alpha version of CosMx instrument was used for this experiment). 20-30 0.753mm × 0.753mm fields of view (FOVs) were placed for data collection in each sample. After all imaging cycles were completed, additional visualization markers for morphology and cell segmentation were added including pan-cytokeratin, CD45, CD3, CD298/B2M, and DAPI. The Alpha-version of CosMx optical system implemented an epifluorescent configuration with a customized water objective (13×, NA 0.82), and widefield illumination, with a mix of lasers and light-emitting diodes (385 nm, 488 nm, 530 nm, 590 nm, 647 nm). A scientific CMOS camera was used for signal detection (pixel size 168nm, Sony). A 3D multichannel image stack (9 z-frames) was obtained at each FOV location, with the step size of 0.6 um. The fluidic system uses a custom interface to draw reagents through the flow cell with a syringe pump. Reagent selection is controlled by a shear valve (Idex Health & Science). A flow sensor between the flow cell and syringe pump was used for flow rate feedback (Sensirion AG). The fluidic interface includes a flat aluminum plate in direct contact with the flow cell. The metal plate temperature was controlled to regulate the reporter hybridization temperature. The enclosure around the instrument was also maintained at a constant temperature using a separate thermoelectric cooler. Image registration, feature extraction, localization, decoding of individual transcripts, and machine-learning based multimodal cell segmentation (developed upon Cellpose ^120^) were performed as previously established ^119^. The final segmentation mapped each transcript in the registered images to the corresponding cell, as well as to subcellular compartments (nuclei, cytoplasm, membrane), where the transcript was located. We added 29 custom genes to a 1k research panel to increase detection of kidney specific cell types and injury markers.

<u>Cell type deconvolutions</u>. CosMx data analysis was performed using Seurat v5.0 and deconvolution of cell types was performed using RCTD from the spacexr package (version 2.2.1, github.com/dmcable/spacexr). Broad cell types (subclass level 1) were first predicted on each sample separately as outlined for Slide-seq2 above. Count matrices were combined across samples and clustered using Pagoda2. The resultant clusters and embeddings were transferred to the Seurat object and annotated to broad subclasses (level 1) based on RCTD predictions, correlation with the reference atlas sub-clusters and expression of marker genes. Clusters that were poorly predicted or showed too low marker gene expression were labeled as ambiguous and removed before repeating the clustering and annotation process. Additional ambiguous clusters from the second round of clustering were also identified and removed. This enabled more reliable level 1 subclass annotations which were then used as the basis for subsequent more resolved cell type predictions (v2.subclass.sp, **Supplementary Table 14**). For this, broad groupings of proximal to intermediate tubules, distal tubules, collecting ducts and non-epithelial cells were independently used for the corresponding level 2 RCTD predictions. Reference objects were subset to remove degenerative and cycling cell states, then downsampled to 5000 nuclei per subclass (v2.subclass.sp). Prediction weights were again scaled to 100 and stored as a separate assay within the Seurat object. Final annotations were assigned from independent clustering of each broad grouping as described for subclass level 1. Accuracy of final predictions was assessed through: correlation of averaged scaled expression values with reference single nucleus RNA subclasses visualized using the corrplot r package; marker gene expression visualized using the DotPlot function in Seurat; correct spatial enrichment (normalized cell type fraction per field of view) in cortex, outer medulla or inner medulla regions (assigned to tissue pucks based on histological sections) visualized using the ggplot2 r package (v3.5.1). Spatial mapping of molecules and cell types was performed in Seurat using the ImageDimPlot function.

### Xenium (10X Genomics) Spatial Transcriptomics

<u>Xenium Sample Preparation and Imaging</u>: We sectioned FFPE tissue at 5 µm thickness onto a 10x Xenium slide to be processed according to manufacturer protocol CG000601 for Xenium *In Situ* Gene Expression. We performed independent Xenium experiments using two different custom probe panels at different institutions (WashU and IU) for validation (**Supplemental Table 6)**. The WashU panel, 300KID, was used for validation of altered PT and TAL and microenvironment of SOX4 / SOX9 cells and neighborhood analysis. The IU 300 custom gene panel was used to localize cell-cell interactions between altered epithelial or stromal cells and target immune cells ^121^. We obtained H&E-stained histological images after the Xenium protocol completion using Hamamatsu whole slide image scanning (300KID experiments) as described above or an EVOS M7000 with Olympus 40x objective (IU). Histological and DAPI-stained nuclei images were co-registered with Xenium Explorer 2.0.0.

<u>Xenium Cell Type Annotation</u>. Xenium data analysis was performed using scanpy(version 1.9.5) and squidpy (version 1.3.1) packages, and deconvolution of cell types was performed using Transfer of Annotations to Cells and their COmbinations (TACCO (version 1.0.0); simonwm/tacco) **(Supplementary Table S6).** For each sample, AnnData objects were generated incorporating the count matrix, cell and gene metadata, and spatial coordinates. After calculating quality control metrics, cells with fewer than 5 total gene counts and genes not detected in any cell were excluded. The data were then log-transformed, normalized, and subjected to principal component analysis. A neighborhood graph was constructed and used for clustering. Broad cell types (subclass level 1) were first predicted on each sample, and then each broad category was subclustered and annotated into further granular subtypes (subclass level 2) using the snRNA-seq annotations. The samples were integrated using reSOLVI from scvi-tools package (version 1.3.0) To ensure accurate cell type annotation, we collapsed the subclass level 2 into a custom level that aligns better with the panel’s resolution, making it less granular than level 2, but more detailed than level 1.

<u>Xenium PT Analysis</u>: Injured tubule cell states are challenging to annotate in ST compared to the granularity that can be achieved in dissociated transcriptome-wide single-cell RNA sequencing methods, as the injured cells start losing differentiation markers, and custom panels are limited in size. We used the TACCO-predicted annotations using our HKAv2 and aligned them to the transcript files for each sample, retaining only the proximal tubules for downstream analysis. Our focus was on quantifying gene expression within the PT and altered PT populations. All other cells were excluded. We then focused on quantifying gene expression within PT and altered PT populations. The fraction of SOX4 or PCK1 (> 2 transcript counts in a cell) PT for each sample was computed, and the indicated comparative analyses were done using a two-tailed t-test (significance p < 0.05) and represented as box plots.

<u>Xenium Neighborhood Analyses</u>: To identify spatially organized cellular niches, we performed neighborhood clustering using BANKSY (Bayesian Analysis of Neighborhoods for Spatial Systems) ^122^. We used BANKSY (banksy-py 0.0.7) to construct spatial expression graphs using with lambda = 0.5, weighting spatial and transcriptional distances equally. Principal component analysis was performed using the top 20 components and neighborhood clusters were defined with a resolution parameter of 0.7. Each resulting BANKSY cluster corresponds to a spatially coherent microenvironment, capturing localized patterns of cell-cell organization. We used our manually curated annotations to reveal and quantify the distribution of known cell types within each neighborhood, and visualized cluster composition using spatial plots and stacked barplots. This analysis enabled us to detect shared and sample-specific tissue niches, as well as spatial shifts in cell-type composition across conditions.

<u>Pathomic Features correlated with Xenium Niches</u>. Pathomic features were extracted from the H&E post-stained sections used in the Xenium experiments, using the Scikit-image python package (version 0.24.0). Nuclear contours defined in the Xenium segmentation were converted to binary spatial maps and registered to hematoxylin segmentations from the H&E section using an affine transformation. Both nuclear and cell boundaries defined in the Xenium analysis were registered using the measured transformation matrix, and features were measured from the brightfield sections. Features included color, shape, and texture, and were Z-normalized per section, across all cell types. Aspect ratio is defined as the major axis divided by the minor axis. Solidity is the area of the object divided by the area of the convex hull, and shape factor is defined as the area divided by the perimeter of the object.

### Spatial Metabolomics

We performed untargeted MALDI-MSI on frozen kidney biopsy sections from nine KPMP participants with diabetes to profile metabolites in individual renal tubules. Tissue was sectioned at 7 µm on a Leica CM1950 cryostat and mounted on either indium tin oxide (ITO) slides for MSI or standard slides for PAS-H staining. A 1,5-diaminonaphthalene (DAN) matrix (12.5 mg/mL in 50% EtOH) was applied to ITO slides using an HTX M3+ sprayer (20 µL/min; 16 passes; 10 psi LN_2_ sheath gas; 1200 mm/min). MALDI-MSI was performed on a Thermo Q Exactive HF-X Orbitrap with a Spectroglyph UV-laser MALDI source in negative ion mode (m/z 100–1000; 20 µm resolution). Data were converted to .imzML using ImageInsight and annotated in METASPACE with HMDB, KEGG, CoreMetabolome, and SwissLipids databases (≤ 20% FDR). Serial sections were formalin-fixed (4% formaldehyde), and PAS-H stained. Autofluorescence and brightfield images (Zeiss Axioscan 7, 20x) were exported as .tiff files and overlaid with MSI data for metabolite localization. Tubules were annotated in QuPath (v0.5.0) as atrophic or non-atrophic, and images were registered to MSI data in SCiLS (Bruker). Metabolite intensities were normalized to total ion count (TIC) and further normalized to the percentage of non-atrophic tubules to reduce batch effects. Statistical analysis was performed in MetaboAnalyst v6.0 ^123^. Volcano plots were generated using a fold change ≥ 1.2 and FDR < 0.05. Isomers were assigned a single name based on literature and an in-house MALDI-MS/MS and LC-MS/MS library; duplicates from multiple adducts were removed. Regulated metabolites were subjected to pathway analysis in MetaboAnalyst using SMPDB and KEGG to identify dysregulated pathways in atrophic tubules.

For the glycolysis and gluconeogenesis panel, TIC-normalized intensities of targeted metabolites were analyzed using MetaboAnalyst. Missing values were imputed with 1/5 of the lowest intensity of the corresponding metabolite across samples. The data were log-transformed and autoscaled (mean-centered and divided by the standard deviation of each variable). The heatmap was generated using the “autoscale samples” standardization option, Euclidean distance measurement, and Ward’s clustering method. Values from the .json file were then plotted in GraphPad Prism 10 to create the final heatmap presented in the figure.

### Imaging Mass Spectrometry (Spatial Lipidomics)

The complete VU TIS multimodal molecular imaging workflow ^124^ is summarized here (protocols.io https://dx.doi.org/10.17504/protocols.io.kqdg39bbeg25/v2); relevant metadata is summarized in **Supplementary Table 8**.

<u>Sample Preparation</u>: Biopsies were cryo-sectioned to 10 µm thickness using a CM3050 S cryostat (Leica Biosystems, Wetzlar, Germany) and mounted onto ITO-coated glass slides. Autofluorescence microscopy was acquired of each section using DAPI, eGFP, and DSRed fluorescent filters on a Zeiss AxioScan.Z1 slide scanner (Carl Zeiss Microscopy GmbH, Oberkochen, Germany), equipped with a Colibri7 LED light source. Tissue sections were then washed with chilled (4 °C) 150 mM ammonium formate (3 times for 45 seconds) and dried with nitrogen gas. An in-house developed sublimation device was used to sublime 4-(dimethylamino)cinnamic acid (DMACA) onto the slide (matrix density ∼0.22 µg/mm^2^). Briefly, the apparatus was heated (190 °C) for 10 minutes under vacuum (110-150 mTorr) while the sample was cooled to ∼80°C using a dry ice and acetone slurry in the cold finger.

<u>IMS Data Collection</u>: Matrix-assisted laser desorption/ionization (MALDI) IMS data acquisition was performed on a timsTOF fleX mass spectrometer (Bruker Daltonik, Bremen, Germany) ^125^. Tissue imaging data were collected in negative ion mode (*m/z* 400 - 2000) at a pixel size of 10 µm × 10 µm, 150 shots per pixel, and ∼30% relative laser power. LC-MS/MS was collected on a serial tissue section to aid in lipid analyte identification. Prior to matrix removal, post-IMS autofluorescence images were acquired using a Zeiss AxioScan.Z1 fluorescence slide scanner using eGFP fluorescence filter and a brightfield image.

<u>Histology & Feature Annotation</u>: Matrix was removed from the tissue slides using a series of ethanol washes, and the samples were stained using PAS. The PAS stains were annotated by a trained pathologist to mark regions of interest, including arteries/arterioles, Tamm-Horsfall protein cast, glomeruli, IFTA, and gross morphological regions including cortex and medulla. Annotations were made in *QuPath*, and the regions of interest were exported in *GeoJSON* file format.

<u>Multimodal Image Registration and Data Analysis</u>: First, MALDI IMS data pre-processing, including peak alignment, data calibration, and normalization, was carried out. MALDI IMS and microscopy images were then co-registered using an in-house developed software *wsireg* and *image2image.* Registered images were stored in the vendor-neutral pyramidal OME-TIFF format at their original spatial resolution (i.e., no downsampling or changes to pixel spacing through the registration process). The *GeoJSON* files (manual feature annotations described above) were transformed to the IMS coordinate system using an affine transformation matrix that was created based on fiducial markers selected from the post-IMS AF and IMS modalities. The transformed files were used as masks to select and extract MALDI IMS pixels associated with each annotation for further data analysis.

<u>Supervised Machine Learning and Shapley Additive Explanations</u>: Manual annotations of 6 classes were used to build eXtreme Gradient Boosting (XGBoost) ^126^ classification models that recognize one of the available classes. We take the one-versus-all approach to multiclass classification: each classification task differentiates the positive class against all the other pixels in the dataset (imbalanced binary classification). Shapley additive explanations (SHAP) ^127,128^ enable us to determine which ion species have a marker-like relationship to each class by quantifying each ion species’ importance to the corresponding classification models. For a given classification task, SHAP measures the global (experiment-wide) and local (per-pixel) relevance (importance) of each ion species. The global SHAP importance scores rank all *m/z* species by decreasing relevance for recognizing a specific class, highlighting a set of highly discriminative molecular species that represent potential biomarker candidates. Conversely, local SHAP importance scores assess the direction of relevance (positive or negative monotonic correlation) and assess the significance of the relationship between an ion species’ intensity and a pixel’s likelihood of belonging to a class. In the summary bubble plots (**ED Figure 10b**), the size of each marker corresponds to the global SHAP importance score of a given molecular species for a given classification task (recognition of a single class); the marker color corresponds to the Spearman rank-order correlation coefficient per molecular species (column) and per histological feature (row).

<u>Spatial N-glycomics analysis</u>: Step-by-step details of the tissue preparation and PNGase F application method can be found in protocols.io: dx.doi.org/10.17504/protocols.io.8epv5j1m4l1b/v. Briefly, FFPE blocks of human kidney biopsies were sectioned at 7 µm thickness and mounted on indium tin oxide (ITO)-coated glass slides. Slides were heated, dewaxed by xylene washes, and rehydrated in serial ethanol (EtOH)/water (v/v) washings and then subjected to antigen retrieval in boiling citraconic buffer followed by PNGase F (N-Zyme Scientifics, 100 µg/mL) spraying using a M5 Sprayer (HTX Technologies), and sample incubation in a relative humidity of 89% for 2 h at 37 °C, as described previously ^129^.

After incubation, α-cyano-4-hydroxycinnamic acid (CHCA, Sigma-Aldrich)– 7 mg/mL (50% ACN and 0.1% TFA in water (v/v))– was sprayed over the tissue sections using the M5 Sprayer, as described previously ^129^. MALDI-MSI experiments were performed using a scimaX 7 Tesla Magnetic Resonance Mass Spectrometer (MRMS; Bruker Daltonics) equipped with a dual ESI/MALDI ion source and a Smart-beam II Nd:YAG (355 nm) laser. The instrument was operated in 1 w, positive ion mode over an m/z range of 1,000–5,000 with an estimated resolving power of 120,000 at m/z 400. The target plate stepping distance (lateral resolution) was 50 μm. The ion m/z 1809.6393 ([M+Na]+ of Hex5 dHex1 HexNAc4) was used as a lock mass for on-line calibration. Imaging data were acquired using FlexImaging (v 4.1, Bruker Daltonics).

Imaging data files were imported into the SCiLS software (Version 2025b), exported to imzML, and the resulting .imzML and .ibd files were then submitted to METASPACE for data processing using the NGlycDB-V1 as the database ^130^. Molecular annotations from NGlycDB-V1 (FDR ≤ 20%) were imported into SCiLs and on tissue regions were segmented. The data was exported to R using the SCiLS API, and data processing was performed in the open-source R package *ROmicsProcessor (v1.1.6)*. Batch effect was minimized by normalizing ion intensities in tissue to a reference tissue section on the same slide. Then the data was then log_2_ transformed and centered at zero. A two-tailed Student’s t-test was performed, and log_2_-fold change was computed between AKI, CKD, HRT samples.

### Confocal immunofluorescence microscopy

Immunofluorescent (IF) staining of MID1 (Rabbit, AB70770), or PXDN (Rabbit, NBP3-47384), on FFPE sections of human kidney was performed on replicate sections from 2 individuals **(Supplemental Table 7)** as previously described ^1^. Briefly, heat mediated antigen retrieval was performed in 10 mM citrate buffer. After blocking with 1% BSA with 0.2% skim milk and 0.3% triton X-100 in 1X PBS, and blocking buffers specific for biotin and streptavidin, the tissue sections were incubated with the primary antibodies and biotinylated-LTL at 1:100 (LTL 1:200) dilution overnight at 4°C, followed by labeling with secondary antibodies (anti-rabbit-Alexa-488 and Alexa-594-streptavidin). Finally, sections were stained with DAPI (1:500) for 5 min to label all cell nuclei.

### CODEX Multiplexed Imaging

Five-micron sections from FFPE blocks were cut onto Superfrost Plus Gold Charged slides. Staining of the slide after deparaffinization followed the recommended protocol provided by Akoya Biosciences with minor adjustments ^131^. Targets shown are listed in **Supplementary Table 7**. Imaging of the tissue was conducted with a 20x objective fitted on the Akoya Biosciences Phenocycler-Fusion 2.0 microscope and fluidics handler. Image stitching and processing was also performed with the Phenocycler-Fusion (v2.0) software.

### Translational Ribosome Affinity Profiling (TRAP) analysis for Sox4, Gdf15

We analysed raw data from TRAP studies of mouse IRI models available from Liu et al ^132^ to show translatomic responses for both Sox4 and Gdf15. Nephron-specific normalized TRAP microarray probe intensities were used to calculate fold changes in expression levels. We applied Tukey’s One-Way ANOVA multiple comparisons test (*p-value < 0.05; **p < 0.01 was defined statistically significant) for statistical significance.

### Disease Models and Clinical Impact Experiments

<u>In vitro gene knockdown</u>. Normal human proximal tubular kidney (NHPTK) cells ^133^ were plated at a density of 4.5 × 104 in a 24-well flat-bottom plate and maintained in Renal Epithelial Growth media (REGM, Lonza, Basel, Switzerland) and 9% fetal bovine serum (HyClone). Cells were diluted to 20–30% confluency three times a week and maintained at 37 °C in 95% humidified atmosphere with 5% CO2. We conducted siRNA-mediated knockdown of *SOX4* in NHPTK cells with lipofectamine. Cells were plated on Day 1 with lipofectamine and a pool of up to 3 directed siRNA constructs (20 nM concentration each) for 2 conditions: (1) Silencer (Cat. No. 4390843), scrambled siRNA control (Cat. No. 4427037), (2) a pool of three *SOX4* siRNA molecules (siRNA ID s13300, s224666, and s13301). Total mRNA expression was measured 48 hours after siRNA or scrambled control transfection. RNA was isolated with the miRNeasy Mini Kit (Cat. No. 217004, Qiagen, Hilden, Germany) and converted to cDNA with the High-Capacity cDNA Reverse Transcription Kit (Cat. No. 4368814, Thermo Fisher) according to the manufacturer’s protocol. Total mRNA was sequenced on an Illumina Next-seq X PLUS, reads were mapped to hg38 with STAR (2.7.10a), and counts were obtained with subread (2.0.3).

<u>Impact of SGLT2i treatment in patients with diabetes</u>. Research Kidney biopsy from adolescents and young adults (N = 16) with youth-onset T2D (12–21 years of age, T2D onset at < 18 years of age, diabetes duration 1–10 years, and HbA1c <11%) from the Renal-HEIR (ClinicalTrials.govIdentifier: NCT03584217) and the IMPROVE-T2D study (ClinicalTrials.gov Identifier: NCT03620773) were included in this analysis. -seq analysis was performed on cell populations obtained from kidney tissue samples of 10 patients treated with an SGLT2i (T2D(+)), 6 patients under standard care(T2D(-)), and 6 healthy reference tissues (HC) from the CROCODILE study. Tissue processing, single-cell isolation, and scRNA-seq data generation were performed according to the protocol developed for the Kidney Precision Medicine Project (dx.doi.org/10.17504/protocols.io.7dthi6n). Details of the scRNA-seq analysis across all kidney cell types were reported previously ^91^.

Reversal pattern of genes upon SGLT2i treatment were defined if the genes were significantly regulated in T2D(-) compared to HC and showed a reversal pattern when compared to the T2D(+) ^91^.We specifically focused on the GRN networks for the TAL trajectory of cell states to define the RNA expression and regulation patterns of downstream targets upon SGLT2i treatment. The downstream gene sets for each TAL trajectory TF were derived from the S**upplementary Table 21**. These genes were mapped to the differentially reversed genes in the TAL nephron segment as mentioned. We found several segments of TF GRN networks to be differentially reversed with the SGLT2i suggestive of failed repair and differentiation as well.

### Cell State-predicted soluble markers associated with Clinical Outcomes in AKI or CKD patients

<u>Validation of state-associated soluble markers in four AKI cohorts</u>. We examined a set of 626 genes that predicted secreted markers from trajectory associated early, mid and failed repair PT, TAL, FIB and MAC cell types (**Supplementary table 26),** and quantified the association of the corresponding proteins with kidney-related outcomes in existing cohorts (TRIBE-AKI, ASSESS-AKI, NAIKID and Hopkins Health Reference).

<u>TRIBE-AKI Study</u>. The TRIBE-AKI study is a prospective study of adults undergoing cardiac surgery who were at high risk for post-operative AKI ^81^. Blood samples were collected within 6 hours post-operatively. In a subset of **784** participants, protein analytes were quantified using a multiplexed modified DNA-based aptamer technology (SomaScan assay). The SomaScan v4 assay included 5,482 aptamers that mapped to 4,746 unique proteins in the UniProt database ^134^. AKI was defined as an increase of serum creatinine concentration of 0.3 mg/dL or more, or at least 50% from the pre-operative serum creatinine value. In a subset of patients (**363**) that experienced AKI (AKI stage 1 or higher), we examined the outcome of AKI progression defined by worsening of AKI stage (from stage 1 to either 2 or 3, from stage 2 to 3, or beginning with stages 2 or 3). To account for multiple comparisons, Benjamini-Hochberg procedure was applied to p-values from logistic regression models for AKI (adjusting for age, sex, diabetes, and baseline eGFR) (**Supplementary Table S27)**. Only the proteins significantly associated with AKI were considered for the secondary outcome of AKI progression.

<u>ASSESS-AKI Study</u>. The ASSESS-AKI study is a prospective cohort of 1538 hospitalized patients with and without AKI (1:1 matching) ^135^. Urine samples were collected 3 months post-discharge. Follow-up study visits were conducted annually with telephone visit every 6 months. In a subset of **174** participants that did (n=87) and did not (n=87) develop CKD progression, urine proteins were measured by Olink Explore 3,072 platform with 2,783 proteins passing all QC checks. To account for protein variations by participant factors, multivariable linear regression models were used to regress out age, sex, and urine creatinine. Residuals from linear regression models were considered adjusted protein levels and used for all subsequent analyses (in all cohorts with Olink measurements). Linear mixed-effects models with random intercepts and slopes were used to determine the associations between adjusted protein levels and longitudinal decline in eGFR. Models adjusted for baseline CKD status prior to hospitalization, AKI status during hospitalization, and EGFR, heart failure, diabetes, and hypertension at 3 months, time and time-protein interaction. We reported the change in eGFR decline rate (percentage per year) per doubling of protein concentration (**Supplementary Table S27)**.

<u>NAIKID Cohort and Hopkins Healthy Reference Cohort</u>. The NAIKID study is an ongoing prospective study of participants with a clinically indicated native kidney biopsy at the Johns Hopkins Hospital. Urine samples were collected at the time of biopsy. The Hopkins Healthy Reference Cohort is a prospective study of healthy volunteers with no known history of chronic disease between the ages of 18 to 80. Participants provided a urine same at the time of the study visit. In a subset of NAIKID participants with confirmed acute tubular injury (**29** participants) and participants from the Hopkins Healthy Reference Cohort (**75** participants), urine proteins were measured by Olink Explore 3,072 and followed the same data processing and linear regression analysis as described for the ASSESS-AKI cohort. Logistic regression models were used to examine the association of adjusted protein levels with ATI and healthy reference. Only proteins significantly associated with AKI in the TRIBE-AKI cohort were examined in the NAIKID and Hopkins Healthy Reference Cohorts without adjustment for multiple comparisons (**Supplementary Table S27)**.

<u>The Boston Kidney Biopsy Cohort (BKBC)</u>. The BKBC is a longitudinal, observational study of adult patients who underwent native kidney biopsies for clinical indications at three academic medical centers in Boston, Massachusetts between September 2006 and October 2018. Individuals were excluded if they were unable to provide written informed consent, were pregnant, had significant anemia, or were participating in conflicting studies. The study protocol and methodology have been detailed previously ^136^. The Mass General Brigham institutional review board approved the study protocol (protocol #2012P000992). Blood samples were collected on the day of biopsy. For this analysis, we included 418 participants with plasma proteomic profiles measured using the SOMAscan platform as previously described using a set of 582 early repair genes not previously reported in ARIC, CRIC and BKBC cohorts (**Supplementary Table 26)** ^53,96^. The primary outcome was progression to end-stage kidney disease (ESKD), defined as initiation of dialysis or kidney transplantation. The occurrence of ESKD was determined through electronic medical records review and linkage with the United States Renal Data System. *Statistical analysis*: Categorical variables were described as frequencies with percentages, and continuous variables as means ± standard deviation or medians with interquartile ranges. We used Cox proportional hazards models to evaluate associations between the proteins of interest and ESKD. The adjusted model included the covariates age, race, sex, eGFR, and proteinuria. A prespecified α level of 1.05 × 10⁻⁴ set by Bonferroni correction (0.05/478 proteins) was used to determine statistical significance **(Supplementary Table S28).** Pathways associated with the 82 significant proteins were determined using ENRICHR (https://maayanlab.cloud/Enrichr/enrich?dataset=72187ea56d63c403b20b1b35f3903ba1).

<u>AKI-CKD recovery and progression groups in KPMP AKI participants.</u> snRNAseq data from biopsies at enrollment of 26 KPMP AKI patients with follow up data were used identifying patients who recovered or progressed in disease severity. To define CKD incidence or progression, any available follow-up serum creatinine values up to 18 months excluding measurements from the Baseline, Enrollment or AKI-only study visits. Baseline CKD was defined as the baseline eGFR (variable sc_egfr_aki_blC) <60 mL/min per 1.73m2. The ASSESS-AKI Consortium CKD Incidence or progression follow-up definition was applied. For participants without CKD, CKD incidence was defined as at least a 25% reduction in eGFR and a fall below 60 mL/min per 1.73m2 ^135^. For participants with CKD, CKD progression was defined as at least a 50% reduction in eGFR or a fall below 15 mL/min per 1.73m2. Dialysis and death information was not provided and therefore is not included in this definition. When available, all outpatient serum creatinine values were considered first, followed by the last available inpatient or unknown source serum creatinine.

## Acknowledgements

We thank Dr. Ying Maggie Chen for her advice on thick ascending limb biology, Amy McMurray and Kristy Conlon and the WashU Kidney Translational Research Center (KTRC) in part for supporting regulatory approvals, tissue procurement and processing and Mid America Transplant in St Louis for infrastructural support for HuBMAP samples, Drs. Nir Hacohen and Deepak Rao for help with the immune cell type annotations, Yi Cui from nanostring technologies for assistance with the technology access program data curation and generation and Karol Balderama for assistance with SlideSeq2 data. The authors acknowledge the University of Michigan Medical School Central Biorepository (RRID:SCR_026845) for providing biospecimen storage, management, and distribution services in support of the research reported in this publication. For computational analysis, the authors acknowledge support by the state of Baden-Württemberg through bwHPC and the German Research Foundation (DFG) through grant INST 35/1597-1 FUGG, as well as the data storage service SDS@hd supported by the Ministry of Science, Research and the Arts Baden-Württemberg (MWK) and the German Research Foundation (DFG) through grant INST 35/1503-1 FUGG and Paul Teschan Research fund NO 2024-01 (RM-F). We are grateful to HuBMAP HIVE Pittsburg data center for assistance with data ingestion and availability for the community and CZI CellxGene team for visualization tools for the atlas.

The Kidney Precision Medicine Project (KPMP) is supported by the National Institute of Diabetes and Digestive and Kidney Diseases (NIDDK) through the following grants: U01DK133081 (JPL, ACR), U01DK133091 (MV, RT), U01DK133092 (SW, SR), U01DK133093 (SGC, GNN), U01DK133095 (AM, PRC, FCB, BT), U01DK133097 (PHN, MLC), U01DK114866 (FPW, CP), U01DK114908 (EP, JO, JS), U01DK133090 (JH, MK), U01DK133113 (JT, WYEH), U01DK133766 (JMS), U01DK133768 (VR, RK, LC), U01DK114907 (MK, JH, NH, OT), U01DK114920 (KS, TA, CA), U01DK114923 (PCD, TA, ME), U01DK114933 (SJ), U24DK114886 (JH, MK), UH3 DK114907 (MK), UH3DK114926 (KK, JB, AB), UH3DK114861 (PP, MR, RM),UH3DK114915 (SW), UH3DK114937 (ZL). We are grateful for the NIH Common Fund supported Human Biomolecular Atlas Program (HuBMAP) grants U54DK134301 (TMA and SJ), OT2OD033753 (PS and SJ) and U54DK13430 (JMS), NIH/NIDDK K23DK116720, as well as Boettcher Foundation (PB), K01DK136973 (IMS), U2CDK114886 opportunity pool and RO1DK129879 (MR), U54DK083912 (MK), P30DK081943 and Human Cell Atlas Kidney Seed Network (MK), R01DK108803 (MEG), P30DK079312 (PCD), P01DK056788 (JCW), R01DK113191, P30DK079310 (SK), R01DK118265 (JAS).

We gratefully acknowledge the essential contributions of our patient participants and the support of the American public through their tax dollars.

The content is solely the responsibility of the authors and does not necessarily represent the official views of the National Institutes of Health.

## Author contributions

First draft: B.B.L., M.T.E., S.J.; Contribution to Assay prep: R.M.F., B.Z, A.L.K, E.A.O, K.Y.C; Contribution to scRNA-seq data generation or analysis: R.M, E.A.O, K.Z, J.B.H; Contribution to snRNA-seq data generation or analysis: B.B.L, A.L.K, M.Ka, E.C, P.V.K, K.Z, M.K, S.J; Contribution to Multiome data generation or analysis: B.B.L, J.B, X.C, D.L.G, B.Z, A.L.K, M.Ka, R.S.S, P.C.D, O.G.T, K.Z, S.J; Contribution to Visium data generation or analysis: R.M.F., R.F, Y.C, P.C.D, M.T.E; Contribution to Visium FFPE data generation or analysis: R.M.F., Y.C, M.T.E; Contribution to CUT& RUN data generation and analysis: J.B, M.B, L.R, M.R; Contribution to CosMx data generation and analysis: B.B.L, S.R, A.K.L, P.C.D, S.J; Contribution to Xenium data generation and analysis: B.B.L, R.M.F., S.R, A.K.L, Y.C, M.T.E, S.J; Contribution to SlideSeqv2 data generation or analysis: B.B.L, E.C, E.Z.M, S.J; Contribution to Spatial metabolomics data generation and analysis: S.Ma, K.V.D, B.L.G, M.A.F, D.V, I.T, C.R.A, J.M.S, K.S, J.B.H; Contribution to CODEX or MxIF data generation and analysis: B.Z, A.R.S, P.C.D, T.M.E, S.J; Contribution to Mouse injury models: F.A, E.K, S.K, S.J; Contribution to mouse aging models: B.B.L; Contribution to mouse atlas: B.B.L, F.A, S.J; Contribution to in vitro and in silico validation: R.M.F., D.L.G, Y.C, M.T.E; Contribution to Data curation and deposition: B.B.L, R.M.F., R.M, A.K.L, K.V.D, A.L.K, M.Ka, M.A.F, D.D, Y.C, N.B, J.M.S, M.K, M.T.E, S.J; Contribution to Patient recruitment or samples: Y.C, F.C.B, M.L.C, S.G.C, R.S.F, E.H.K, K.K, J.F.O, P.M.P, E.P.R, A.C.R, S.E.R, P.R, M.M.S, J.R.S, R.D.T, A.T, S.S.W, J.C.W, F.P.W, E.S.W, M.T.E, S.J; Contribution to Pathology descriptor scoring or evaluation: C.E.A, U.G.B, L.B, D.De, A.B.F, J.M.H, L.H, G.W.M, P.S.R, A.Z.R, S.S, J.P.G, J.B.H; Contribution to Clinical adjudication: I.M.S, C.E.A, L.B, I.H.B., J.M.H, L.H, G.W.M, A.Z.R, J.A.S, S.S, S.G.C, K.K, J.P.L, R.T.M, P.M.P, S.E.R, F.P.W, T.M.E, J.H, M.K, J.B.H, C.R.P; Contribution to Clinical biomarker outcomes: H.T, I.M.S, A.S, S.M, I.H.B., J.P.L, M.E.G, C.R.P, S.J; Contribution to AKI progression: H.T, C.R.P, S.J; Contribution to Molecular-pathomics integration: B.B.L, R.M.F., N.L, P.S, M.T.E, S.J; Contribution to pathway analysis: B.B.L, J.Ha, R.M, R.I, S.J; Contribution to Drug targets: S.R, F.A, S.J; Contribution to Ligand-Receptor data and analysi: B.B.L, R.M.F., R.M, R.F, J.T, J.S., M.T.E, S.J; Contribution to Diabetic models: V.N, F.A, P.B, M.K; Contribution to Ontology: B.B.L, R.M, Y.H, N.B, M.K, S.J. Reviewed or edited the manuscript: all authors. Other consortium collaborative efforts: K.P.M.P; Led the study: S.J.

## Competing interests

B.B.L., D.D., K.Y.C. are and E.C. and P.V.K. were full-time employees of Altos Labs Inc. L.B. consults for Sangamo, Protalix, Uniquire, and Idorsia; is on scientific advisory boards for Vertex and Nephcure; serves as a grant reviewer for ASN; is on the editorial board of the Journal of Glomerular Diseases. J.M.H. has research funding from: Evotec, Novo Nordisk, Pfizer, Visterra; consults for Novartis. M.L.C. has consultant fees from Armana, Bayer AG, Bayer Pharmaceuticals, Novo Nordisk; has royalties from Up-to-Date; has research grants from Breakthrough T1D, Boehringer Ingelheim, Eli Lilly, Bayer Pharmaceuticals (all paid to institution). P.M.P. did prior consulting for Chiesi USA. S.E.R. receives research funding from Bayer, Astra Zeneca (paid to institution), serves on Steering Committees for FineOne; serves on Scientific Advisory Board for Bayer, AstraZeneca, Travere, Novo Nordisk; is employed by Joslin Diabetes Center at Beth Israel Deaconess Medical Center; Immediate Past President of the National Kidney Foundation. J.R.S. consults for Maze and Goldfinch and receives royalties from Sanofi Genzyme. S.S.W. serves as an expert witness consulting on patent issues, dialysis laboratory testing, and drug safety (Dechert, DLA Piper, Finnegan, Ropes and Gray, Tucker Ellis); consults for industry/pharma on drug development, safety, etc. (Aditum - Motric, Bain, CANbridge, Dechert, Delix, Goldfinch, Ikena, Merck, Mineralys, Ono Pharma, PepGen, Quinn Emanuel, Strataca, Vertex); receives grants to institution (Vertex, Pfizer, J&J, Natera). F.P.W. receives grant support from AHRQ, DOD, Amgen, AstraZeneca; consults for Whoop, WndrHlth. E.Z.M. is a paid consultant for Atlas Bio. J.S. reports in the last 3 years funding from GSK and Pfizer and fees/honoraria from Travere Therapeutics, Stadapharm, Astex, Owkin, Pfizer, Grunenthal, Tempus and Moderna. N.B. owns equity interest in Thermo Fisher Scientific. P.S. serves on the advisory board of DigPath Inc. FUSION software is protected by ©Copyright 2023-25 University of Florida Research Foundation, Inc. K.Z is a full-time employee of Altos Labs Inc.; Co-founder, equity holder and serves on the scientific advisory board of Singlera Genomics. M.K. reports grants and contracts through the University of Michigan outside of this work from AstraZeneca, NovoNordisk, Eli Lilly, Boehringer-Ingelheim, European Union Innovative Medicine Initiative, Certa Therapeutics, RenalytixAI, Regeneron, Novo Nordisk, Sanofi, Dimerix, Travere and Vera Therapeutics; has received consulting fees through the University of Michigan from Novo Nordisk, Alexion, Novartis, Roche Diagnostics and Vera Therapeutics; with V.N. has a patent PCT/EP2014/073413 “Biomarkers and methods for progression prediction for chronic kidney disease” licensed. C.R.P. is a member of the advisory board of and owns equity in RenalytixAI, and serves as a consultant for Genfit and Novartis. S.J. has consulted with Athenium, receives royalties from Elsevier and with P.S. and T.M.A. has an intellectual property invention disclosure on FUSION histology-omics software tool and may receive royalties from commercial use.

## Information for Data Requests and Correspondence

Chirag R. Parikh

Michael T. Eadon

Sanjay Jain

## Data Availability

Supplemental tables 1-9 detail the source and location of each sample used in the study, below are general landing pages that direct to these datasets.

Analytical pipelines and code used are available at https://github.com/KPMP-Scientific/KPMP-Atlas-v2.

*Processed and raw (except sequencing files) data, interactive and visualization tools: The snRNA-seq and scRNAseq visualization of the entire data (HuBMAP, KPMP, KTRC) can be accessed publicly on CxG website at:* https://cellxgene.cziscience.com/collections/9c9d04c4-8899-417f-bb6f-6107dcadf14f.

The snRNA-seq and scRNAseq visualization of the entire data (HuBMAP, KPMP, KTRC) can be accessed in the KPMP Atlas Explorer by October 31, 2025. The KPMP biopsy generated scRNA-seq, snRNA-seq, Multiome, Visium and spatial metabolomics data can be accessed from KPMP Atlas Repository by October 31, 2025 (DOI: 10.48698/16dd-vj20) except that raw sequencing files can be requested from by signing a data use agreement with KPMP.

The HuBMAP or KTRC reference or disease tissue data can be access through the HuBMAP publication page upon acceptance of the peer-reviewed manuscript. This includes WashU and IU Xenium, CosMx, SlideSeq2, Visium fresh frozen and Multiome data sets. Visium FFPE samples will be available from GEO (GSE307817). Visium data and histological images coregistration can be viewed using the FUSION tool (https://athena.rc.ufl.edu/#folder/67e31a8b8733e17e29775c98; instructions at <u>10.5281/zenodo.17296697</u>; user-public, password – public for navigating in FUSION) (see Methods and Border et al., 2025). The HuBMAP raw sequencing data will be available for download from the database of Genotypes and Phenotypes (dbGaP, phs002249) upon acceptance of peer-reviewed manuscript.

The raw KTRC sequencing files can be requested by contacting and signing a data use agreement to protect patient confidentiality.

Mouse ageing and injury models Multiome data are available from GEO (GSE308709).

*Additional published/public data sets:* The following publicly available RNA-seq data sets were used in this study: Human fibroblast scRNA-seq data from Buechler et al., 2021 was obtained from https://fibroXplorer.com; human fibroblast scRNA-seq from Korsunsky et al. 2022 was obtained from https://doi.org/10.5072/zenodo.772596; human myeloid scRNA-seq from Eraslan et al., 2022 was obtained from the GTEx Portal (www.gtexportal.org); mouse myeloid scRNA-seq from Conway et al., 2020 was obtained from the authors and is available from GEO (GSE140023); mouse IRI time-series snRNA-seq from Kirita et al., 2020 was obtained from GEO (GSE139107); mouse IRI time-series 10X multiome data from Gerhardt et al., 2023 was obtained from the authors and is available from GEO (GSE209610); mouse snRNA-seq from diabetic/hypertensive mice from Wu et al., 2022 was obtained from GEO (GSE184652).

Human SGLT2i data from Schaub et al. 2023 was obtained from GEO (GSE220939). Clinical data from ASSESS-AKI study can be accessed at https://repository.niddk.nih.gov/study/74. The TRIBE-AKI study, NAIKID Study, and Hopkins Health Reference Study data are under controlled access, contact Dr. Chirag Parikh at.

*Figures*: Source data are provided with this paper as a source data file. Schemata of the human nephron and renal corpuscle were developed by KPMP/HuBMAP (https://doi.org/10.48698/DEM4-0Q93). Additional schemata were generated using BioRender.)

## Code Availability

Code to reproduce figures is available from https://github.com/KPMP-Scientific/KPMP-Atlas-v2 upon acceptance of the peer-reviewed manuscript. No additional custom computational code was generated in this study.

## Kidney Precision Medicine Project Collaborative author list

Stewart H Lecker^80^, Alexander Morales^80^, Mark E Williams^80^, Steve Bogen^81^, Dongwon Lee^82^, Stephanie J Aw^83^, Laurence H Beck^83^, Marie F Calixte^83^, Kifle Gebre^83^, Molly C Geraghty^83^, Courtney Huynh^83^, Astrid Larson^83^, Minxin Lu^83^, Shana Maikhor^83^, Keyvona Moultrie^83^, R Narasimhan^83^, Ingrid F Onul^83^, Florencia A Rojas-Miguez^83^, Sophia H Rosan^83^, Ashish Upadhyay^83^, Ashish Verma^83^, Pranav Yadati^83^, BA Guanghao Yu^83^, Yan Zhou^83^, Mia R Colona^84^, Gearoid Michael McMahon^84^, Helmut Rennke^84^, Michael Todd Valerius^84^, Astrid Weins^84^, Anna Greka^85^, Nir Hacohen^85^, Jamie L Marshall^85^, Mark P Aulisio^86^, William S Bush^86^, Dana C Crawford^86^, Lakeshia Bush^87^, Leslie Cooperman^87^, Crystal A Gadegbeku^87^, Vivian Jeffers^87^, Stacey Jolly^87^, Kiasha Jones^87^, Michael Kuperman^87^, Marina Markovic^87^, Charles O’Malley^87^, Ellen Palmer^87^, Emilio D Poggio^87^, Teresa Randle^87^, Dianna Sendrey^87^, Kassandra Spates-Harden^87^, Jonathan J Taliercio^87^, Paul S Appelbaum^88^, Olivia Balderes^88^, Jonathan Barasch^88^, Andrew S Bomback^88^, Pietro A Canetta^88^, Vivette D’Agati^88^, Karla Mehl^88^, German varela^88^, Joana P Gonçalves^89^, Roy Lardenoije^89^, Lukasz G Migas^89^, Raf Van de Plas^89^, Mahla Asghari^2^, Daria Barwinska^2^, William S Bowen^2^, Kenneth W Dunn^2^, Michael Ferkowicz^2^, Danielle Janosevic^2^, Katherine J Kelley^2^, Azuma Nanamatsu^2^, Mohammad A Sohail^2^, Timothy A Sutton^2^, Kristine Conlon^6^, Reetika Ghag^6^, Amy McMurray^6^, Anitha Vijayan^6^, Akhil Ambekar^90^, Thomas M Coffman^90^, Xiang Li^90^, Bangchen Wang^90^, Andrew Janowczyk^91^, Anant Madabhushi^91^, Lun Ai^92^, Theodore Alexandrov^92^, Taneisha Campbell^93^, Jia-Yun Chen^94^, Nils Gehlenborg^94^, Mark S Keller^94^, Jia-Ren Lin^94^, Seymour Rosen^94^, Sandro Santagata^94^, Yi Zhang^94^, Charlotte Boys^95^, Leonie Küchenhoff^95^, Jini Ashok Bhanushali^96^, Sharon B Bledsoe^96^, Katy Börner^96^, Andreas Bueckle^96^, Bruce W Herr II^96^, Ellen M Quardokus^96^, Elizabeth G Record^96^, Marcelino Rivera^96^, Jennifer Stashevsky^96^, Abraham Verdoes^96^, Curtis Warfield^96^, Stephanie Wofford^96^, Devin M Wright^96^, Brittany C Minor^97^, Mohamed G Atta^98^, Mitali Barik^98^, Maria Chilo Bejarano^98^, Lauren Bernard^98^, Celia P Corona-Villalobos^98^, Derek M Fine^98^, Jeanine Hernandez^98^, Badra Kalil^98^, Jose M Monroy-Trujillo^98^, Sonya Shah^98^, C John Sperati^98^, Ashley R Wang^98^, Yumeng Wen^98^, Alan Xu^98^, Sophia Xu^98^, Sophia A Angus^99^, Sarah W Chen^99^, Isabel Donohoe^99^, Asari Henshaw^99^, Camille Johansen^99^, Mallory Mandel^99^, Jenny Molina-Guzman^99^, Neil Roy^99^, Melissa D Rubinsky^99^, Imane H Samari^99^, Paolo S Silva^99^, Anna Kate Stawicki^99^, Jennifer K Sun^99^, Julia A Welch^99^, Gabriel Zeinoun^99^, Evren U Azeloglu^100^, Kirk N Campbell^100^, Lili Chan^100^, Marina deCos^100^, Ashveena L Dighe^100^, Lorraine Evo-Ortega^100^, Lili Gai^100^, Ronald E Gordon^100^, Mark L Green^100^, Ritu Gupta^100^, Jonathan Haydak^100^, John Cijiang He^100^, Carol R Horowitz^100^, Gina Koch^100^, Patricia Kovatch^100^, Brandon G Larson^100^, Sean Lefferts^100^, Kristin Meliambro^100^, Girish N Nadkarni^100^, Marissa Patel^100^, Timothy D Quinn^100^, Tejas Rao^100^, Rosamond Rhodes^100^, Glenda V Roberts^100^, Daniel Stalbow^100^, Isaac E Stillman^100^, Joji Tokita^100^, Rachel Ustoyev^100^, Stephen C Ward^100^, Samuel Mon-Wei Yu^100^, Gek Cher Chan^101^, Pottumarthi V Prasad^102^, Samir V Parikh^103^, Brad H Rovin^103^, Jessica Lukowski^104^, Ljiljana Paša-Tolić^104^, Heidi L Vandyk^104^, George (Holt) Oliver^105^, Weiguang Mao^106^, Ksenia Sokolova^106^, Aaron Wong^106^, Ari Pollack^107^, Brandon Ginley^108^, Brendon Lutnick^108^, Thajudeen b^109^, David H Beyda^109^, Erika R Bracamonte^109^, Baltazar Campos^109^, Austin Derma^109^, Daniel Damian Duran^109^, Griselda Gamez^109^, Nicole Marquez^109^, Katherine Mendoza^109^, Ana Celina Sanora^109^, Raymond Scott^109^, Gregory Woodhead^109^, Kavya Anjani^110^, James G Cimino^110^, Zoltan G Laszik^110^, Tariq Mukatash^110^, Dane Munar^110^, Tara K Sigdel^110^, Leah Guthrie^111^, Milda R Saunders^112^, Ashley R Burg^113^, Hsieh EWY^114^, Joshua M Thurman^114^, Carissa Vinovskis^114^, Julia Wrobel^114^, Samuel Border^115^, Manoj Kumar Galla^115^, Harshit Lohaan^115^, Sayat Mimar^115^, Ahmed Naglah^115^, Anindya S Paul^115^, Joed Ancheta^116^, James T Bui^116^, Eunice Carmona-Powell^116^, Monica L Fox^116^, Ron C Gaba^116^, Tanika N Kelly^116^, Natalie Meza^116^, Arabela Quiroga^116^, Devona Redmond^116^, Amada Renteria^116^, Aaron Scroggins^116^, Kim Silva^116^, Anand Srivastava^116^, Michael Tanious^116^, Francesca Annese^117^, Heather K Ascani^117^, Victoria M Blanc^117^, Ninive Conser^117^, Nathan Creger^117^, Rachel Dull^117^, Sean Eddy^117^, Renee Frey^117^, Josh Hartley^117^, John Hartman^117^, Wenjun Ju^117^, Chrysta C Lienczewski^117^, Lili Liu^117^, Laura H Mariani^117^, Phillip J McCown^117^, Abhijit S Naik^117^, Rebecca Reamy^117^, Michael P Rose^117^, Cathy Smith^117^, Becky Steck^117^, Lalita Subramanian^117^, Haneen Tout^117^, Zach Wright^117^, Oyedele A Adeyi^118^, Alison Bunio Alvear^118^, Cathy A Bagne^118^, Jerica M Berge^118^, Alyson Coleman^118^, Yanli Ding^118^, PE Drawz^118^, Donna D’Souza^118^, Siobhan M Flanagan^118^, Ann Gentry^118^, Tasma Harindhanavudhi^118^, Dori Henderson^118^, Christopher J Jones^118^, Rachel R Kaspari^118^, Susan Klett^118^, Sisi Ma^118^, Patrick H Nachman^118^, Oluwatosin Oluwole^118^, Salma Rabi^118^, Via Rao^118^, Nicolas J Rauwolf ^118^, Elizabeth A Rogers^118^, Michael S Rosenberg^118^, Sami Safadi^118^, Sandeep Sharma^118^, Michelle L Snyder^118^, Susan M Wolf ^118^, Zoe Wright^118^, Seth Winfree^119^, Tashas Cameron-Wheeler^120^, Mary M Collie^120^, Anne Froment^120^, Samuel Haddad^120^, J Charles Jennette^120^, Jennifer L Jones^120^, Dhatri Kakarla^120^, Nicole Keefe^120^, Sara S Kelley^120^, Sora Lee^120^, Priya Mody^120^, Vanessa Moreno^120^, Amy K Mottl^120^, Sandhya Sundar Rajan^120^, Saad Mohammed Shariff^120^, Fernanda Ochoa Toro^120^, Evan M Zeitler^120^, Adam Burgess^121^, Michele M Elder^121^, Matthew Gilliam^121^, Daniel E Hall^121^, John A Kellum^121^, Raghavan Murugan^121^, Matthew R Rosengart^121^, Roderick Tan^121^, Tina Vita^121^, James Winters^121^, Bhupendra Kumar Gurung^122^, Annapurna Pamreddy^122^, Nagarjunachary Ragi^122^, Manjeri venkatachalam^122^, Hongping Ye^122^, Guanshi Zhang^122^, Shiqi Zhang^122^, Qi Cai^123^, Catherine Campbell^123^, S Susan Hedayati^123^, Allen R Hendricks^123^, Sanjeeva P Kalva^123^, Asra Kermani^123^, Simon C Lee^123^, Shihong Ma^123^, Meredith C McAdams^123^, Choudhary Moaz^123^, Harold Park^123^, Jiten Patel^123^, Boris S Patlis^123^, Anil Pillai^123^, Jose R Torrealba^123^, Miguel A Vazquez^123^, Nancy Wang^123^, Natasha Wen^123^, Mona Babaie^124^, Ashley C Berglund^124^, Brooke Berry^124^, Kristina N Blank^124^, Keith D Brown^124^, Jonas M Carson^124^, Matthew Dekker^124^, Frederick Dowd^124^, Stephanie M Grewenow^124^, Lynda Hayashi^124^, Andrew N Hoofnagle^124^, Nichole M Jefferson^124^, Cienn N Joyeux^124^, Richard A Knight^124^, Christine P Limonte^124^, Robyn L McClelland^124^, Yunbi Nam^124^, Christopher Park^124^, Jimmy Phuong^124^, Alexa Plisiewicz^124^, Laura Pyle^124^, Kasra A Rezaei^124^, Natalya Sarkisova^124^, Kelly D Smith^124^, Jaime Snyder^124^, Christy Stutzke^124^, Katherine R Tuttle^124^, Ruikang Wang^124^, Artit Wangperawong^124^, Adam Wilcox^124^, Kayleen Williams^124^, Bessie A Young^124^, Jamie L Allen^125^, Madeline E Colley^125^, Yarieli Cuevas-Rios^125^, Mark P de Caestecker^125^, Ruining Deng^125^, Martin Dufresne^125^, Yuankai Huo^125^, Angela R.S Kruse^125^, Tanima Arora^126^, Liam Brown^126^, Tif anny Budiman^126^, Lloyd G Cantley^126^, Vijayakumar R Kakade^126^, Candice A Kent^126^, Petra M Leite^126^, Dennis G Moledina^126^, Melissa M Shaw^126^, Jeffrey M Turner^126^, Ugochukwu Ugwuowo^126^, Angela M Victoria-Castro^126^, Joseph Ardayfio^127^, Jack Bebiak^127^, Roy Pinkeney^127^, John Saul^127^

## Kidney Precision Medicine Project Affiliations

^80^Beth Israel Deaconess Medical Center, Boston, MA 02215, ^81^Boston Cell Standards, Boston, MA 02111, ^82^Boston Children’s Hospital, Boston, MA 02115, ^83^Boston Medical Center, Boston, MA 02118, ^84^Brigham and Women’s Hospital, Boston, MA 02115, ^85^Broad Institute of MIT and Harvard, Cambridge, MA 02142, ^86^Case Western Reserve University, Cleveland, OH 44106, ^87^Cleveland Clinic, Cleveland, OH 44195, ^88^Columbia University, New York, NY 10027, ^89^Delft University of Technology, Delft, Netherlands, ^90^Duke University, Durham, NC 27708, ^91^Emory University, Atlanta, GA 30322, ^92^European Molecular Biology Laboratory, Meyerhofstraße 1, 69117 Heidelberg Germany, ^93^Gift of Life Michigan, Ann Arbor, MI 48108, ^94^Harvard University, Cambridge, MA 02138, ^95^Heidelberg University, 69117 Heidelberg, Germany, ^96^Indiana University, Bloomington, IN 47405, ^97^Institute of Informtics, Department of Medicine, Washington University School of Medicine, St. Louis, MO 63110, USA, ^98^Johns Hopkins University, Baltimore, MD 21218, ^99^Joslin Diabetes Center, Boston, MA 02215, ^100^Mount Sinai, New York, NY 10029, ^101^National University of Singapore, Singapore, Singapore 119077, ^102^Northwestern University, Evanston, IL 60208, ^103^Ohio State University, Columbus, OH 43210, ^104^Pacific Northwest National Laboratory, Richland, WA 99354, ^105^Parkland Health and Hospital System, Dallas, TX 75235, ^106^Princeton University, Princeton, NJ 08544, ^107^Seattle Children’s Hospital, Seattle, WA 98105, ^108^SUNY Buffalo, Buffalo, NY 14260, ^109^University of Arizona, Tucson, AZ 85719, ^110^University of California San Francisco, San Francisco, CA 94143, ^111^University of California, Berkeley, Berkeley, CA 94074, ^112^University of Chicago Medicine, Chicago, IL 60637, ^113^University of Cincinnati, Cincinnati, OH 45221, ^114^University of Colorado, Boulder, CO 80309, ^115^University of Florida, Gainesville, FL 32611, ^116^University of Illinois Chicago, Chicago, IL 60607, ^117^University of Michigan, Ann Arbor, MI 48109, ^118^University of Minnesota, Minneapolis, MN 55455, ^119^University of Nebraska Medical Center, Omaha, NE 68198, ^120^University of North Carolina, Chapel Hill, NC 27599, ^121^University of Pittsburgh, Pittsburgh, PA 15260, ^122^University of Texas Health Science Center at San Antonio, San Antonio, TX 78229, ^123^University of Texas Southwestern, Dallas, TX 75390, ^124^University of Washington, Seattle, WA 98195, ^125^Vanderbilt University, Nashville, TN 37235, ^126^Yale University, New Caven, CT 06520, ^127^Unaffiliated

## Notes

https://www.kpmp.org/doi-collection/10-48698-16dd-vj20

https://cellxgene.cziscience.com/collections/9c9d04c4-8899-417f-bb6f-6107dcadf14f

https://portal.hubmapconsortium.org/browse/publication/348186d1bda6d6dc764f746bbe94785e

### Summary of Updates

The initial Suppl. Table 20 that showed subtype-selective pathways was replaced by the correct Suppl. Table 20 that shows condition-selective pathways. We updated the Source Data sheet Fif2a,f; Fig3a; EDFig4 since it contained hash tag symbols instead of numeric values in some of its columns. We replaced the pie chart for moMAC-C3+ by the pie chart for moMAC-HBEGF+ in Figure 2g. We update Data Availability statement to include available links to visualization tools and data

## REFERENCES

1 Lake, B. B. et al. An atlas of healthy and injured cell states and niches in the human kidney. Nature 619, 585–594 (2023). 10.1038/s41586-023-05769-3

2 de Boer, I. H. et al. Rationale and design of the Kidney Precision Medicine Project. Kidney Int 99, 498–510 (2021). 10.1016/j.kint.2020.08.039

3 Jain, S. et al. Advances and prospects for the Human BioMolecular Atlas Program (HuBMAP). Nature Cell Biology 25, 1089–1100 (2023). 10.1038/s41556-023-01194-w

4 Abedini, A. et al. Single-cell multi-omic and spatial profiling of human kidneys implicates the fibrotic microenvironment in kidney disease progression. Nat Genet 56, 1712–1724 (2024). 10.1038/s41588-024-01802-x

5 Muto, Y. et al. Single cell transcriptional and chromatin accessibility profiling redefine cellular heterogeneity in the adult human kidney. Nat Commun 12, 2190 (2021). 10.1038/s41467-021-22368-w

6 Kramann, R. et al. Perivascular Gli1+ progenitors are key contributors to injury-induced organ fibrosis. Cell Stem Cell 16, 51–66 (2015). 10.1016/j.stem.2014.11.004

7 Hansen, J. et al. A reference tissue atlas for the human kidney. Sci Adv 8, eabn4965 (2022). 10.1126/sciadv.abn4965

8 Buechler, M. B. et al. Cross-tissue organization of the fibroblast lineage. Nature 593, 575–579 (2021). 10.1038/s41586-021-03549-5

9 Conway, B. R. et al. Kidney Single-Cell Atlas Reveals Myeloid Heterogeneity in Progression and Regression of Kidney Disease. Journal of the American Society of Nephrology 31, 2833–2854 (2020). 10.1681/asn.2020060806

10 Domínguez Conde, C., et al. Cross-tissue immune cell analysis reveals tissue-specific features in humans. Science 376, eabl5197 (2022). doi:10.1126/science.abl5197

11 Eraslan, G. et al. Single-nucleus cross-tissue molecular reference maps toward understanding disease gene function. Science 376, eabl4290 (2022). doi:10.1126/science.abl4290

12 Gao, Y. et al. Cross-tissue human fibroblast atlas reveals myofibroblast subtypes with distinct roles in immune modulation. Cancer Cell 42, 1764–1783.e1710 (2024). 10.1016/j.ccell.2024.08.020

13 Gerhardt, L. M. S., Liu, J., Koppitch, K., Cippà, P. E. & McMahon, A. P. Single-nuclear transcriptomics reveals diversity of proximal tubule cell states in a dynamic response to acute kidney injury. Proceedings of the National Academy of Sciences 118, e2026684118 (2021). doi:10.1073/pnas.2026684118

14 Kirita, Y., Wu, H., Uchimura, K., Wilson, P. C. & Humphreys, B. D. Cell profiling of mouse acute kidney injury reveals conserved cellular responses to injury. Proceedings of the National Academy of Sciences 117, 15874–15883 (2020). doi:10.1073/pnas.2005477117

15 Korsunsky, I. et al. Cross-tissue, single-cell stromal atlas identifies shared pathological fibroblast phenotypes in four chronic inflammatory diseases. Med 3, 481–518.e414 (2022). 10.1016/j.medj.2022.05.002

16 Ransick, A. et al. Single-Cell Profiling Reveals Sex, Lineage, and Regional Diversity in the Mouse Kidney. Developmental Cell 51, 399–413.e397 (2019). 10.1016/j.devcel.2019.10.005

17 Rodríguez-Morales, P. & Franklin, R. A. Macrophage phenotypes and functions: resolving inflammation and restoring homeostasis. Trends in Immunology 44, 986–998 (2023). 10.1016/j.it.2023.10.004

18 Gerhardt, L. M. S. et al. Lineage Tracing and Single-Nucleus Multiomics Reveal Novel Features of Adaptive and Maladaptive Repair after Acute Kidney Injury. J Am Soc Nephrol 34, 554–571 (2023). 10.1681/asn.0000000000000057

19 Hansen, J., Meretzky, D., Woldesenbet, S., Stolovitzky, G. & Iyengar, R. A flexible ontology for inference of emergent whole cell function from relationships between subcellular processes. Sci Rep 7, 17689 (2017). 10.1038/s41598-017-16627-4

20 Noorani, B. et al. Venetoclax pharmacokinetics in subjects with end-stage renal disease undergoing hemodialysis. Journal of Clinical Oncology 41, e19042–e19042 (2023). 10.1200/JCO.2023.41.16_suppl.e19042

21 Rieder, F. et al. Fibrosis: cross-organ biology and pathways to development of innovative drugs. Nature Reviews Drug Discovery 24, 543–569 (2025). 10.1038/s41573-025-01158-9

22 Buechler, M. B., Fu, W. & Turley, S. J. Fibroblast-macrophage reciprocal interactions in health, fibrosis, and cancer. Immunity 54, 903–915 (2021). 10.1016/j.immuni.2021.04.021

23 Li, Z., Nagai, J. S., Kuppe, C., Kramann, R. & Costa, I. G. scMEGA: single-cell multi-omic enhancer-based gene regulatory network inference. Bioinformatics Advances 3 (2023). 10.1093/bioadv/vbad003

24 Chen, X. et al. Mapping disease regulatory circuits at cell-type resolution from single-cell multiomics data. Nature Computational Science 3, 644–657 (2023). 10.1038/s43588-023-00476-5

25 Karsdal, M. A. et al. The good and the bad collagens of fibrosis - Their role in signaling and organ function. Adv Drug Deliv Rev 121, 43–56 (2017). 10.1016/j.addr.2017.07.014

26 Hazell, G. G. et al. PI16 is a shear stress and inflammation-regulated inhibitor of MMP2. Sci Rep 6, 39553 (2016). 10.1038/srep39553

27 Singhmar, P. et al. The fibroblast-derived protein PI16 controls neuropathic pain. Proc Natl Acad Sci U S A 117, 5463–5471 (2020). 10.1073/pnas.1913444117

28 Reyes, N. S. et al. Sentinel *p16*^INK4a+^ cells in the basement membrane form a reparative niche in the lung. Science 378, 192–201 (2022). doi:10.1126/science.abf3326

29 Qi, X. Y. et al. Fibroblast inward-rectifier potassium current upregulation in profibrillatory atrial remodeling. Circ Res 116, 836–845 (2015). 10.1161/circresaha.116.305326

30 Xing, C., Bao, L., Li, W. & Fan, H. Progress on role of ion channels of cardiac fibroblasts in fibrosis. Front Physiol 14, 1138306 (2023). 10.3389/fphys.2023.1138306

31 Southard, K. M. et al. Comprehensive transcription factor perturbations recapitulate fibroblast transcriptional states. Nature Genetics (2025). 10.1038/s41588-025-02284-1

32 Chakarov, S. et al. Two distinct interstitial macrophage populations coexist across tissues in specific subtissular niches. Science 363, eaau0964 (2019). doi:10.1126/science.aau0964

33 Burster, T. Processing and regulation mechanisms within antigen presenting cells: a possibility for therapeutic modulation. Curr Pharm Des 19, 1029–1042 (2013). 10.2174/1381612811319060005

34 Liu, Y., An, Y., Li, G. & Wang, S. Regulatory mechanism of macrophage polarization based on Hippo pathway. Frontiers in Immunology Volume 14-2023 (2023). 10.3389/fimmu.2023.1279591

35 Ruffell, D. et al. A CREB-C/EBPβ cascade induces M2 macrophage-specific gene expression and promotes muscle injury repair. Proceedings of the National Academy of Sciences 106, 17475–17480 (2009). doi:10.1073/pnas.0908641106

36 Shang, Y. et al. The transcriptional repressor Hes1 attenuates inflammation by regulating transcription elongation. Nat Immunol 17, 930–937 (2016). 10.1038/ni.3486

37 Chung, E. Y. et al. Interleukin-10 expression in macrophages during phagocytosis of apoptotic cells is mediated by homeodomain proteins Pbx1 and Prep-1. Immunity 27, 952–964 (2007). 10.1016/j.immuni.2007.11.014

38 Zhang, F. et al. IFN-γ and TNF-α drive a CXCL10+ CCL2+ macrophage phenotype expanded in severe COVID-19 lungs and inflammatory diseases with tissue inflammation. Genome Medicine 13, 64 (2021). 10.1186/s13073-021-00881-3

39 Cui, A. et al. Dictionary of immune responses to cytokines at single-cell resolution. Nature 625, 377–384 (2024). 10.1038/s41586-023-06816-9

40 Kuo, D. et al. HBEGF(+) macrophages in rheumatoid arthritis induce fibroblast invasiveness. Sci Transl Med 11 (2019). 10.1126/scitranslmed.aau8587

41 Zhu, B. et al. Uncoupling of macrophage inflammation from self-renewal modulates host recovery from respiratory viral infection. Immunity 54, 1200–1218.e1209 (2021). 10.1016/j.immuni.2021.04.001

42 Ouyang, J. F. et al. Systems level identification of a matrisome-associated macrophage polarisation state in multi-organ fibrosis. eLife 12, e85530 (2023). 10.7554/eLife.85530

43 Fang, L. et al. Transcriptional factor EB regulates macrophage polarization in the tumor microenvironment. OncoImmunology 6, e1312042 (2017). 10.1080/2162402X.2017.1312042

44 Liu, B. C., Tang, T. T., Lv, L. L. & Lan, H. Y. Renal tubule injury: a driving force toward chronic kidney disease. Kidney Int 93, 568–579 (2018). 10.1016/j.kint.2017.09.033

45 Adams, T. S. et al. Single-cell RNA-seq reveals ectopic and aberrant lung-resident cell populations in idiopathic pulmonary fibrosis. Science Advances 6, eaba1983 (2020). doi:10.1126/sciadv.aba1983

46 Franzén, L. et al. Mapping spatially resolved transcriptomes in human and mouse pulmonary fibrosis. Nature Genetics 56, 1725–1736 (2024). 10.1038/s41588-024-01819-2

47 Jiang, M. et al. Mitochondrial dysfunction and the AKI-to-CKD transition. Am J Physiol Renal Physiol 319, F1105–f1116 (2020). 10.1152/ajprenal.00285.2020

48 Balzer, M. S. et al. Single-cell analysis highlights differences in druggable pathways underlying adaptive or fibrotic kidney regeneration. Nat Commun 13, 4018 (2022). 10.1038/s41467-022-31772-9

49 Hato, T. et al. Bacterial sepsis triggers an antiviral response that causes translation shutdown. J Clin Invest 129, 296–309 (2019). 10.1172/jci123284

50 Wynn, T. A. & Ramalingam, T. R. Mechanisms of fibrosis: therapeutic translation for fibrotic disease. Nat Med 18, 1028–1040 (2012). 10.1038/nm.2807

51 Hills, C.-E., Siamantouras, E. & Squires, P.-E. Cell adhesion in renal tubular epithelial cells: Biochemistry, biophysics or both. BIOCELL 46, 937--940 (2022).

52 Goodwin, K. et al. Cell-cell and cell-extracellular matrix adhesions cooperate to organize actomyosin networks and maintain force transmission during dorsal closure. Mol Biol Cell 28, 1301–1310 (2017). 10.1091/mbc.E17-01-0033

53 Schmidt, I. M. et al. Plasma proteomics of acute tubular injury. Nature Communications 15, 7368 (2024). 10.1038/s41467-024-51304-x

54 Gisch, D. L. et al. The chromatin landscape of healthy and injured cell types in the human kidney. Nature Communications 15, 433 (2024). 10.1038/s41467-023-44467-6

55 Ledru, N. et al. Predicting proximal tubule failed repair drivers through regularized regression analysis of single cell multiomic sequencing. Nature Communications 15, 1291 (2024). 10.1038/s41467-024-45706-0

56 Wang, C.-H. et al. TMPRSS4 facilitates epithelial-mesenchymal transition of hepatocellular carcinoma and is a predictive marker for poor prognosis of patients after curative resection. Scientific Reports 5, 12366 (2015). 10.1038/srep12366

57 Gokey, J. J. et al. MEG3 is increased in idiopathic pulmonary fibrosis and regulates epithelial cell differentiation. JCI Insight 3 (2018). 10.1172/jci.insight.122490

58 Aggarwal, S. et al. SOX9 switch links regeneration to fibrosis at the single-cell level in mammalian kidneys. Science 383, eadd6371 (2024). doi:10.1126/science.add6371

59 Stanzick, K. J. et al. Discovery and prioritization of variants and genes for kidney function in >1.2 million individuals. Nature Communications 12, 4350 (2021). 10.1038/s41467-021-24491-0

60 Verissimo, T. et al. PCK1 is a key regulator of metabolic and mitochondrial functions in renal tubular cells. Am J Physiol Renal Physiol 324, F532–f543 (2023). 10.1152/ajprenal.00038.2023

61 Wang, H. et al. Perturbed gut microbiome and fecal and serum metabolomes are associated with chronic kidney disease severity. Microbiome 11, 3 (2023). 10.1186/s40168-022-01443-4

62 Sharma, K. et al. Endogenous adenine mediates kidney injury in diabetic models and predicts diabetic kidney disease in patients. J Clin Invest 133 (2023). 10.1172/jci170341

63 Sharma, K. et al. Metabolomics reveals signature of mitochondrial dysfunction in diabetic kidney disease. J Am Soc Nephrol 24, 1901–1912 (2013). 10.1681/asn.2013020126

64 Ferreira, R. M. et al. A MEF2C transcription factor network regulates proliferation of glomerular endothelial cells in diabetic kidney disease. bioRxiv (2024). 10.1101/2024.09.27.615250

65 Schiessl, I. M. The Role of Tubule-Interstitial Crosstalk in Renal Injury and Recovery. Semin Nephrol 40, 216–231 (2020). 10.1016/j.semnephrol.2020.01.012

66 Yamashita, N. & Kramann, R. Mechanisms of kidney fibrosis and routes towards therapy. Trends Endocrinol Metab 35, 31–48 (2024). 10.1016/j.tem.2023.09.001

67 Border, S. et al. FUSION: A web-based application for in-depth exploration of multi-omics data with brightfield histology. bioRxiv (2024). 10.1101/2024.07.09.602778

68 Khang, A. et al. Automated prediction of fibroblast phenotypes using mathematical descriptors of cellular features. Nature Communications 16, 2841 (2025). 10.1038/s41467-025-58082-0

69 Ciminieri, C. et al. IL-1β Induces a Proinflammatory Fibroblast Microenvironment that Impairs Lung Progenitors’ Function. Am J Respir Cell Mol Biol 68, 444–455 (2023). 10.1165/rcmb.2022-0209OC

70 Varshney, P., Yadav, V. & Saini, N. Lipid rafts in immune signalling: current progress and future perspective. Immunology 149, 13–24 (2016). 10.1111/imm.12617

71 Lim, H.-K. et al. Phosphatidic Acid Regulates Systemic Inflammatory Responses by Modulating the Akt-Mammalian Target of Rapamycin-p70 S6 Kinase 1 Pathway*. Journal of Biological Chemistry 278, 45117–45127 (2003). 10.1074/jbc.M303789200

72 Privratsky, J. R. et al. Interleukin 1 receptor (IL-1R1) activation exacerbates toxin-induced acute kidney injury. Am J Physiol Renal Physiol 315, F682–f691 (2018). 10.1152/ajprenal.00104.2018

73 Narasimhan, H. et al. An aberrant immune–epithelial progenitor niche drives viral lung sequelae. Nature 634, 961–969 (2024). 10.1038/s41586-024-07926-8

74 Wang, F. et al. Regulation of epithelial transitional states in murine and human pulmonary fibrosis. The Journal of Clinical Investigation 133 (2023). 10.1172/JCI165612

75 Usman, K. et al. Interleukin-1α inhibits transforming growth factor-β1 and β2-induced extracellular matrix production, remodeling and signaling in human lung fibroblasts: Master regulator in lung mucosal repair. Matrix Biology 132, 47–58 (2024). 10.1016/j.matbio.2024.06.007

76 Eggert, T. et al. Distinct Functions of Senescence-Associated Immune Responses in Liver Tumor Surveillance and Tumor Progression. Cancer Cell 30, 533–547 (2016). 10.1016/j.ccell.2016.09.003

77 Liu, J. et al. Renoprotective and Immunomodulatory Effects of GDF15 following AKI Invoked by Ischemia-Reperfusion Injury. J Am Soc Nephrol 31, 701–715 (2020). 10.1681/asn.2019090876

78 Jones, K. et al. SOX4 and RELA Function as Transcriptional Partners to Regulate the Expression of TNF-Responsive Genes in Fibroblast-Like Synoviocytes. Front Immunol 13, 789349 (2022). 10.3389/fimmu.2022.789349

79 Lourenço, A. R. & Coffer, P. J. SOX4: Joining the Master Regulators of Epithelial-to-Mesenchymal Transition? Trends in Cancer 3, 571–582 (2017). 10.1016/j.trecan.2017.06.002

80 Ikushima, H. et al. Autocrine TGF-beta signaling maintains tumorigenicity of glioma-initiating cells through Sry-related HMG-box factors. Cell Stem Cell 5, 504–514 (2009). 10.1016/j.stem.2009.08.018

81 Parikh, C. R. et al. Postoperative biomarkers predict acute kidney injury and poor outcomes after adult cardiac surgery. J Am Soc Nephrol 22, 1748–1757 (2011). 10.1681/asn.2010121302

82 Mou, L. et al. Integrative analysis of COL6A3 in lupus nephritis: insights from single-cell transcriptomics and proteomics. Frontiers in Immunology Volume 15 - 2024 (2024). 10.3389/fimmu.2024.1309447

83 Rasmussen, D. G. K. et al. Urinary endotrophin predicts disease progression in patients with chronic kidney disease. Scientific Reports 7, 17328 (2017). 10.1038/s41598-017-17470-3

84 Williams, L., Layton, T., Yang, N., Feldmann, M. & Nanchahal, J. Collagen VI as a driver and disease biomarker in human fibrosis. The FEBS Journal 289, 3603–3629 (2022). 10.1111/febs.16039

85 Kobayashi, H. et al. Neuroblastoma suppressor of tumorigenicity 1 is a circulating protein associated with progression to end-stage kidney disease in diabetes. Science Translational Medicine 14, eabj2109 (2022). doi:10.1126/scitranslmed.abj2109

86 Maksimowski, N. A. et al. Follistatin-Like-1 (FSTL1) Is a Fibroblast-Derived Growth Factor That Contributes to Progression of Chronic Kidney Disease. Int J Mol Sci 22 (2021). 10.3390/ijms22179513

87 Péterfi, Z. et al. Peroxidasin is secreted and incorporated into the extracellular matrix of myofibroblasts and fibrotic kidney. Am J Pathol 175, 725–735 (2009). 10.2353/ajpath.2009.080693

88 Templeton, E. M. et al. Identifying Candidate Protein Markers of Acute Kidney Injury in Acute Decompensated Heart Failure. Int J Mol Sci 23 (2022). 10.3390/ijms23021009

89 Fu, S. et al. Effects of Selenium on Chronic Kidney Disease: A Mendelian Randomization Study. Nutrients 14 (2022). 10.3390/nu14214458

90 Wu, Y. et al. Selenoprotein Gene mRNA Expression Evaluation During Renal Ischemia-Reperfusion Injury in Rats and Ebselen Intervention Effects. Biol Trace Elem Res 201, 1792–1805 (2023). 10.1007/s12011-022-03275-7

91 Schaub, J. A. et al. SGLT2 inhibitors mitigate kidney tubular metabolic and mTORC1 perturbations in youth-onset type 2 diabetes. J Clin Invest 133 (2023). 10.1172/jci164486

92 Chen, Q., Gao, C., Wang, M., Fei, X. & Zhao, N. TRIM18-Regulated STAT3 Signaling Pathway via PTP1B Promotes Renal Epithelial–Mesenchymal Transition, Inflammation, and Fibrosis in Diabetic Kidney Disease. Frontiers in Physiology Volume 12 - 2021 (2021). 10.3389/fphys.2021.709506

93 Sitole, B. N. & Mavri-Damelin, D. Peroxidasin is regulated by the epithelial-mesenchymal transition master transcription factor Snai1. Gene 646, 195–202 (2018). 10.1016/j.gene.2018.01.011

94 Wu, H. et al. Mapping the single-cell transcriptomic response of murine diabetic kidney disease to therapies. Cell Metab 34, 1064–1078.e1066 (2022). 10.1016/j.cmet.2022.05.010

95 Argentieri, M. A. et al. Proteomic aging clock predicts mortality and risk of common age-related diseases in diverse populations. Nature Medicine 30, 2450–2460 (2024). 10.1038/s41591-024-03164-7

96 Dubin, R. F. et al. Proteomics of CKD progression in the chronic renal insufficiency cohort. Nature Communications 14, 6340 (2023). 10.1038/s41467-023-41642-7

97 Eadon, M. T. et al. Kidney Histopathology and Prediction of Kidney Failure: A Retrospective Cohort Study. Am J Kidney Dis 76, 350–360 (2020). 10.1053/j.ajkd.2019.12.014

98 Barisoni, L. et al. Reproducibility of the NEPTUNE descriptor-based scoring system on whole-slide images and histologic and ultrastructural digital images. Mod Pathol 29, 671–684 (2016). 10.1038/modpathol.2016.58

99 Tervaert, T. W. et al. Pathologic classification of diabetic nephropathy. J Am Soc Nephrol 21, 556–563 (2010). 10.1681/asn.2010010010

100 Kefaloyianni, E. et al. Proximal Tubule-Derived Amphiregulin Amplifies and Integrates Profibrotic EGF Receptor Signals in Kidney Fibrosis. J Am Soc Nephrol 30, 2370–2383 (2019). 10.1681/asn.2019030321

101 Gayoso, A., Shor, J., Carr, A. J., Sharma, R. & Pe’er, D. GitHub: DoubletDetection. Zenodo (2019).

102 Hao, Y. et al. Dictionary learning for integrative, multimodal and scalable single-cell analysis. Nature Biotechnology 42, 293–304 (2024). 10.1038/s41587-023-01767-y

103 Stuart, T., Srivastava, A., Madad, S., Lareau, C. A. & Satija, R. Single-cell chromatin state analysis with Signac. Nature Methods 18, 1333–1341 (2021). 10.1038/s41592-021-01282-5

104 Ogata, J. D. et al. excluderanges: exclusion sets for T2T-CHM13, GRCm39, and other genome assemblies. Bioinformatics 39 (2023). 10.1093/bioinformatics/btad198

105 Xu, C. et al. Automatic cell-type harmonization and integration across Human Cell Atlas datasets. Cell 186, 5876–5891.e5820 (2023). 10.1016/j.cell.2023.11.026

106 Dumas, S. J. et al. Phenotypic diversity and metabolic specialization of renal endothelial cells. Nature Reviews Nephrology 17, 441–464 (2021). 10.1038/s41581-021-00411-9

107 Zappia, L. & Oshlack, A. Clustering trees: a visualization for evaluating clusterings at multiple resolutions. GigaScience 7 (2018). 10.1093/gigascience/giy083

108 Saxena, V. et al. Kidney intercalated cells are phagocytic and acidify internalized uropathogenic Escherichia coli. Nature Communications 12, 2405 (2021). 10.1038/s41467-021-22672-5

109 Schubert, M. et al. Perturbation-response genes reveal signaling footprints in cancer gene expression. Nature Communications 9, 20 (2018). 10.1038/s41467-017-02391-6

110 Jiang, L. et al. Systematic reconstruction of molecular pathway signatures using scalable single-cell perturbation screens. bioRxiv, 2024.2001.2029.576933 (2024). 10.1101/2024.01.29.576933

111 Schep, A. N., Wu, B., Buenrostro, J. D. & Greenleaf, W. J. chromVAR: inferring transcription-factor-associated accessibility from single-cell epigenomic data. Nature Methods 14, 975–978 (2017). 10.1038/nmeth.4401

112 Granja, J. M. et al. ArchR is a scalable software package for integrative single-cell chromatin accessibility analysis. Nature Genetics 53, 403–411 (2021). 10.1038/s41588-021-00790-6

113 Chen, X. et al. Mapping disease regulatory circuits at cell-type resolution from single-cell multiomics data. Nat Comput Sci 3, 644–657 (2023). 10.1038/s43588-023-00476-5

114 Kamimoto, K. et al. Dissecting cell identity via network inference and in silico gene perturbation. Nature 614, 742–751 (2023). 10.1038/s41586-022-05688-9

115 Badia-i-Mompel, P. et al. decoupleR: ensemble of computational methods to infer biological activities from omics data. Bioinformatics Advances 2 (2022). 10.1093/bioadv/vbac016

116 Dimitrov, D. et al. LIANA+ provides an all-in-one framework for cell–cell communication inference. Nature Cell Biology 26, 1613–1622 (2024). 10.1038/s41556-024-01469-w

117 Melo Ferreira, R., et al. Integration of spatial and single-cell transcriptomics localizes epithelial cell-immune cross-talk in kidney injury. JCI Insight 6 (2021). 10.1172/jci.insight.147703

118 Satija, R., Farrell, J. A., Gennert, D., Schier, A. F. & Regev, A. Spatial reconstruction of single-cell gene expression data. Nat Biotechnol 33, 495–502 (2015). 10.1038/nbt.3192

119 He, S. et al. High-plex imaging of RNA and proteins at subcellular resolution in fixed tissue by spatial molecular imaging. Nat Biotechnol 40, 1794–1806 (2022). 10.1038/s41587-022-01483-z

120 Stringer, C., Wang, T., Michaelos, M. & Pachitariu, M. Cellpose: a generalist algorithm for cellular segmentation. Nature Methods 18, 100–106 (2021). 10.1038/s41592-020-01018-x

121 Melo Ferreira, R. et al. A <em>MEF2C</em> transcripton factor network regulates proliferation of glomerular endothelial cells in diabetic kidney disease. bioRxiv, 2024.2009.2027.615250 (2024). 10.1101/2024.09.27.615250

122 Singhal, V. et al. BANKSY unifies cell typing and tissue domain segmentation for scalable spatial omics data analysis. Nature Genetics 56, 431–441 (2024). 10.1038/s41588-024-01664-3

123 Pang, Z. et al. MetaboAnalyst 6.0: towards a unified platform for metabolomics data processing, analysis and interpretation. Nucleic Acids Res 52, W398–w406 (2024). 10.1093/nar/gkae253

124 Farrow, M. A. et al. A lipid atlas of the human kidney. Science Advances 11, eadu3730 (2025). doi:10.1126/sciadv.adu3730

125 Spraggins, J. M. et al. High-Performance Molecular Imaging with MALDI Trapped Ion-Mobility Time-of-Flight (timsTOF) Mass Spectrometry. Analytical Chemistry 91, 14552–14560 (2019). 10.1021/acs.analchem.9b03612

126 Chen, T. & Guestrin, C. in Proceedings of the 22nd ACM SIGKDD International Conference on Knowledge Discovery and Data Mining 785–794 (Association for Computing Machinery, San Francisco, California, USA, 2016).

127 Lundberg, S. M. et al. From local explanations to global understanding with explainable AI for trees. Nature Machine Intelligence 2, 56–67 (2020). 10.1038/s42256-019-0138-9

128 Tideman, L. E. M. et al. Automated biomarker candidate discovery in imaging mass spectrometry data through spatially localized Shapley additive explanations. Analytica Chimica Acta 1177, 338522 (2021). 10.1016/j.aca.2021.338522

129 Veličković, D., Sharma, K., Alexandrov, T., Hodgin, J. B. & Anderton, C. R. Controlled Humidity Levels for Fine Spatial Detail Information in Enzyme-Assisted N-Glycan MALDI MSI. Journal of the American Society for Mass Spectrometry 33, 1577–1580 (2022). 10.1021/jasms.2c00120

130 Veličković, D. et al. Rapid Automated Annotation and Analysis of N-Glycan Mass Spectrometry Imaging Data Sets Using NGlycDB in METASPACE. Analytical Chemistry 93, 13421–13425 (2021). 10.1021/acs.analchem.1c02347

131 Asghari, M. et al. Integration of spatial protein imaging and transcriptomics in the human kidney tracks the regenerative potential of proximal tubules. Science Advances 11, eadv8918 (2025). doi:10.1126/sciadv.adv8918

132 Liu, J. et al. Cell-specific translational profiling in acute kidney injury. J Clin Invest 124, 1242–1254 (2014). 10.1172/jci72126

133 Herbert, B. S. et al. A telomerase immortalized human proximal tubule cell line with a truncation mutation (Q4004X) in polycystin-1. PLoS One 8, e55191 (2013). 10.1371/journal.pone.0055191

134 Liu, R. X. et al. Comparison of proteomic methods in evaluating biomarker-AKI associations in cardiac surgery patients. Transl Res 238, 49–62 (2021). 10.1016/j.trsl.2021.07.005

135 Hsu, C. Y. et al. Post-Acute Kidney Injury Proteinuria and Subsequent Kidney Disease Progression: The Assessment, Serial Evaluation, and Subsequent Sequelae in Acute Kidney Injury (ASSESS-AKI) Study. JAMA Intern Med 180, 402–410 (2020). 10.1001/jamainternmed.2019.6390

136 Srivastava, A. et al. The Prognostic Value of Histopathologic Lesions in Native Kidney Biopsy Specimens: Results from the Boston Kidney Biopsy Cohort Study. J Am Soc Nephrol 29, 2213–2224 (2018). 10.1681/asn.2017121260

