## Supplementary material for "Cellular and Spatial Drivers of Unresolved Injury and Functional Decline in the Human Kidney": SD Figure

#### Affiliations

[1]Altos Labs San Diego Institute of Science, San Diego, CA, USA, [2]Department of Medicine, Indiana University School of Medicine, Indianapolis, IN 46202, USA, [3]Department of Pharmacological Sciences, Icahn School of Medicine at Mount Sinai, New York, NY 10029, USA, [4]Mount Sinai Institute for Systems Biomedicine, Icahn School of Medicine at Mount Sinai, New York, NY 10029, USA, [5]Department of Computational Medicine and Bioinformatics, University of Michigan, Ann Arbor, MI 48109, USA, [6]Division of Nephrology, Department of Medicine, Washington University School of Medicine, St. Louis, MO 63110, USA, [7]Division of Nephrology, Johns Hopkins School of Medicine, Baltimore, MD 21287, USA, [8]Institute for Computational Biomedicine, Heidelberg University and Heidelberg University Hospital, Heidelberg, Germany, [9]Center for Computational Biology, Flatiron Institute, Simons Foundation, New York, NY, USA, [10]Lewis-Sigler Institute of Integrative Genomics, Princeton University, Princeton, NJ, USA, [11]Center for Precision Medicine, The University of Texas at San Antonio, San Antonio, TX, USA, [12]Department of Cell and Developmental Biology and Mass Spectrometry Research Center, Vanderbilt University School of Medicine, Nashville, TN 37232, USA, [13]Pacific Northwest National Laboratory, Richland, WA, USA, [14]Department of Medicine-Nephrology & Intelligent Critical Care Center, University of Florida, Gainesville, FL, USA, [15]Department of Medicine, Indiana University School of Medicine, Indianapolis, IN, 46202, USA, [16]Department of Medicine, Boston University Chobanian & Avedisian School of Medicine, Boston, MA, USA, [17]Department of Internal Medicine, Division of Nephrology, University of Michigan, Ann Arbor, MI 48109, USA, [18]Center for Computational Medicine and Bioinformatics,University of Michigan, Ann Arbor, MI 48109, USA, [19]Department of Medicine, Washington University School of Medicine, St. Louis, MO 63110, USA, [20]Institute of Informatics, Department of Medicine, Washington University School of Medicine, St. Louis, MO 63110, USA, [21]Center for Precision Medicine, The University of Texas Health San Antonio, San Antonio, TX, USA, [22]Division of Nephrology, Department of Medicine, The University of Texas Health San Antonio, San Antonio, TX, USA, [23]Unit for Laboratory Animal Medicine, Department of Learning Health Sciences, University of Michigan, Ann Arbor, MI 48109, USA, [24]Department of Biochemistry and Molecular Biology, Saint Louis University School of Medicine, St. Louis, MO, 63103, USA, [25]St. Louis Veteran Affairs Medical Center, St. Louis, MO, 63106, USA, [26]Department of Medicine, New York University, New York, USA, [27]Department of Laboratory Medicine and Pathology, University of Washington, Seattle, Washington, USA, [28]Department of Pathology, University of Michigan, Ann Arbor, MI 48109, USA, [29]Department of Pathology, Division of AI and Computational Pathology, Duke University, Durham, NC, USA, [30]Department of Medicine, Division of Nephrology, Duke University, Durham, NC, USA, [31]Kidney Research Institute, Division of Nephrology, 325 Ninth Avenue, Box 359606, Seattle, WA 98104, USA, [32]Department of Pathology, Microbiology and Immunology, Vanderbilt University Medical Center, Nashville, TN 37232 USA, [33]Department of Pathology and Laboratory Medicine, Boston University Chobanian & Avedisian School of Medicine, Boston, MA, USA, [34]Department of Anatomic Pathology, Cleveland Clinic, Cleveland, OH, USA, [35]Department of Pathology, Yale University, New Haven, CT, USA, [36]Department of Pathology,Thomas E Starzl Transplant Insititute & University of Pittsburgh School of Medicine, Pittsburgh, PA 15213, USA, [37]Department of Pathology, Johns Hopkins University, Baltimore, MD, USA, [38]Department of Pathology, University of Illinois at Chicago, Chicago, IL, USA, [39]Department of Medicine, University of Arizona, Tucson, AZ, USA, [40]Department of Endocrinology, Diabetes and Metabolism, Cleveland Clinic Foundation, Cleveland, OH, USA, [41]Division of Nephrology, Icahn School of Medicine at Mount Sinai, New York, NY, USA, [42]Washington University in Saint Louis, St. Louis, MO, 63103, USA, [43]Department of Surgery, University of Nevada Reno School of Medicine, Reno, NV 89502, USA, [44]Department of Physiology and Cell Biology, University of Nevada Reno School of Medicine, Reno, NV 89502, USA, [45]Department of Medicine, Columbia University, New York, NY 10032, USA, [46]Department of Medicine, University of Illinois Chicago, Chicago, IL, USA, [47]Department of Internal Medicine, University of Texas Southwestern Medical Center, Dallas, TX, USA, [48]Medicine Service, VA North Texas Health Care System, Dallas, TX, USA, [49]Pak Center for Mineral Metabolism and Clinical Research, University of Texas Southwestern Medical Center, Dallas, TX, USA, [50]Lerner Research and Medical Specialties Institutes, Cleveland Clinic, Cleveland, OH 44195, USA, [51]Department of Medicine, University of Pittsburgh School of Medicine, Pittsburgh, PA 15213, USA, [52]Division of Nephrology, Department of Medicine, Massachusetts General Hospital, Boston, MA, USA, [53]Kidney and Hypertension Unit, Joslin Diabetes Center and Harvard Medical School, Boston, MA 02215, USA, [54]Department of Medicine, Division of Nephrology and Hypertension, University of North Carolina School of Medicine, Chapel Hill, NC, USA, [55]Division of Multi-Organ Transplantation, Department of Surgery, University of California, San Francisco, San Francisco, CA, USA, [56]Department of Internal Medicine, UT Southwestern Medical Center, Dallas, TX 75390, USA, [57]Division of Nephrology, Istanbul School of Medicine, Istanbul, Turkey, [58]Section of Nephrology, Boston University School of Medicine and Boston Medical Center, Boston, MA 02118, USA, [59]Department of Anatomy, Cell Biology & Physiology, Indiana University School of Medicine, Indianapolis, IN 46202, USA, [60]Clinical and Translational Research Accelerator, Department of Medicine, Yale University School of Medicine, New

Haven, CT 06510, USA, [61]Department of Surgery, University of Cincinnati, Cincinnati, OH, USA, [62]Broad Institute of Harvard and MIT, Cambridge, MA 02142, USA, [63]European Bioinformatics Institute, Wellcome Genome Campus, Hinxton, Cambridgeshire, CB10 1SD, UK, [64]Heidelberg University, Faculty of Medicine, and Heidelberg University Hospital, Institute for Computational Biomedicine (ICB), Heidelberg, [65]University of Washington Medicine Diabetes Institute, University of Washington, Seattle, Washington, USA, [66]Department of Medicine, Division of Nephrology, University of Colorado School of Medicine, Aurora, CO, USA, [67]Department of Computer Science, Princeton University, Princeton, NJ 08544, USA, [68]Lewis-Sigler Institute for Integrative Genomics, Princeton University, Princeton, NJ 08544, USA, [69]Princeton Precision Health, Princeton University, Princeton, NJ 08544, USA, [70]Center for Computational Biology, Flatiron Institute, Simons Foundation, New York, NY 10010, USA, [71]Department of Medicine – Section of Quantitative Health, University of Florida, Gainesville, FL, USA, [72]Department of Medicine, Regenerative Medicine Institute, Cedars Sinai Medical Center, Los Angeles, CA, 90048, USA, [73]Icahn School of Medicine at Mount Sinai, New York, NY 10029, USA, [74]Department of Pathology and Immunology, Washington University School of Medicine, St. Louis, MO 63110, USA, [75]Department of Pediatrics, Washington University School of Medicine, St. Louis, MO 63110, USA, [76]Kidney Translational Research Center, Washington University School of Medicine, St. Louis, MO 63110, USA

[77]Co-first authors: Blue B. Lake, Ricardo Melo Ferreira, Jens Hansen, Rajasree Menon
[78]Contributed equally: Jeannine Basta, Heather Thiessen Philbrook, Stephanie Reinert, Robin Fallegger, Asmita K. Lagwankar, Xi Chen, Soumya Maity, Katerina V. Djambazova, Brittney L. Gorman, Nicholas Lucarelli, Debora L. Gisch, Insa M. Schmidt, Viji Nair, Fadhl Alakwaa, Eirini Kefaloyianni
[79]Co-senior authors: Kun Zhang, Ravi Iyengar, Matthias Kretzler, Jeffrey B. Hodgin, Chirag R. Parikh, Michael T. Eadon, Sanjay Jain
[*] A list of authors and their affiliations appears at the end of the paper

#Correspondence
Chirag R. Parikh:
Michael T. Eadon:
Sanjay Jain:

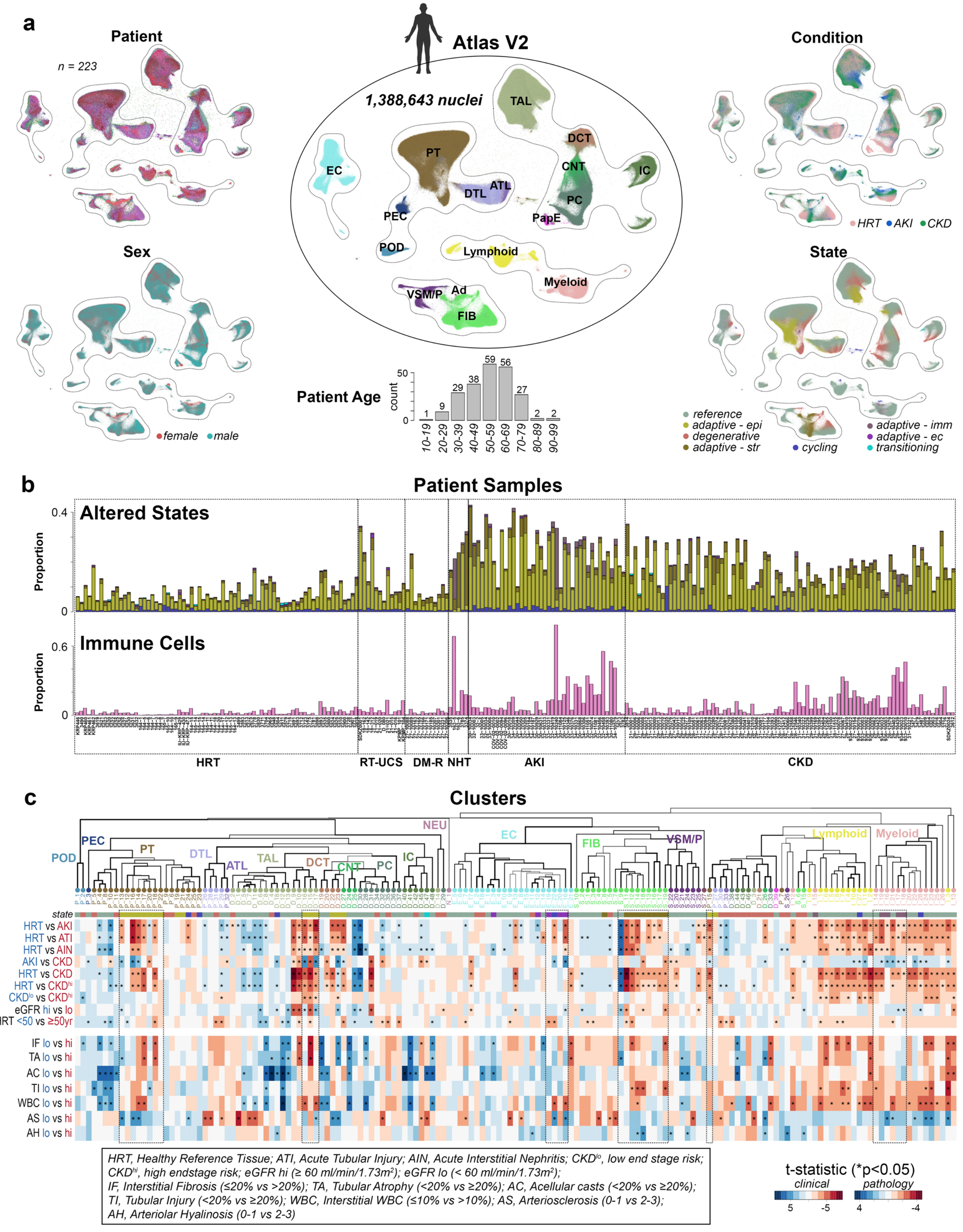
a
Patient
n = 223
Sex
female
male
Atlas V2
1,388,643 nuclei
TAL
DCT
CNT
IC
PC
PapE
PT
EC
ATL
DTL
PEC
POD
Lymphoid
Myeloid
VSM/P
Ad
FIB
Patient Age
count
50
0
1
9
29
38
59
56
27
2
2
10-19
20-29
30-39
40-49
50-59
60-69
70-79
80-89
90-99
Condition
HRT
AKI
CKD
State
reference
adaptive - epi
degenerative
adaptive - str
cycling
adaptive - imm
adaptive - ec
transitioning
b
Patient Samples
Altered States
Proportion
0.4
0
Immune Cells
0.6
HRT
RT-UCS
DM-R
NHT
AKI
CKD
c
Clusters
POD
PEC
PT
DTL
ATL
TAL
DCT
CNT
PC
IC
NEU
EC
FIB
VSM/P
Lymphoid
Myeloid
state
HRT vs AKI
HRT vs ATI
HRT vs AIN
AKI vs CKD
HRT vs CKD
HRT vs CKD$^{hi}$
CKD$^{lo}$ vs CKD$^{hi}$
eGFR hi vs lo
HRT <50 vs ≥50yr
IF lo vs hi
TA lo vs hi
AC lo vs hi
TI lo vs hi
WBC lo vs hi
AS lo vs hi
AH lo vs hi
HRT, Healthy Reference Tissue; ATI, Acute Tubular Injury; AIN, Acute Interstitial Nephritis; CKD$^{lo}$, low end stage risk; CKD$^{hi}$, high endstage risk; eGFR hi (≥ 60 ml/min/1.73m²); eGFR lo (< 60 ml/min/1.73m²);
IF, Interstitial Fibrosis (≤20% vs >20%); TA, Tubular Atrophy (<20% vs ≥20%); AC, Acellular casts (<20% vs ≥20%);
TI, Tubular Injury (<20% vs ≥20%); WBC, Interstitial WBC (≤10% vs >10%); AS, Arteriosclerosis (0-1 vs 2-3);
AH, Arteriolar Hyalinosis (0-1 vs 2-3)
t-statistic (*p<0.05)
clinical
pathology
5
-5
4
-4

**SD Figure 1. HKAv2 quality control. a**. UMAP embeddings showing broad subclasses (center plot) and the associated distribution of patients, sexes, disease conditions and altered states (outer plots). Barplot shows number of patients sampled for each age bin. **b**. Barplots showing the proportion of altered states (top) as indicated in (a) and the proportion of immune cells (bottom) across the different patients and disease conditions. **c**. Heatmaps showing cell type enrichments as t statistics for the comparison of cluster abundances between two different patient groupings. Asterisk indicates p values less than 0.05. Comparisons were done for clinical categories and pathology descriptor scoring. Only biopsy samples (excluding nephrectomy and deceased donor) were used for all categories except age associations. Number of patients for clinical: HRT (41), AKI (38), CKD (73), eGFRhi (69), eGFRlo (56), HRT <50 (34), HRT ≥50 (38), ATI (20), AIN (11), CKDhi (26), CKDlo (9). Number of patients for pathology: IF lo (48), IF hi (33), TA lo (45), TA hi (36), AC lo (72), AC hi (8), TI lo (38), TI hi (43), WBC lo (45), WBC hi (36), AS lo (45), AS hi (34), AH lo (49), AH hi (32). Clusters were ordered through hierarchical clustering of averaged variable gene expression values. Source data provided.

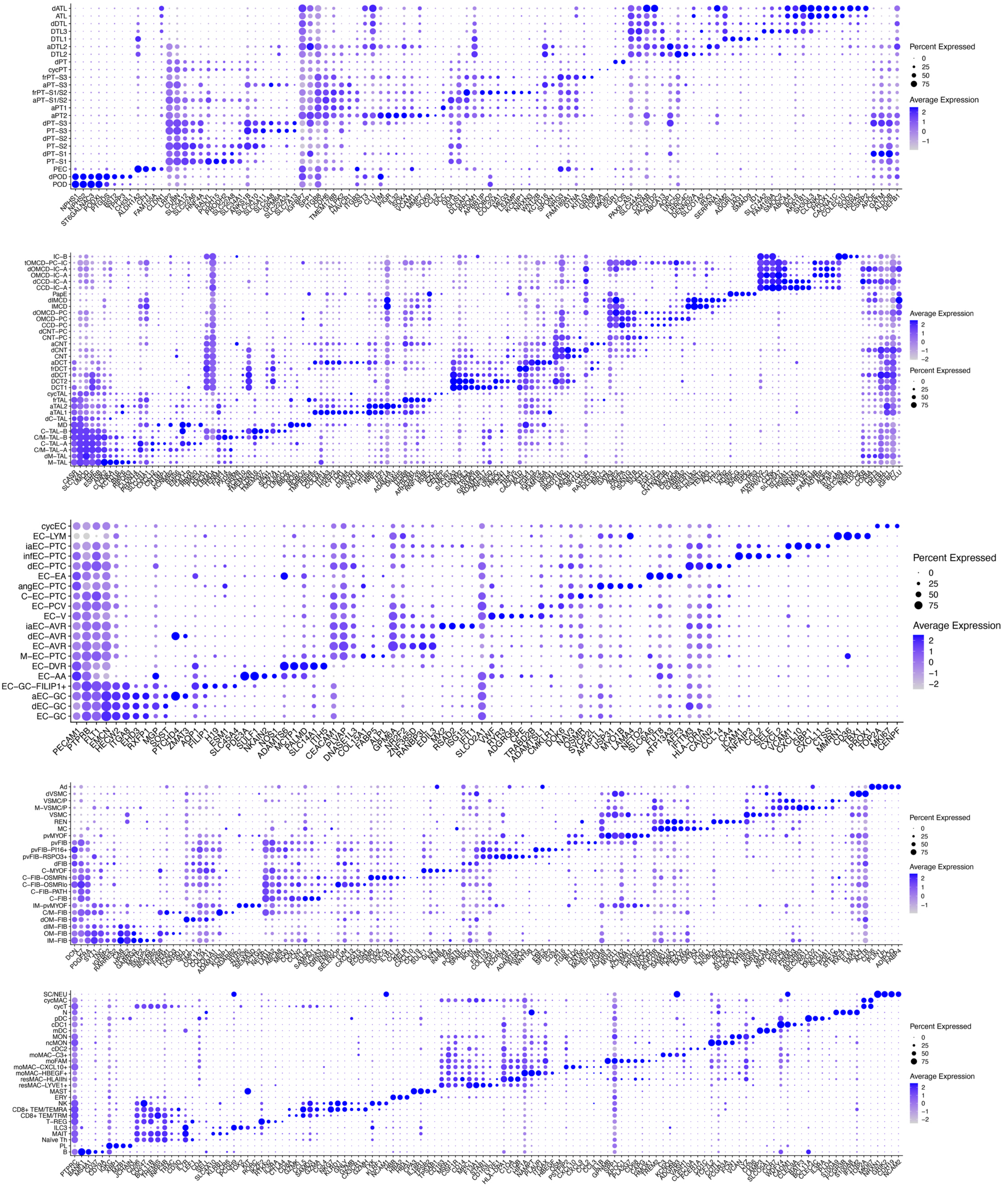
dATL
ATL
dDTL
DTL3
DTL1
aDTL2
DTL2
dPT
cycPT
frPT–S3
aPT–S3
frPT–S1/S2
aPT–S1/S2
aPT1
aPT2
dPT–S3
PT–S3
dPT–S2
PT–S2
dPT–S1
PT–S1
PEC
dPOD
POD
Percent Expressed
0
25
50
75
Average Expression
2
1
0
−1
IC–B
tOMCD–PC–IC
dOMCD–IC–A
OMCD–IC–A
dCCD–IC–A
CCD–IC–A
PapE
dIMCD
IMCD
dOMCD–PC
OMCD–PC
CCD–PC
dCNT–PC
CNT–PC
aCNT
dCNT
CNT
aDCT
frDCT
dDCT
DCT2
DCT1
cycTAL
frTAL
aTAL2
aTAL1
dC–TAL
MD
C–TAL–B
C/M–TAL–B
C–TAL–A
C/M–TAL–A
dM–TAL
M–TAL
Average Expression
2
1
0
−1
−2
Percent Expressed
0
25
50
75
cycEC
EC–LYM
iaEC–PTC
infEC–PTC
dEC–PTC
EC–EA
angEC–PTC
C–EC–PTC
EC–PCV
EC–V
iaEC–AVR
dEC–AVR
EC–AVR
M–EC–PTC
EC–DVR
EC–AA
EC–GC–FILIP1+
aEC–GC
dEC–GC
EC–GC
Percent Expressed
0
25
50
75
Average Expression
2
1
0
−1
−2
Ad
dVSMC
VSMC/P
M–VSMC/P
VSMC
REN
MC
pvMYOF
pvFIB
pvFIB–PI16+
pvFIB–RSPO3+
dFIB
C–MYOF
C–FIB–OSMRhi
C–FIB–OSMRlo
C–FIB–PATH
C–FIB
IM–pvMYOF
C/M–FIB
dOM–FIB
dIM–FIB
OM–FIB
IM–FIB
Percent Expressed
0
25
50
75
Average Expression
2
1
0
−1
SC/NEU
cycMAC
cycT
N
pDC
cDC1
mDC
MON
ncMON
cDC2
moMAC–C3+
moFAM
moMAC–CXCL10+
moMAC–HBEGF+
resMAC–HLAIIhi
resMAC–LYVE1+
MAST
ERY
NK
CD8+ TEM/TEMRA
CD8+ TEM/TRM
T–REG
ILC3
MAIT
Naïve Th
PL
B
Percent Expressed
0
25
50
75
Average Expression
2
1
0
−1
−2

**SD Figure 2. Cell type gene markers.** Dot plots showing average gene expression values for cell type markers genes across all subclasses.

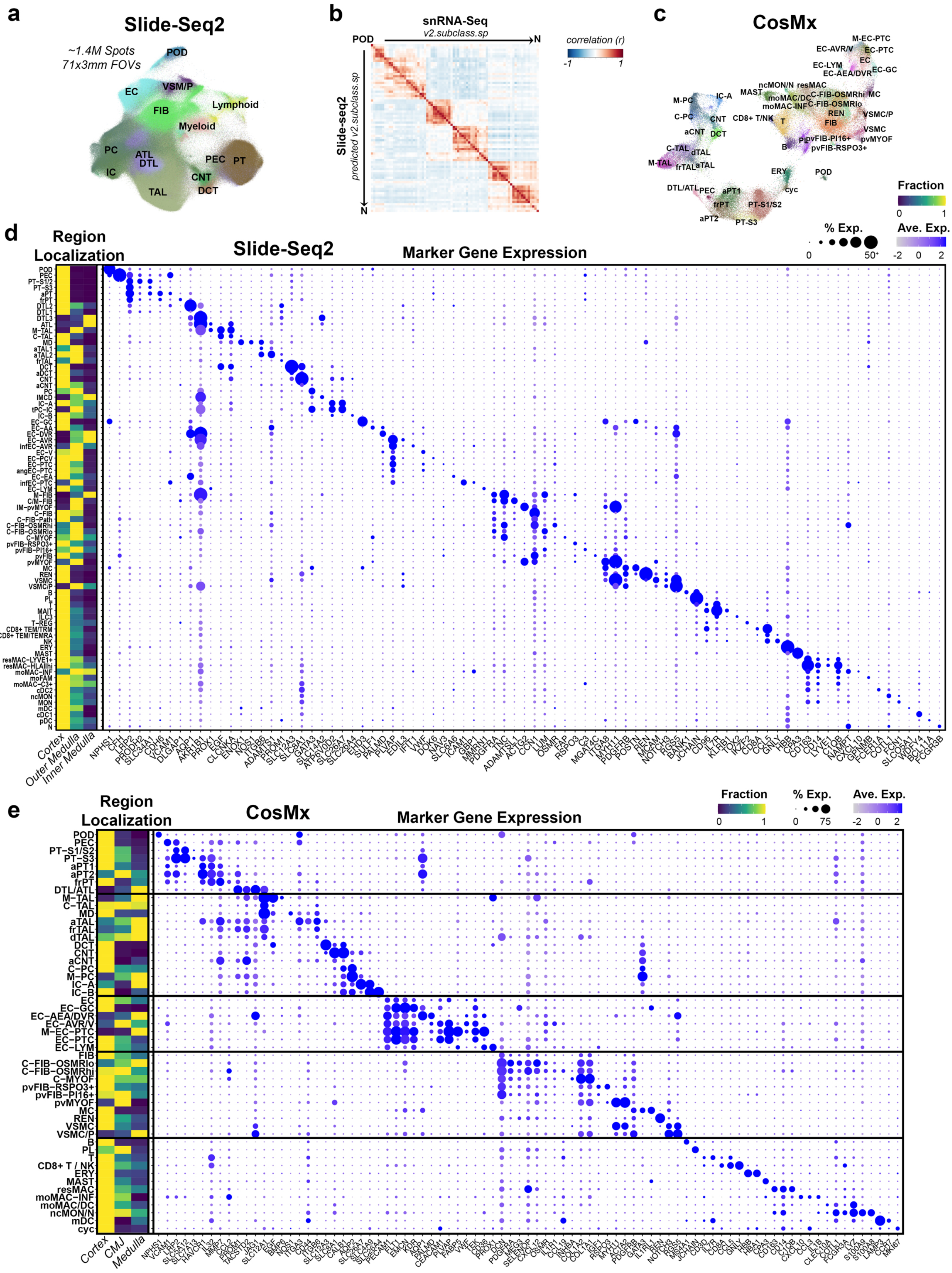

a
Slide-Seq2
~1.4M Spots
71x3mm FOVs
POD
EC
VSM/P
FIB
Lymphoid
Myeloid
PC
ATL
DTL
IC
PEC
PT
CNT
DCT
TAL
b
snRNA-Seq
v2.subclass.sp
POD
N
Slide-seq2
predicted v2.subclass.sp
correlation (r)
-1
1
c
CosMx
Fraction
0
1
% Exp.
Ave. Exp.
d
Region Localization
Slide-Seq2
Marker Gene Expression
Cortex
Outer Medulla
Inner Medulla
e
Region Localization
CosMx
Marker Gene Expression
Fraction
% Exp.
Ave. Exp.
Cortex
CMJ
Medulla

**SD Figure 3. Slide-seq2 and CosMx orthogonal validation and quality control. a**. UMAP embeddings for all Slide-seq2 spots showing broad subclass identities. **b**. Heatmap showing correlation of average scaled gene expression values for spatially resolved subclasses between Slide-seq2 and the reference single-nucleus RNA-seq data. **c**. UMAP embeddings for all CosMx cells showing final resolved subclasses. **d**. Left, heatmap of spatial enrichments for Slide-seq2 subclass abundances across cortex, outer medulla and inner medulla regions. Right, dotplot showing averaged gene expression values for cell type specific marker genes. **e**. Left, heatmap of spatial enrichments for CosMx subclass abundances across cortex, corticomedullary junction (CMJ) and medulla regions. Right, dotplot showing averaged gene expression values for cell type specific marker genes. Source data provided.

### SD Fig 4

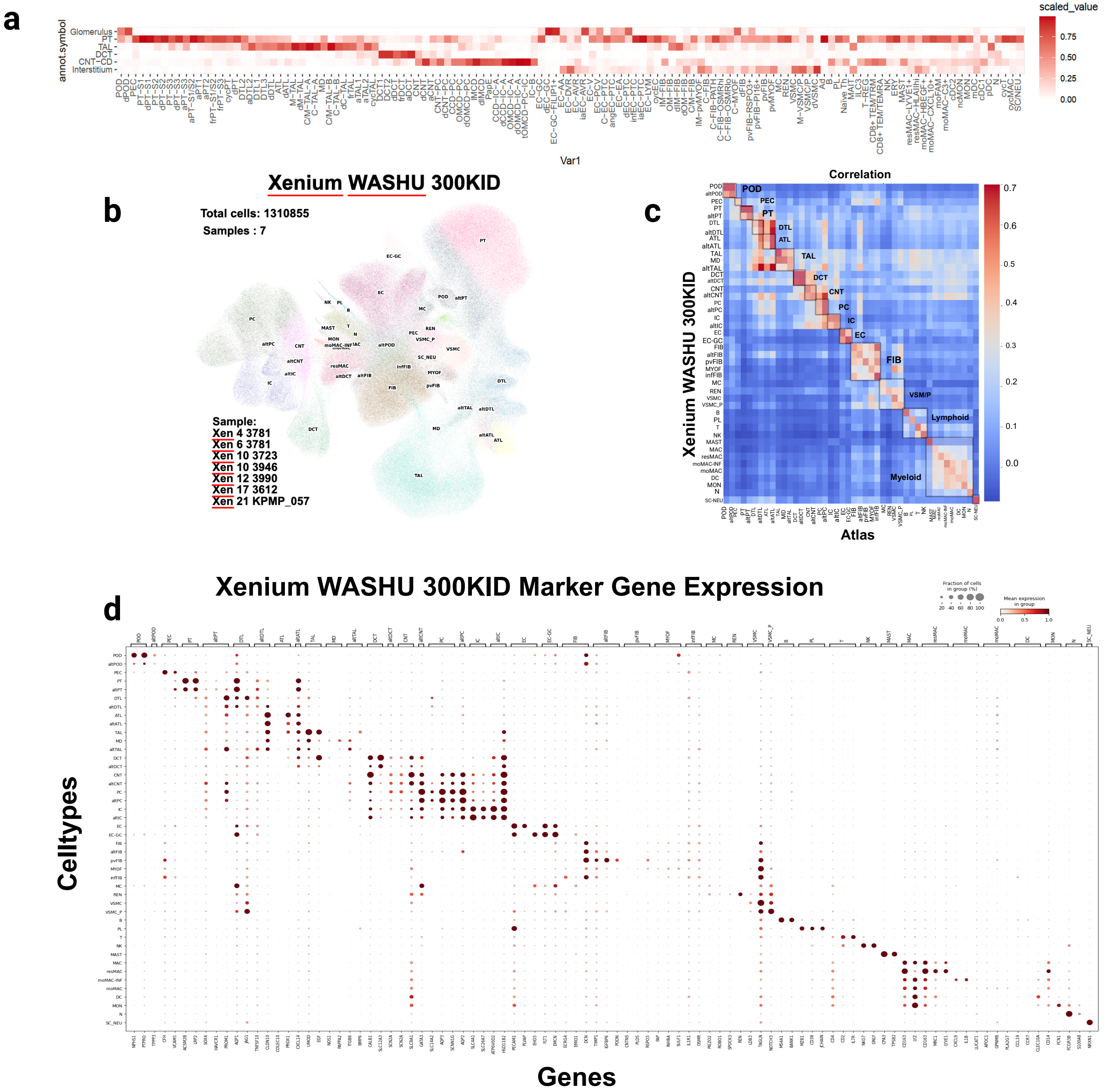
a
scaled_value
annot.symbol
Glomerulus
PT
TAL
DCT
CNT−CD
Interstitium
Var1
b
Xenium WASHU 300KID
Total cells: 1310855
Samples : 7
Sample:
Xen 4 3781
Xen 6 3781
Xen 10 3723
Xen 10 3946
Xen 12 3990
Xen 17 3612
Xen 21 KPMP_057
c
Correlation
POD
PEC
PT
DTL
ATL
TAL
DCT
CNT
PC
IC
EC
FIB
VSM/P
Lymphoid
Myeloid
Xenium WASHU 300KID
Atlas
d
Xenium WASHU 300KID Marker Gene Expression
Fraction of cells in group (%)
Mean expression in group
Celltypes
Genes

**SD Figure 4. Visium and Xenium orthogonal validation and quality control. a.** Heatmap presenting mapping proportions over histologically validated spots using Visium on OCT frozen samples. **b**. UMAP embeddings show the cell type populations identified on the integrated object for the Xenium WASHU 300KID Panel. **c.** Correlation Plot showing the similarity between the cell types identified on the xenium data and the atlas HKAv2 object. Cell types were grouped and ordered based on biological relevance. **d**. Dot plot showing the FTU/cell type-specific marker gene expression of the genes chosen on the Xenium panel. Dot size indicates the fraction of cells within that cell type, and dot color intensity represents mean expression within the group.  Created in BioRender. Jain, S. (2025) https://BioRender.com/hl0fdn6. Source data provided.

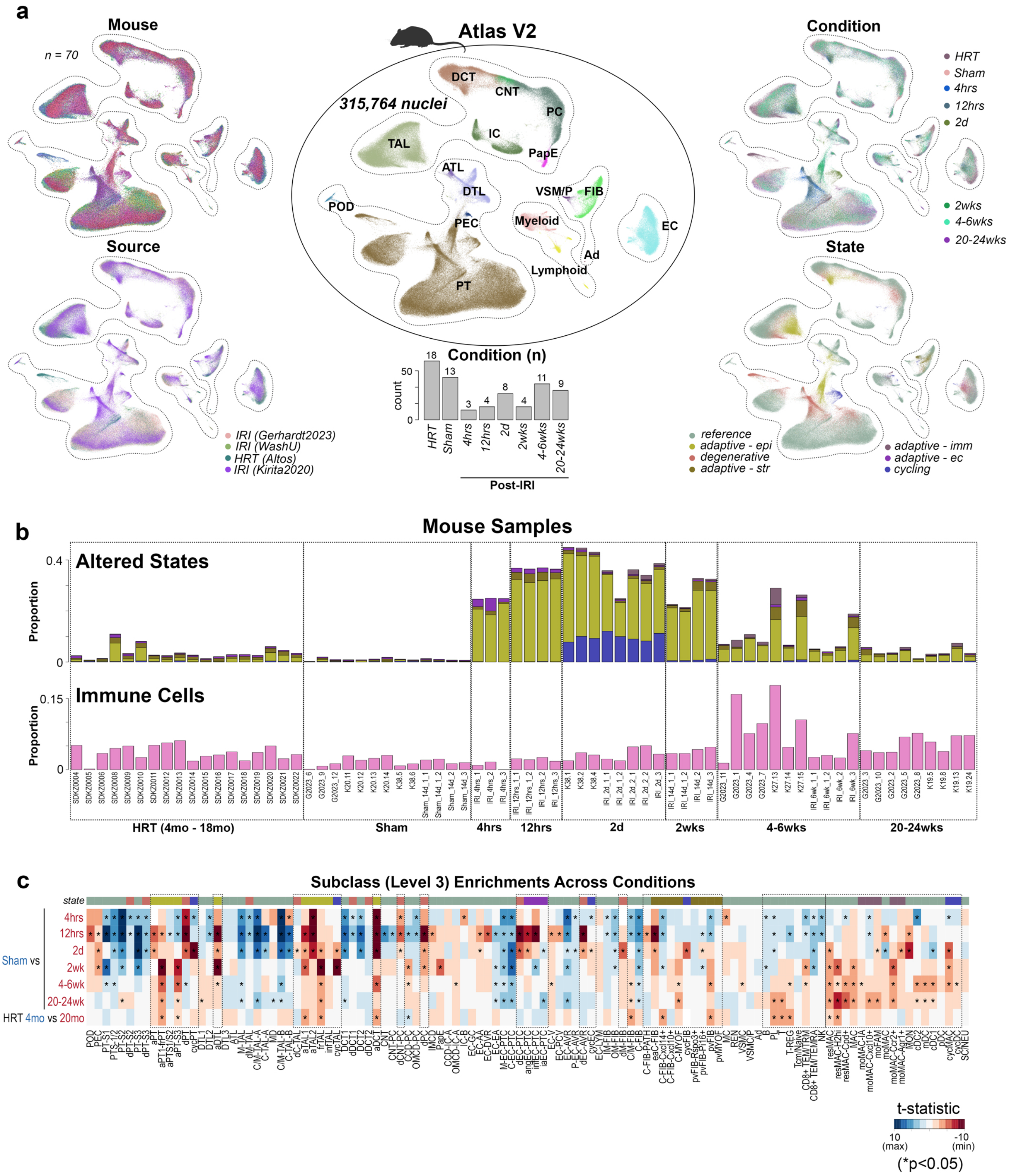
a
Mouse
n = 70
Source
IRI (Gerhardt2023)
IRI (WashU)
HRT (Altos)
IRI (Kirita2020)
Atlas V2
315,764 nuclei
DCT
CNT
PC
IC
PapE
TAL
ATL
DTL
POD
PEC
VSM/P
FIB
Myeloid
EC
Ad
Lymphoid
PT
Condition (n)
count
HRT
Sham
4hrs
12hrs
2d
2wks
4-6wks
20-24wks
Post-IRI
Condition
State
reference
adaptive - epi
degenerative
adaptive - str
adaptive - imm
adaptive - ec
cycling
b
Mouse Samples
Altered States
Immune Cells
Proportion
HRT (4mo - 18mo)
Sham
4hrs
12hrs
2d
2wks
4-6wks
20-24wks
c
Subclass (Level 3) Enrichments Across Conditions
state
Sham vs
4hrs
12hrs
2d
2wk
4-6wk
20-24wk
HRT 4mo vs 20mo
t-statistic
10 (max)
-10 (min)
(*p<0.05)

**SD Figure 5. Mouse atlas quality control. a**. UMAP embeddings showing broad subclasses (center plot) and the associated distribution of mouse samples, tissue sources, injury conditions and altered states (outer plots). Barplot shows number of mice sampled for each condition category. **b**. Barplots showing the proportion of altered states (top) as indicated in (a) and the proportion of immune cells (bottom) across the different mice and conditions. **c**. Heatmap showing cell type enrichments as t statistics for the comparison of subclass abundances between two different condition groupings. Asterisk indicates p values less than 0.05.

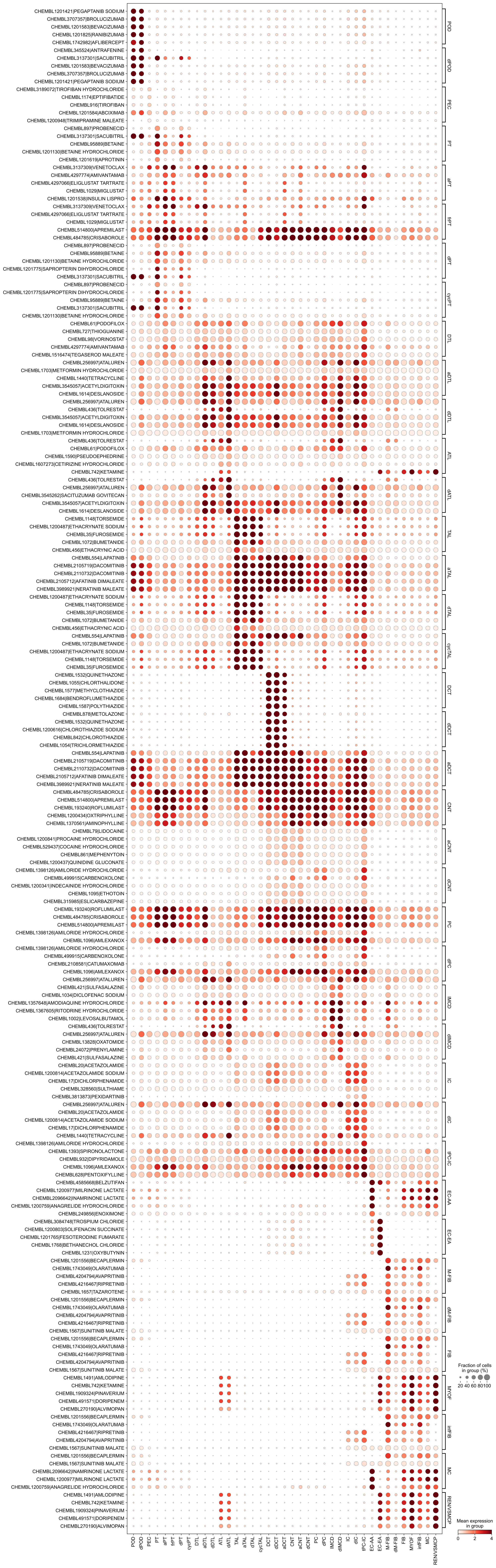

CHEMBL1201421|PEGAPTANIB SODIUM
CHEMBL3707357|BROLUCIZUMAB
CHEMBL1201583|BEVACIZUMAB
CHEMBL1201825|RANIBIZUMAB
CHEMBL1742982|AFLIBERCEPT
CHEMBL345524|ANTRAFENINE
CHEMBL3137301|SACUBITRIL
CHEMBL1201583|BEVACIZUMAB
CHEMBL3707357|BROLUCIZUMAB
CHEMBL1201421|PEGAPTANIB SODIUM
CHEMBL3189072|TIROFIBAN HYDROCHLORIDE
CHEMBL1174|EPTIFIBATIDE
CHEMBL916|TIROFIBAN
CHEMBL1201584|ABCIXIMAB
CHEMBL1200948|TRIMIPRAMINE MALEATE
CHEMBL897|PROBENECID
CHEMBL3137301|SACUBITRIL
CHEMBL95889|BETAINE
CHEMBL1201130|BETAINE HYDROCHLORIDE
CHEMBL1201619|APROTININ
CHEMBL3137309|VENETOCLAX
CHEMBL4297774|AMIVANTAMAB
CHEMBL4297066|ELIGLUSTAT TARTRATE
CHEMBL1029|MIGLUSTAT
CHEMBL1201538|INSULIN LISPRO
CHEMBL3137309|VENETOCLAX
CHEMBL4297066|ELIGLUSTAT TARTRATE
CHEMBL1029|MIGLUSTAT
CHEMBL514800|APREMILAST
CHEMBL484785|CRISABOROLE
CHEMBL897|PROBENECID
CHEMBL95889|BETAINE
CHEMBL1201130|BETAINE HYDROCHLORIDE
CHEMBL1201775|SAPROPTERIN DIHYDROCHLORIDE
CHEMBL3137301|SACUBITRIL
CHEMBL897|PROBENECID
CHEMBL1201775|SAPROPTERIN DIHYDROCHLORIDE
CHEMBL95889|BETAINE
CHEMBL3137301|SACUBITRIL
CHEMBL1201130|BETAINE HYDROCHLORIDE
CHEMBL61|PODOFILOX
CHEMBL727|THIOGUANINE
CHEMBL98|VORINOSTAT
CHEMBL4297774|AMIVANTAMAB
CHEMBL1516474|TEGASEROD MALEATE
CHEMBL256997|ATALUREN
CHEMBL1703|METFORMIN HYDROCHLORIDE
CHEMBL1440|TETRACYCLINE
CHEMBL3545057|ACETYLDIGITOXIN
CHEMBL1614|DESLANOSIDE
CHEMBL256997|ATALUREN
CHEMBL436|TOLRESTAT
CHEMBL3545057|ACETYLDIGITOXIN
CHEMBL1614|DESLANOSIDE
CHEMBL1703|METFORMIN HYDROCHLORIDE
CHEMBL436|TOLRESTAT
CHEMBL61|PODOFILOX
CHEMBL1590|PSEUDOEPHEDRINE
CHEMBL1607273|CETIRIZINE HYDROCHLORIDE
CHEMBL742|KETAMINE
CHEMBL436|TOLRESTAT
CHEMBL256997|ATALUREN
CHEMBL3545262|SACITUZUMAB GOVITECAN
CHEMBL3545057|ACETYLDIGITOXIN
CHEMBL1614|DESLANOSIDE
CHEMBL1148|TORSEMIDE
CHEMBL1200487|ETHACRYNATE SODIUM
CHEMBL35|FUROSEMIDE
CHEMBL1072|BUMETANIDE
CHEMBL456|ETHACRYNIC ACID
CHEMBL554|LAPATINIB
CHEMBL2105719|DACOMITINIB
CHEMBL2110732|DACOMITINIB
CHEMBL2105712|AFATINIB DIMALEATE
CHEMBL3989921|NERATINIB MALEATE
CHEMBL1200487|ETHACRYNATE SODIUM
CHEMBL1148|TORSEMIDE
CHEMBL35|FUROSEMIDE
CHEMBL1072|BUMETANIDE
CHEMBL456|ETHACRYNIC ACID
CHEMBL554|LAPATINIB
CHEMBL1072|BUMETANIDE
CHEMBL1200487|ETHACRYNATE SODIUM
CHEMBL1148|TORSEMIDE
CHEMBL35|FUROSEMIDE
CHEMBL1532|QUINETHAZONE
CHEMBL1055|CHLORTHALIDONE
CHEMBL1577|METHYCLOTHIAZIDE
CHEMBL1684|BENDROFLUMETHIAZIDE
CHEMBL1587|POLYTHIAZIDE
CHEMBL878|METOLAZONE
CHEMBL1532|QUINETHAZONE
CHEMBL1200616|CHLOROTHIAZIDE SODIUM
CHEMBL842|CHLOROTHIAZIDE
CHEMBL1054|TRICHLORMETHIAZIDE
CHEMBL554|LAPATINIB
CHEMBL2105719|DACOMITINIB
CHEMBL2110732|DACOMITINIB
CHEMBL2105712|AFATINIB DIMALEATE
CHEMBL3989921|NERATINIB MALEATE
CHEMBL484785|CRISABOROLE
CHEMBL514800|APREMILAST
CHEMBL193240|ROFLUMILAST
CHEMBL1200434|OXTRIPHYLLINE
CHEMBL1370561|AMINOPHYLLINE
CHEMBL79|LIDOCAINE
CHEMBL1200841|PROCAINE HYDROCHLORIDE
CHEMBL529437|COCAINE HYDROCHLORIDE
CHEMBL861|MEPHENYTOIN
CHEMBL1200437|QUINIDINE GLUCONATE
CHEMBL1398126|AMILORIDE HYDROCHLORIDE
CHEMBL499915|CARBENOXOLONE
CHEMBL1200341|INDECAINIDE HYDROCHLORIDE
CHEMBL1095|ETHOTOIN
CHEMBL315985|ESLICARBAZEPINE
CHEMBL193240|ROFLUMILAST
CHEMBL484785|CRISABOROLE
CHEMBL514800|APREMILAST
CHEMBL1398126|AMILORIDE HYDROCHLORIDE
CHEMBL1096|AMLEXANOX
CHEMBL1398126|AMILORIDE HYDROCHLORIDE
CHEMBL499915|CARBENOXOLONE
CHEMBL2108581|CATUMAXOMAB
CHEMBL1096|AMLEXANOX
CHEMBL256997|ATALUREN
CHEMBL421|SULFASALAZINE
CHEMBL1034|DICLOFENAC SODIUM
CHEMBL1357648|AMODIAQUINE HYDROCHLORIDE
CHEMBL1367605|RITODRINE HYDROCHLORIDE
CHEMBL1002|LEVOSALBUTAMOL
CHEMBL436|TOLRESTAT
CHEMBL256997|ATALUREN
CHEMBL13828|OXATOMIDE
CHEMBL24072|PRENYLAMINE
CHEMBL421|SULFASALAZINE
CHEMBL20|ACETAZOLAMIDE
CHEMBL1200814|ACETAZOLAMIDE SODIUM
CHEMBL17|DICHLORPHENAMIDE
CHEMBL328560|SULTHIAME
CHEMBL3813873|PEXIDARTINIB
CHEMBL256997|ATALUREN
CHEMBL20|ACETAZOLAMIDE
CHEMBL1200814|ACETAZOLAMIDE SODIUM
CHEMBL17|DICHLORPHENAMIDE
CHEMBL1440|TETRACYCLINE
CHEMBL1398126|AMILORIDE HYDROCHLORIDE
CHEMBL1393|SPIRONOLACTONE
CHEMBL932|DIPYRIDAMOLE
CHEMBL1096|AMLEXANOX
CHEMBL628|PENTOXIFYLLINE
CHEMBL4585668|BELZUTIFAN
CHEMBL1200977|MILRINONE LACTATE
CHEMBL2096642|INAMRINONE LACTATE
CHEMBL1200759|ANAGRELIDE HYDROCHLORIDE
CHEMBL249856|ENOXIMONE
CHEMBL3084748|TROSPIUM CHLORIDE
CHEMBL1200803|SOLIFENACIN SUCCINATE
CHEMBL1201765|FESOTERODINE FUMARATE
CHEMBL1768|BETHANECHOL CHLORIDE
CHEMBL1231|OXYBUTYNIN
CHEMBL1201556|BECAPLERMIN
CHEMBL1743049|OLARATUMAB
CHEMBL4204794|AVAPRITINIB
CHEMBL4216467|RIPRETINIB
CHEMBL1657|TAZAROTENE
CHEMBL1201556|BECAPLERMIN
CHEMBL1743049|OLARATUMAB
CHEMBL4204794|AVAPRITINIB
CHEMBL4216467|RIPRETINIB
CHEMBL1567|SUNITINIB MALATE
CHEMBL1201556|BECAPLERMIN
CHEMBL1743049|OLARATUMAB
CHEMBL4216467|RIPRETINIB
CHEMBL4204794|AVAPRITINIB
CHEMBL1567|SUNITINIB MALATE
CHEMBL1491|AMLODIPINE
CHEMBL742|KETAMINE
CHEMBL1909324|PINAVERIUM
CHEMBL491571|DORIPENEM
CHEMBL270190|ALVIMOPAN
CHEMBL1201556|BECAPLERMIN
CHEMBL1743049|OLARATUMAB
CHEMBL4216467|RIPRETINIB
CHEMBL4204794|AVAPRITINIB
CHEMBL1567|SUNITINIB MALATE
CHEMBL1201556|BECAPLERMIN
CHEMBL1567|SUNITINIB MALATE
CHEMBL2096642|INAMRINONE LACTATE
CHEMBL1200977|MILRINONE LACTATE
CHEMBL1200759|ANAGRELIDE HYDROCHLORIDE
CHEMBL1491|AMLODIPINE
CHEMBL742|KETAMINE
CHEMBL1909324|PINAVERIUM
CHEMBL491571|DORIPENEM
CHEMBL270190|ALVIMOPAN
POD
dPOD
PEC
PT
aPT
frPT
dPT
cycPT
DTL
aDTL
dDTL
ATL
dATL
TAL
aTAL
dTAL
cycTAL
DCT
dDCT
aDCT
CNT
aCNT
dCNT
PC
dPC
IMCD
dIMCD
IC
dIC
tPC-IC
EC-AA
EC-EA
M-FIB
dM-FIB
FIB
MYOF
infFIB
MC
REN/VSMCP
Fraction of cells in group (%)
20 40 60 80 100
Mean expression in group
0 2 4

**SD Figure 6. Druggable Targets Dotplot.** Dotplot showing predicted drug–cell type associations as inferred by Drug2cell, which scores the expression of drug target genes across annotated cell populations from our kidney atlas v2. Each dot represents a specific drug–cell type pair. Dot size indicates the fraction of cells within that cell type expressing the drug's target genes, and dot color intensity represents mean expression of those targets within the group. Cell types were grouped and ordered based on biological relevance. Dotplot is shown using a fixed color scale to enable fair comparisons across plots.

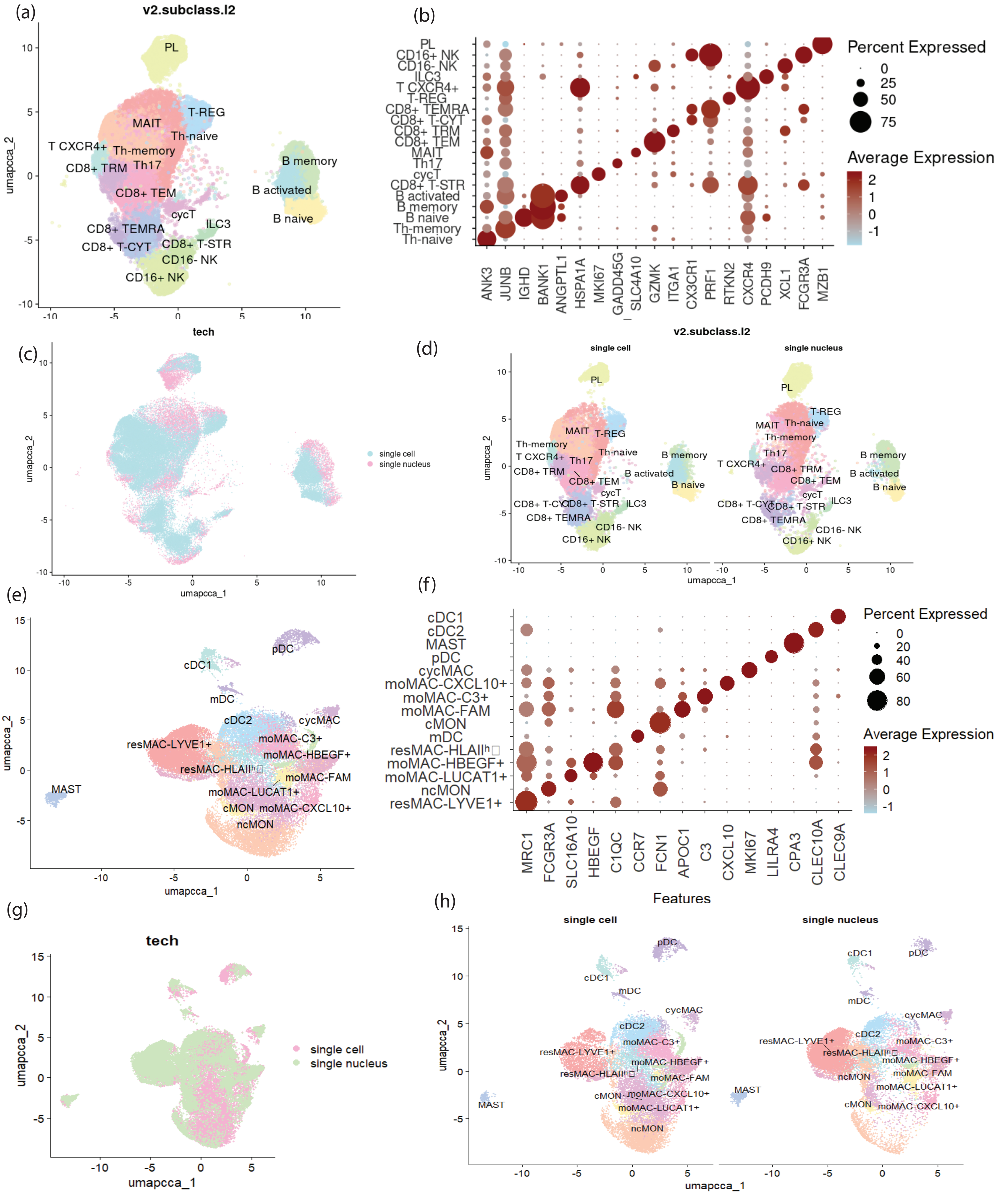
(a)
v2.subclass.l2
umapcca_2
PL
T-REG
MAIT
Th-naive
T CXCR4+
Th-memory
CD8+ TRM
Th17
B memory
B activated
B naive
CD8+ TEM
cycT
CD8+ TEMRA
ILC3
CD8+ T-CYT
CD8+ T-STR
CD16- NK
CD16+ NK
(b)
Percent Expressed
0
25
50
75
Average Expression
ANK3
JUNB
IGHD
BANK1
ANGPTL1
HSPA1A
MKI67
GADD45G
SLC4A10
GZMK
ITGA1
CX3CR1
PRF1
RTKN2
CXCR4
PCDH9
XCL1
FCGR3A
MZB1
tech
(c)
single cell
single nucleus
umapcca_1
(d)
single cell
single nucleus
(e)
pDC
cDC1
mDC
cDC2
cycMAC
moMAC-C3+
resMAC-LYVE1+
moMAC-HBEGF+
resMAC-HLAIIʰ
moMAC-FAM
MAST
moMAC-LUCAT1+
cMON
moMAC-CXCL10+
ncMON
(f)
Percent Expressed
0
20
40
60
80
Average Expression
MRC1
FCGR3A
SLC16A10
HBEGF
C1QC
CCR7
FCN1
APOC1
C3
CXCL10
MKI67
LILRA4
CPA3
CLEC10A
CLEC9A
Features
(g)
tech
single cell
single nucleus
(h)
single cell
single nucleus

**SD Figure 7. SD Figure 7. Integration of single cell/single nucleus myeloid and lymphoid cell populations.** a. UMAP embedding of myeloid cell types in single cell and single nucleus data. b. Dot plot showing the top markers of myeloid cell types identified in the integrated myeloid dataset c. UMAP embedding of myeloid cells in single cell and single nucleus data. d. UMAP embedding of myeloid cell types in single cell and single nucleus data shown separately. e. UMAP embedding of myeloid cell types in single cell and single nucleus data. f. Dot plot showing the top markers of lymphoid cell types identified in the integrated lymphoid dataset g. UMAP embedding of lymphoid cells in single cell and single nucleus data. h. UMAP embedding of lymphoid cell types in single cell and single nucleus data shown separately.

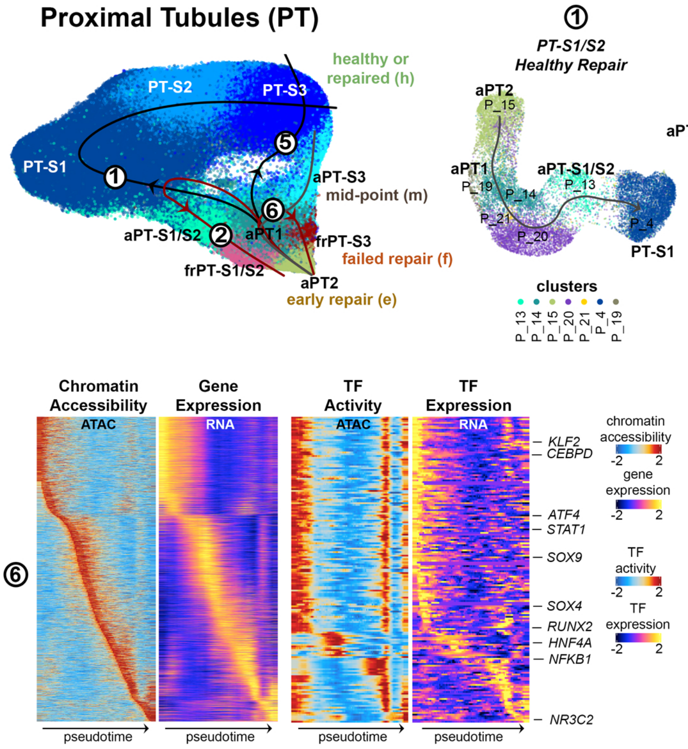

Proximal Tubules (PT)
healthy or repaired (h)
PT-S2
PT-S3
PT-S1
aPT-S3
mid-point (m)
aPT-S1/S2
aPT1
frPT-S3
failed repair (f)
frPT-S1/S2
aPT2
early repair (e)
PT-S1/S2 Healthy Repair
PT-S1/S2 Failed Repair
PT-S3 Healthy Repair
PT-S3 Failed Repair
clusters
Chromatin Accessibility
ATAC
Gene Expression
RNA
TF Activity
ATAC
TF Expression
RNA
KLF2
CEBPD
ATF4
STAT1
SOX9
SOX4
RUNX2
HNF4A
NFKB1
NR3C2
chromatin accessibility
gene expression
TF activity
TF expression
pseudotime

**SD Figure 8. Epithelial repair states.** **a**. UMAP embedding showing proximal tubule (PT) states and associated Slingshot predicted trajectories. **b**. UMAP embeddings for individual trajectories shown in (**a**) colored by the associated cluster identities. PT clusters were grouped into subclasses in part based on their localization along these trajectories (early, mid and late subsets). **c**. Left, scMEGA heatmaps showing chromatin accessibility and associated expression for the top pseudotime-variable genes. Right, heatmaps showing both the binding site activity and corresponding gene expression values for select TFs identified across the same trajectory. These formed the basis for trajectory-associated gene regulatory networks (GRNs).

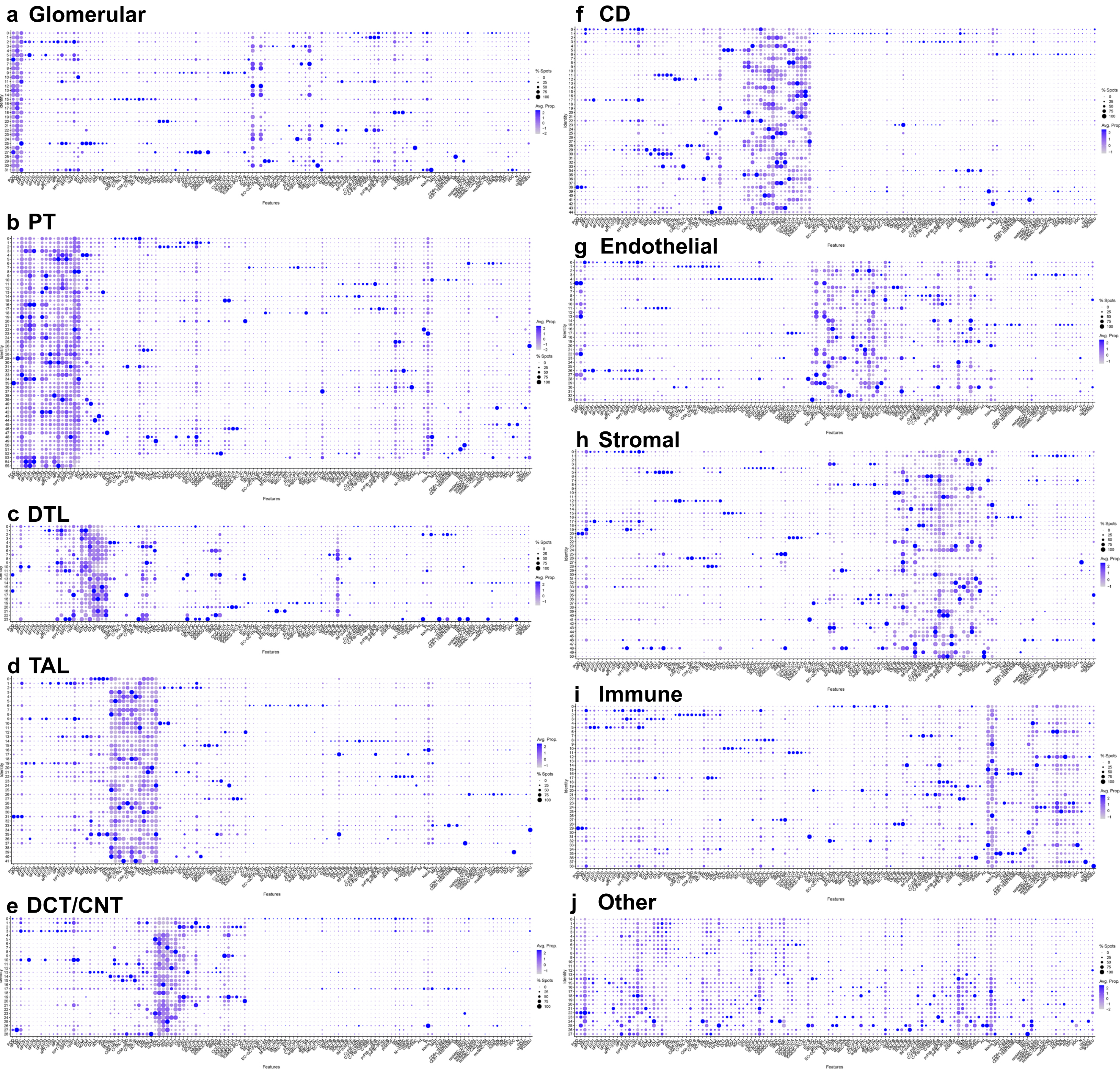

a Glomerular
b PT
c DTL
d TAL
e DCT/CNT
f CD
g Endothelial
h Stromal
i Immune
j Other
Features
Identity
% Spots
Avg. Prop.

**SD Figure 9. Visium Niches.** Dotplots displaying the cell type composition of individual niches for the groups **a.** Glomerular **b.** PT **c.** DTL **d.** TAL **e.** DCT/CNT **f.** CD **g.** Endothelial cells **h.** Stromal cells **i.** Immune cells **j.** Other.

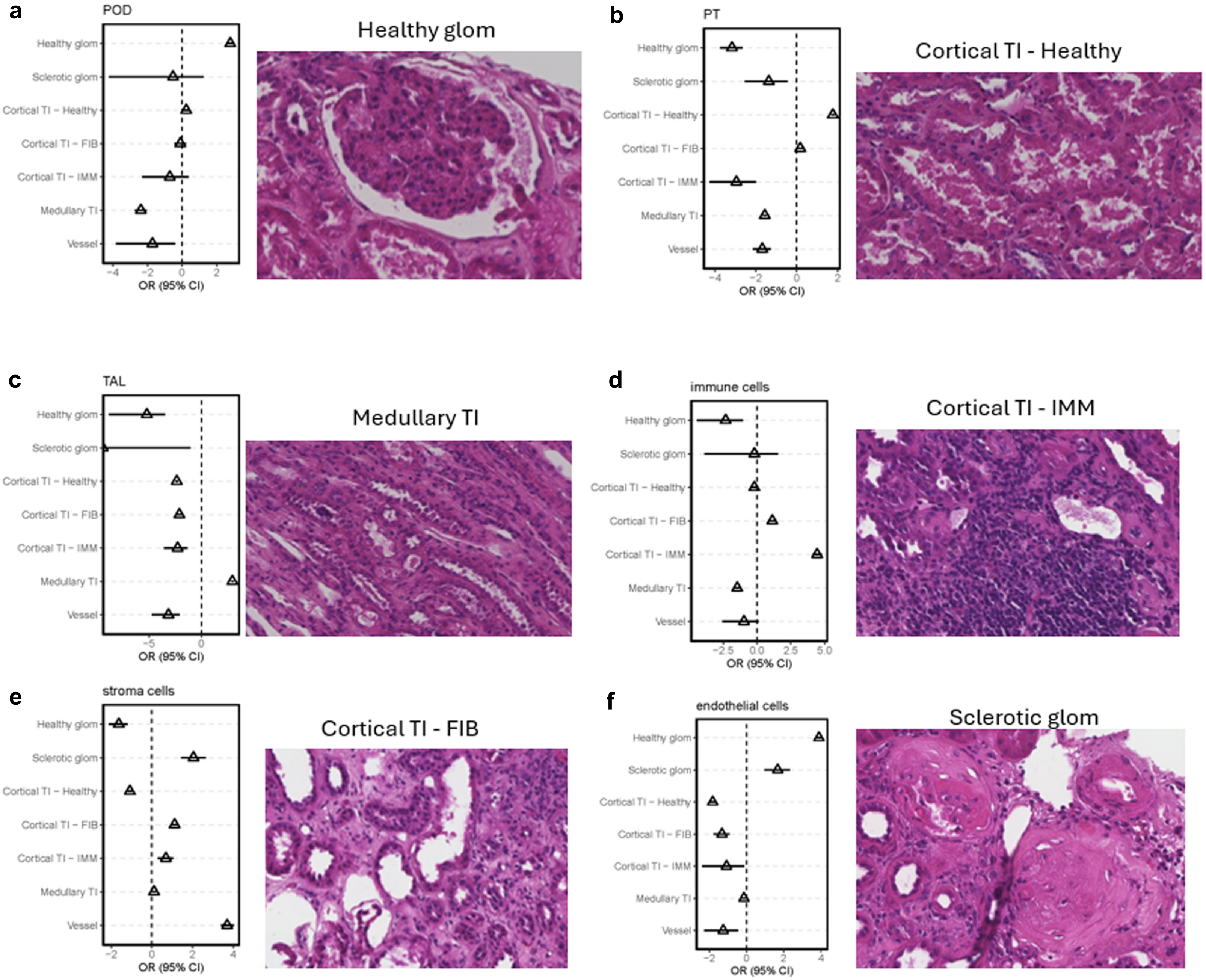
a
POD
Healthy glom
Sclerotic glom
Cortical TI – Healthy
Cortical TI – FIB
Cortical TI – IMM
Medullary TI
Vessel
OR (95% CI)
Healthy glom
b
PT
OR (95% CI)
Cortical TI - Healthy
c
TAL
OR (95% CI)
Medullary TI
d
immune cells
OR (95% CI)
Cortical TI - IMM
e
stroma cells
OR (95% CI)
Cortical TI - FIB
f
endothelial cells
OR (95% CI)
Sclerotic glom

**SD Figure 10. Mapping of cell type signatures to annotated histology.** Forest plot showing odds ratio of niche mapping over histologically annotated regions grouped by main cell type signature. A representative example of a histologically annotated region is shown. **a.** Niches with podocyte signatures are associated with healthy glomeruli. **b.** Niches with proximal tubule signatures are associated with healthy cortical tubulo-interstitium. **c.** Niches with thick ascending limb signatures are associated with medullary tubulo-interstitium. **d.** Niches with immune cell signatures are associated with tubulo-interstitium and immune infiltration. **e.** Niches with stromal signatures are associated with tubulo-interstitium and fibrosis. **f.** Niches with endothelial cell signatures are associated with sclerotic glomeruli. Source data provided.

### SD Fig 11

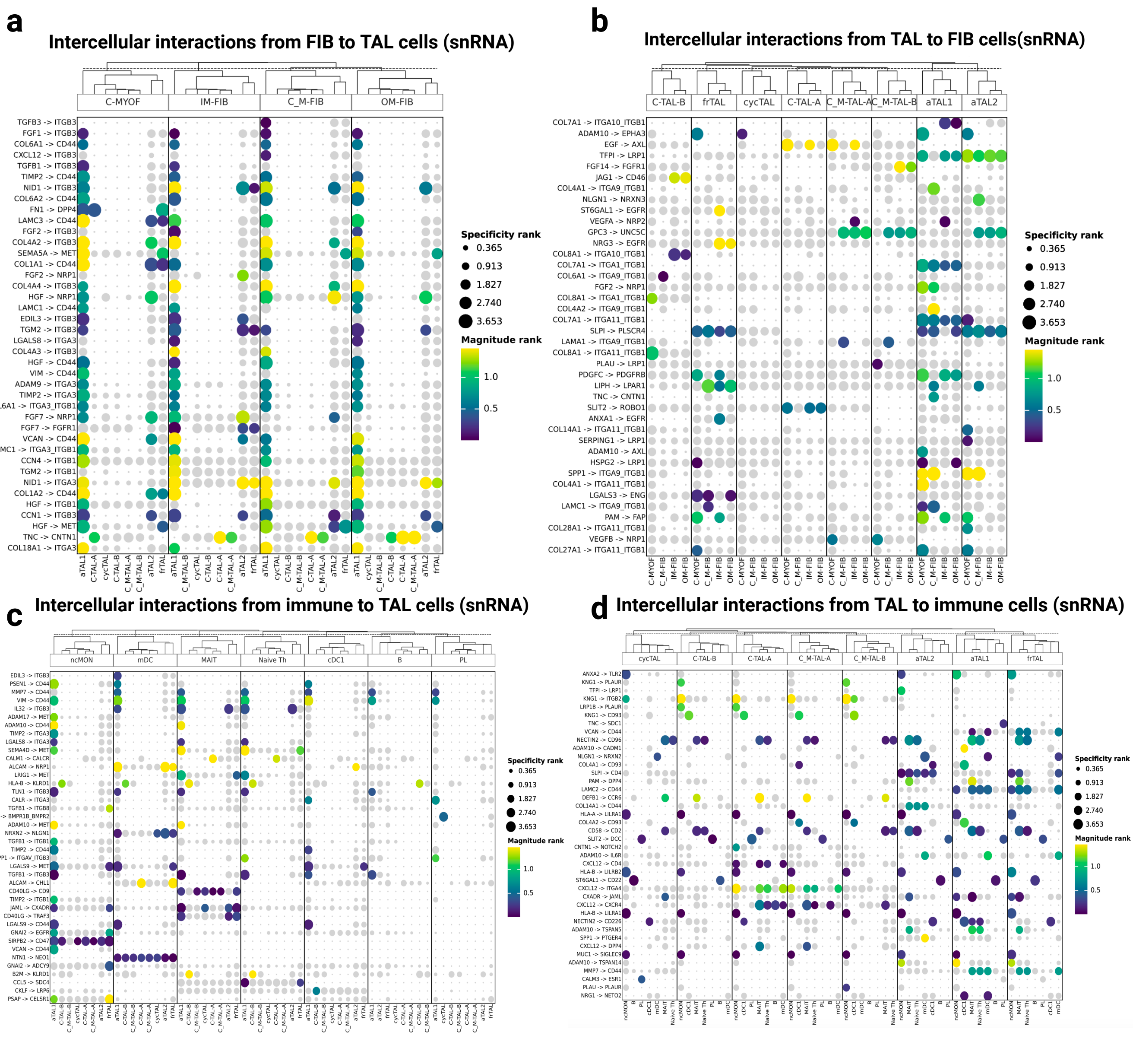
a
Intercellular interactions from FIB to TAL cells (snRNA)
C-MYOF
IM-FIB
C_M-FIB
OM-FIB
TGFB3 -> ITGB3
FGF1 -> ITGB3
COL6A1 -> CD44
CXCL12 -> ITGB3
TGFB1 -> ITGB3
TIMP2 -> CD44
NID1 -> ITGB3
COL6A2 -> CD44
FN1 -> DPP4
LAMC3 -> CD44
FGF2 -> ITGB3
COL4A2 -> ITGB3
SEMA5A -> MET
COL1A1 -> CD44
FGF2 -> NRP1
COL4A4 -> ITGB3
HGF -> NRP1
LAMC1 -> CD44
EDIL3 -> ITGB3
TGM2 -> ITGB3
LGALS8 -> ITGA3
COL4A3 -> ITGB3
HGF -> CD44
VIM -> CD44
ADAM9 -> ITGA3
TIMP2 -> ITGA3
COL6A1 -> ITGA3_ITGB1
FGF7 -> NRP1
FGF7 -> FGFR1
VCAN -> CD44
LAMC1 -> ITGA3_ITGB1
CCN4 -> ITGB1
TGM2 -> ITGB1
NID1 -> ITGA3
COL1A2 -> CD44
HGF -> ITGB1
CCN1 -> ITGB3
HGF -> MET
TNC -> CNTN1
COL18A1 -> ITGA3
aTAL1
C-TAL-A
cycTAL
C-TAL-B
C_M-TAL-A
C_M-TAL-B
aTAL2
frTAL
Specificity rank
0.365
0.913
1.827
2.740
3.653
Magnitude rank
1.0
0.5
b
Intercellular interactions from TAL to FIB cells(snRNA)
C-TAL-B
frTAL
cycTAL
C-TAL-A
C_M-TAL-A
C_M-TAL-B
aTAL1
aTAL2
COL7A1 -> ITGA10_ITGB1
ADAM10 -> EPHA3
EGF -> AXL
TFPI -> LRP1
FGF14 -> FGFR1
JAG1 -> CD46
COL4A1 -> ITGA9_ITGB1
NLGN1 -> NRXN3
ST6GAL1 -> EGFR
VEGFA -> NRP2
GPC3 -> UNC5C
NRG3 -> EGFR
COL8A1 -> ITGA10_ITGB1
COL7A1 -> ITGA1_ITGB1
COL6A1 -> ITGA9_ITGB1
FGF2 -> NRP1
COL8A1 -> ITGA1_ITGB1
COL4A2 -> ITGA9_ITGB1
COL7A1 -> ITGA11_ITGB1
SLPI -> PLSCR4
LAMA1 -> ITGA9_ITGB1
COL8A1 -> ITGA11_ITGB1
PLAU -> LRP1
PDGFC -> PDGFRB
LIPH -> LPAR1
TNC -> CNTN1
SLIT2 -> ROBO1
ANXA1 -> EGFR
COL14A1 -> ITGA11_ITGB1
SERPING1 -> LRP1
ADAM10 -> AXL
HSPG2 -> LRP1
SPP1 -> ITGA9_ITGB1
COL4A1 -> ITGA11_ITGB1
LGALS3 -> ENG
LAMC1 -> ITGA9_ITGB1
PAM -> FAP
COL28A1 -> ITGA11_ITGB1
VEGFB -> NRP1
COL27A1 -> ITGA11_ITGB1
C-MYOF
C_M-FIB
IM-FIB
OM-FIB
Specificity rank
0.365
0.913
1.827
2.740
3.653
Magnitude rank
1.0
0.5
c
Intercellular interactions from immune to TAL cells (snRNA)
ncMON
mDC
MAIT
Naive Th
cDC1
B
PL
EDIL3 -> ITGB3
PSEN1 -> CD44
MMP7 -> CD44
VIM -> CD44
IL32 -> ITGB3
ADAM17 -> MET
ADAM10 -> CD44
TIMP2 -> ITGA3
LGALS8 -> ITGA3
SEMA4D -> MET
CALM1 -> CALCR
ALCAM -> NRP1
LRIG1 -> MET
HLA-B -> KLRD1
TLN1 -> ITGB3
CALR -> ITGA3
TGFB1 -> ITGB8
BMP6 -> BMPR1B_BMPR2
ADAM10 -> MET
NRXN2 -> NLGN1
TGFB1 -> ITGB1
TIMP2 -> CD44
SPP1 -> ITGAV_ITGB3
LGALS9 -> MET
TGFB1 -> ITGB3
ALCAM -> CHL1
CD40LG -> CD9
TIMP2 -> ITGB1
JAML -> CXADR
CD40LG -> TRAF3
LGALS9 -> CD44
GNAI2 -> EGFR
SIRPB2 -> CD47
VCAN -> CD44
NTN1 -> NEO1
GNAI2 -> ADCY9
B2M -> KLRD1
CCL5 -> SDC4
CKLF -> LRP6
PSAP -> CELSR1
aTAL1
C-TAL-B
C_M-TAL-B
cycTAL
C-TAL-A
C_M-TAL-A
aTAL2
frTAL
Specificity rank
0.365
0.913
1.827
2.740
3.653
Magnitude rank
1.0
0.5
d
Intercellular interactions from TAL to immune cells (snRNA)
cycTAL
C-TAL-B
C-TAL-A
C_M-TAL-A
C_M-TAL-B
aTAL2
aTAL1
frTAL
ANXA2 -> TLR2
KNG1 -> PLAUR
TFPI -> LRP1
KNG1 -> ITGB2
LRP1B -> PLAUR
KNG1 -> CD93
TNC -> SDC1
VCAN -> CD44
NECTIN2 -> CD96
ADAM10 -> CADM1
NLGN1 -> NRXN2
COL4A1 -> CD93
SLPI -> CD4
PAM -> DPP4
LAMC2 -> CD44
DEFB1 -> CCR6
COL14A1 -> CD44
HLA-A -> LILRA1
COL4A2 -> CD93
SLIT2 -> DCC
CNTN1 -> NOTCH2
ADAM10 -> IL6R
CXCL12 -> CD4
HLA-B -> LILRB2
ST6GAL1 -> CD22
CXCL12 -> ITGA4
CXADR -> JAML
CXCL12 -> CXCR4
HLA-B -> LILRA1
NECTIN2 -> CD226
ADAM10 -> TSPAN5
SPP1 -> PTGER4
CXCL12 -> DPP4
MUC1 -> SIGLEC9
ADAM10 -> TSPAN14
MMP7 -> CD44
CALM3 -> ESR1
PLAU -> PLAUR
NRG1 -> NETO2
ncMON
B
PL
cDC1
mDC
MAIT
Naive Th
Specificity rank
0.365
0.913
1.827
2.740
3.653
Magnitude rank
1.0
0.5

**SD Figure 11. TAL L-R analyses.** Most variable ligand-receptor interactions across TAL cell states from **a.** Fibroblasts to TAL **b.** TAL to Fibroblasts **c.** immune cells to TAL and **d.** from TAL to immune cells as found by LIANA+. The color bar indicates the magnitude and the size the specificity of the interaction (both a –log10 transformed). Non-specific interactions (specificity > 0.01) are in grey. Created in BioRender. Jain, S. (2025) https://BioRender.com/2dhbi4c.
